# Prey-specific toxins provide broad venom activity in cephalopods

**DOI:** 10.64898/2026.02.02.703354

**Authors:** Thomas Lund Koch, Ho Yan Yeung, Joshua P. Torres, Zachary Brandt, Seph Palomino, Iris Bea L. Ramiro, Zildjian G. Acyatan, Nicholas C. Schumann, Alina Anhnhi Vo, Kevin Chase, Ebbe Engholm, Knud J. Jensen, Lasse Holst Hansen, Randall T. Peterson, Michael J. Robertson, Amol Patwardhan, Katrine T. Schjoldager, Helena Safavi-Hemami

## Abstract

Cephalopods are among the ocean’s most sophisticated predators that use camouflage, complex behaviors, and venom to subdue a wide range of prey. However, the functional role of venom across diverse prey remains poorly understood, particularly whether cephalopods deploy venom to capture fish. Through comprehensive transcriptomic profiling of venom glands, we identify toxins with molecular signatures of prey-specific adaptation, including a previously unrecognized family of peptide toxins, octotensins, that evolved through convergent evolution to mimic the vertebrate hormone neurotensin. Functional assays and cryo-electron microscopy demonstrate that octotensins potently activate fish and human neurotensin receptor 1, engage this target in a near-identical manner to the chordate hormone, and induce acute hypotension in rodents. Together, our findings demonstrate that cephalopods achieve broad venom activity through phylum-specific toxins, including those targeting fish, revealing an evolutionary strategy by which generalist predators can capture phylogenetically diverse prey.

**One-Sentence Summary:** Cephalopod venom comprises prey-specific toxins, including neurotensin-mimicking peptides that target fish.

## Main Text

Predation is a complex trait that relies on genetic, morphological, physiological, and behavioral adaptations. Coleoid cephalopods (octopus, squid, and cuttlefish) are advanced predators that employ an intricate visual system, remarkable camouflage, behavioral flexibility, and venom to capture prey (*1-3*). As generalist predators, cephalopods feed on diverse phyla, including arthropods (crustaceans), mollusks (bivalves and gastropods), and chordates (teleost fish) (Note S1, Movie S1-S2). Venom is essential in several cephalopod hunting strategies (*4*): octopuses drill holes through the protective shell of mollusks and inject venom for paralysis (*5*). Similarly, for crab hunting, venom can be injected either by drilling through the exoskeleton or via eye puncture, resulting in paralysis and death (*6*). However, the role of venom in fish predation remains unknown.

Many venomous predators specialize on particular prey and tune their venom composition to specific targets (11-15). Generalist predators, such as cephalopods, must on the other hand produce venom effective across phylogenetically diverse organisms. How generalists achieve broad venom potency remains largely unknown, but three strategies are likely: the presence of promiscuous toxins, multiple prey-specific toxins, or both. Cephalopod venoms are primarily protein-based, and most of the half dozen characterized toxins are enzymes, such as proteases and lipidases, which likely act broadly on phylogenetically diverse prey (*7-10*). The presence of prey-specific toxins in cephalopods remains unexamined, yet resolving this question is critical for linking venom evolution to ecological and behavioral strategies and for exploring possible biomedical uses.

To investigate such prey-specific toxins, we focus on “doppelgänger peptides”, venom components that mimic the hormones and neuropeptides of prey. Because their molecular targets are often restricted to particular phyla, we hypothesize that identifying such toxins will offer insight into whether cephalopods employ toxins tailored to specific prey lineages, including fish.

### Cephalopod toxins can be classified into multiple toxin superfamilies

Cephalopod venom is produced by two paired glands: the anterior (ASG) and posterior salivary glands (PSG) (*7*) (Fig. 1A). To identify potential signatures of prey-specific toxins, we first surveyed all highly expressed transcripts encoding secreted proteins of five octopodiformes and two decapodiformes ASG and PSG, as these likely represent toxins (Table S1). We identified 822 putative toxin transcripts belonging to 44 superfamilies (SFs) (File S1, Fig. S1).

**Fig. 1.**
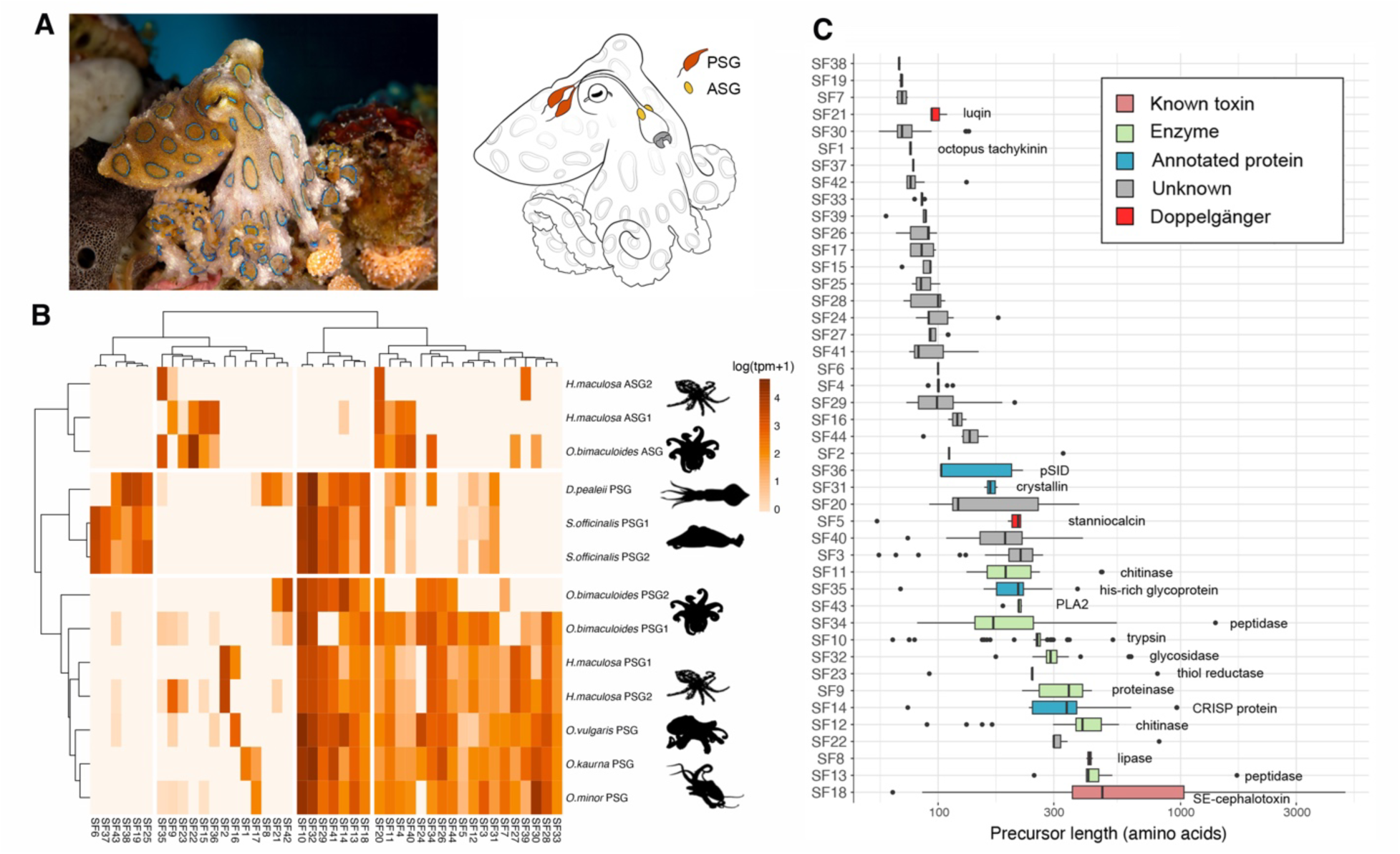
Cephalopod venom system and expression profiles. (A) Cephalopods produce venom in paired anterior (ASG) and posterior salivary glands (PSG), illustrated here for the blue-ringed octopus, *Hapalochlaena maculosa.* (B) Heatmap of highly expressed secreted proteins from ASG and PSG across five octopods (*Octopus vulgaris, Octopus bimaculoides, Octopus kaurna, Octopus minor,* and *H. maculosa*) and two decapods (*Doryteuthis pealeii*, *Sepia officinalis*). Toxins were grouped into superfamilies (SFs) based on sequence similarity. (C) Size distribution and annotations of toxin superfamilies shown as boxplot with outliers, minimum, 25^th^ percentile, median, 75^th^ percentile and maximum. Image: David Benz. Illustration by ilusea Studio and PhyloPic.

Hierarchical clustering of the tissue expressions places the ASGs as an outgroup of the PSGs with further subdivision of the octopod and decapod PSGs (Fig. 1B). While certain SFs were ubiquitously expressed in the PSGs, others were restricted to specific samples. We observed intraspecies variation in the expression profiles of the ASG and PSG of the blue-ringed octopus, *Hapalochlaena maculosa*, as well as the PSG of the California two-spot octopus, *Octopus bimaculoides*, consistent with previous observations (*2, 11*).

Several of the identified toxins were previously isolated and characterized from venom. For instance, SE-cephalotoxin from the golden cuttlefish, *Sepia esculenta* (*12*), belongs to SF18 and octopus tachykinins from the common octopus, *Octopus vulgaris* (*11, 13*), belong to SF1. Many of the secreted proteins could be annotated as enzymes. This included lipases (SF8 and SF43), proteinases (SF9, SF10, SF13, and SF34), chitinases (SF11 and SF12), and thiol reductases (SF23). Additionally, we identified 27 SFs of unknown function. These encode short proteins (median size < 300 amino acids), many of which have no homologs outside of Cephalopoda (Fig. 1C). Three of these (SF14, SF26, SF41) have previously been suggested to be cephalotoxins (*8, 10*), but their functional roles have not been assessed.

### Sequence homology and motif-based searches identify cephalopod doppelgänger toxins

Following the identification and classification of putative cephalopod toxins, we next interrogated the presence of potential doppelgänger peptides. Neuropeptides and peptide hormones (collectively referred to as neuropeptides here) are key regulators of animal physiology and behavior (*14*). Their critical roles make neuropeptide signaling systems particularly susceptible to exploitation, and many venomous organisms have evolved toxins that mimic the neuropeptides of their target species (*15, 16*). These doppelgänger toxins manipulate endogenous signaling pathways and are increasingly recognized as valuable tools for biomedical research and as potential drug leads. Detecting doppelgänger toxins, however, is challenging due to their small size and limited sequence similarity to endogenous neuropeptides (*15*).

To overcome this challenge, we combined BLASTp, profile hidden Markov models (pHMM), and a custom program, Fuzreg, which constructs modified regular expressions from alignments and searches for sequences within a given Levenshtein distance. Specifically, we generated pHMM and Fuzreg models for 54 physiologically important chordate, 28 crustacean, and 33 molluscan neuropeptides, which we then used as queries against all putative cephalopod toxin sequences.

Using these approaches, we identified doppelgänger toxins mimicking stanniocalcin (SF5), luqin (SF21), myosuppressin (SF1), tachykinins (SF1), and neurotensin peptides (neurotensin and neuromedin N) (SF1) (Fig. 2, Fig. S2). While BLASTp successfully detected stanniocalcin and luqin, only pHMM and Fuzreg were able to identify myosuppressin-, tachykinin-, and neurotensin-like doppelgänger, with Fuzreg showing greater sensitivity than pHMMs (Fig. 2B). Some additional hits were detected by Fuzreg, but these lack key neuropeptide characteristics and were therefore considered likely false positives, pending further validation (*e.g.*, Fig. S3).

**Fig. 2.**
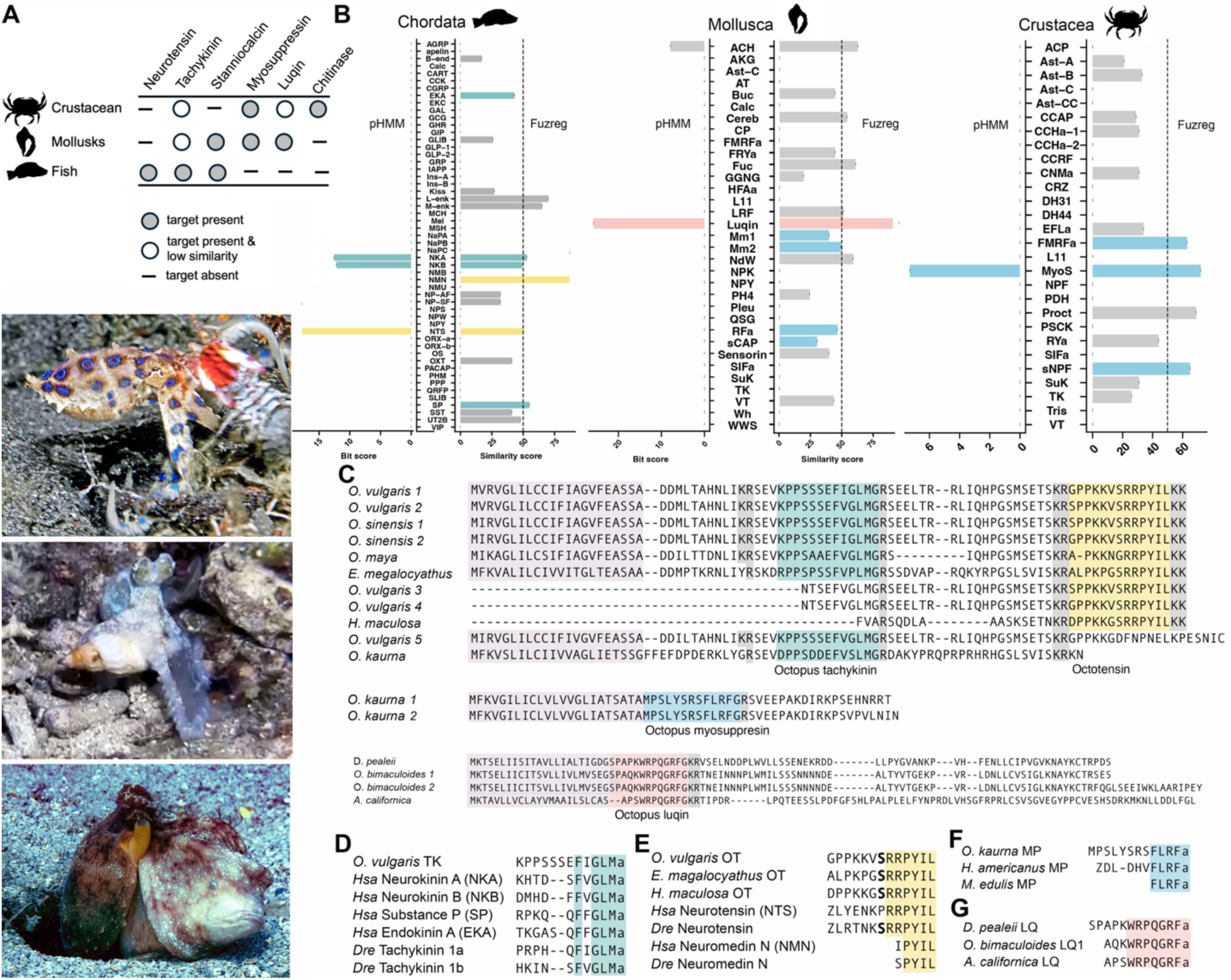
Prey-phyla specific doppelgänger toxins expressed in cephalopod venom glands. (A) Presence (gray filled circles) and absence (–) of prey-specific toxins. White filled circles indicated presence of the target but with low similarity compared to other prey phyla. Images of octopuses feeding on diverse prey (top to bottom: crab, mollusk, fish). Image credit: Ned and Anna DeLoach, Roger Hanlon. (B) Identified doppelgänger of chordate, molluscan, and crustacean neuropeptides. Left: bit scores from profile hidden Markov models (pHMM). Right: similarity scores from Fuzreg. Tachykinins (dark green), neurotensins (yellow), luqins (light red), myosuppressins (blue), and putative false positives (gray). See Note 2 for peptide abbreviations. (C) Alignments of doppelgänger-containing precursors from venom gland transcriptomes, color-coded as in panel B. Gray = cleavage sites; light purple = signal peptides. (D–G) Alignments of predicted mature doppelgänger toxins from *O. vulgaris*, *O. kaurna*, and *D. pealeii* with representative prey neuropeptides. (D) Tachykinin mimics and tachykinins from human (*Hsa*) and zebrafish (*Dre*). (E) Neurotensin mimics and *Hsa* and *Dre* neurotensins. (F) Myosuppressin mimics and myosuppressins. (G) Luqin mimics and luqins. ‘a’ = amidation; ‘Z’ = pyroglutamate; bold S = predicted glycosylation site.

Because the peptide systems being mimicked by these doppelgänger toxins are largely restricted to specific prey lineages (Fig. 2A), their identification provides direct insight into cephalopod predatory strategies. Stanniocalcins are large glycosylated hormones involved in calcium and phosphate homeostasis, and found in most metazoans but are absent from arthropods including crustaceans (*17*). Luqins are cardio-excitatory neuropeptides that are present in protostome invertebrates, including crustaceans and molluscs, but are absent in chordates including fish (*18*). Myosuppressin and other FLRFamide-like peptides occur only in protostomes and are known to modulate muscle contraction and heart rate in crustaceans (*19*). In contrast, neurotensins (NTS and NMN) are restricted to chordates, strongly suggesting that cephalopod neurotensin doppelgänger, hereafter referred to as octotensins, evolved specifically to target fish prey (Fig. 2A).

### SF1 is an unusual venom superfamily that harbors multiple doppelgänger toxins

Unexpectedly, we observed that octotensins are encoded on the same protein precursors as tachykinin-like toxins within SF1 (Fig. 2C). Octopus tachykinins are venom peptides that share sequence similarity with tachykinins (*11, 20*), metazoan neuropeptides that regulate smooth muscle contraction, vascular tone, and neurotransmission through activation of the neurokinin 1-3 receptors (*21*). Although octopus tachykinins have previously been described in some species, the presence of NTS-like toxins, octotensins, has not been reported.

Using the retrieved SF1 precursors as queries, we performed additional homology searches to uncover the full diversity of SF1 toxins, leading to the identification of 31 transcripts from Octopoda. Graph-based sequence clustering revealed distinct subgroups within SF1, including ten sequences that encode both octotensins and tachykinins (Fig. S4). These precursors contain conserved (di)basic cleavage sites flanking the predicted peptides, enabling proteolytic release. Predictions using the DeepPeptide tool support the release of mature octotensins with high confidence (*22*) (Fig. S5). Evolutionary trace analysis further shows that the six most C-terminal amino acids of octotensins (RRPYIL) are highly conserved, mirroring chordate NTS (Fig. 2C-E, Fig. S6-7). Unlike NTS, tachykinins are not restricted to chordates, however, we observe that the tachykinin doppelgänger display far higher similarity to chordate tachykinins (EKA, NKA, NKB, and SP) (Fig. 2D) than to any invertebrate tachykinins (Fig. S8, File S2) and are capable of potently activating the human neurokinin 1 and 3 receptors when compared to the endogenous ligand Substance P (Fig. S9). These observations strongly suggest that both tachykinin and octotensin doppelgänger are targeted at vertebrate prey, with fish being the only known vertebrate that cephalopods prey on. Interestingly, two precursors from the southern sand octopus, *Octopus kaurna,* encode neither tachykinins nor octotensins, but instead the myosuppressin-like peptides, which are unique, cardioactive neuropeptides in protostomes (Fig. 2C, F).

Thus, SF1 represents a rare toxin superfamily that has diversified to mimic multiple distinct neuropeptides that share no evolutionary relationship.

### Superfamily-1 toxins are evolutionarily unrelated to the neuropeptides they mimic

To understand the evolutionary origin of these diverse doppelgänger toxins within SF1, we investigated whether they arose from the recruitment of endogenous neuropeptides or from *de novo* innovation. Most doppelgänger toxins evolve from the recruiting of the venomous animal’s own neuropeptides into the venom system (*23-25*). However, in rare cases, they can evolve *de novo* from unrelated genetic material (*15, 26*). Gene structural analyses reveal that the SF1 superfamily has a unique organization compared to the endogenous cephalopod neuropeptides it mimics (tachykinin, and myosuppressin) and could have derived from (Fig 3A). This strongly supports their convergent, *de novo* origin. Additionally, as neurotensin is a chordate specific neuropeptide (*27*), direct recruitment from an endogenous cephalopod neuropeptide is not possible. Similarly, horizontal gene transfer from a vertebrate is also extremely unlikely as vertebrate neurotensin and octotensins have distinct gene structures (Fig 3A).

**Fig. 3.**
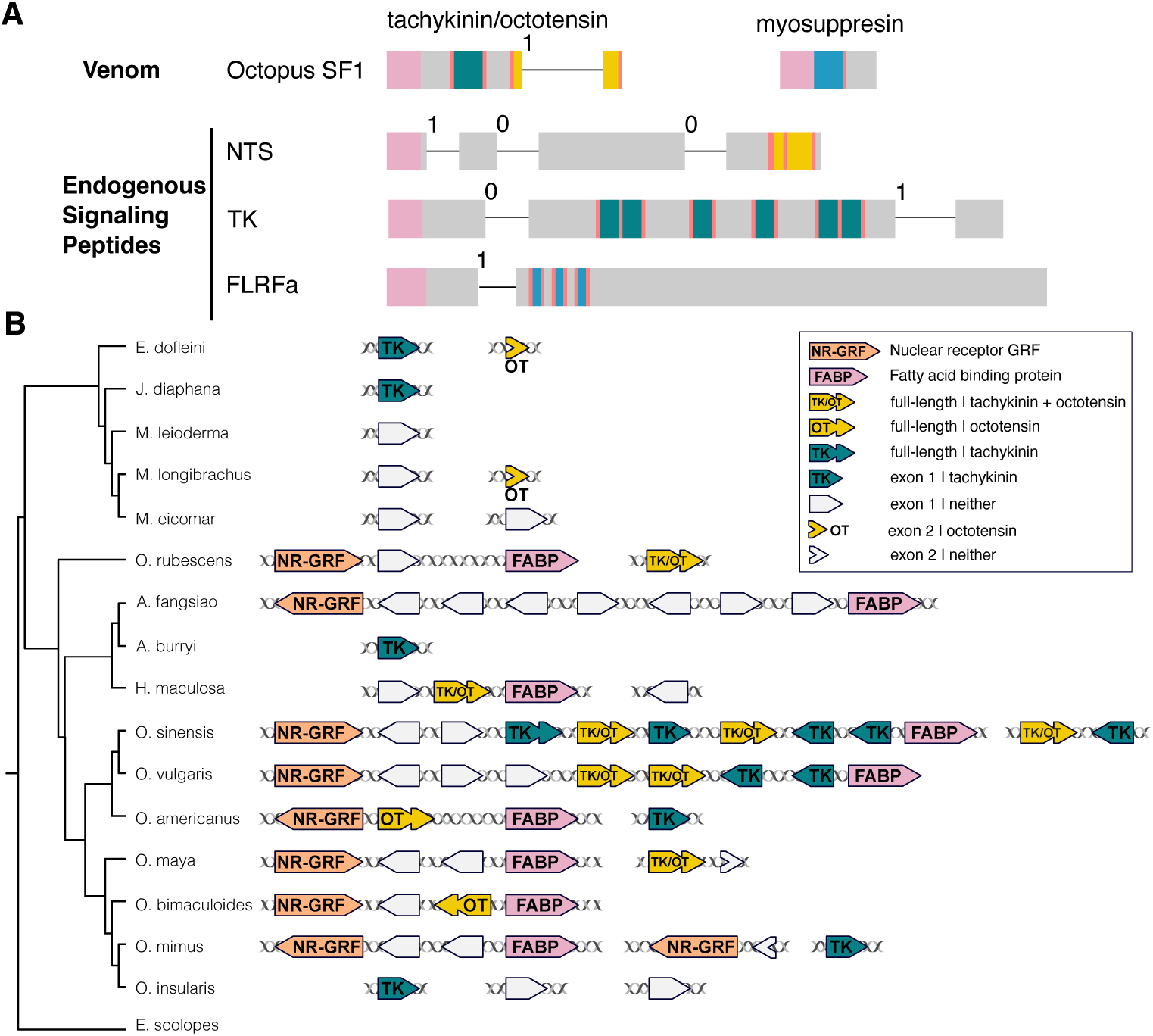
Superfamily 1 encodes multiple convergently evolved doppelgänger toxins and is unique to octopodiformes. (A) Gene structure comparisons show that SF1 genes have a distinct organization relative to the neuropeptides they mimic: NTS (*D. rerio* NTS), TK (*O. sinensis* tachykinin), and FLRFa (*O. sinensis* FLRFamide precursor). Superscripts indicate intron positions and phases. (B) Left: Phylogenetic reconstruction of Octopoda using published genomes. Right: Synteny of genomic regions surrounding SF1 genes, colored as in the legend. SF1 genes are exclusive to octopodiformes, occur in tandem arrays, and are consistently flanked by NR-GRF and FABP genes. See Table S2 for genomes analyzed.

Synteny analyses further indicate that SF1 genes are confined to octopodiformes, where local gene duplications have greatly expanded this family (Fig. 3B), as has been observed for other cephalopod protein families (*28*). We recovered 1-10 SF1 genes across octopodiforme genomes (Table S2), with particularly large expansions in the two sister species *O. vulgaris* and *Octopus sinensis*, and the relative, *Amphioctopus fangsiao*. These genes are consistently flanked by fatty acid binding protein and nuclear receptor GRF on opposite sides. As coleoid cephalopods have undergone extensive chromosomal rearrangements (*29*), large gene synteny comparisons to species outside of this lineage is not feasible. Extensive searches in other cephalopod genomes failed to identify related sequences, suggesting that SF1 is a unique innovation of octopodiformes.

Collectively, our findings demonstrate that cephalopod venoms contain doppelgänger toxins with clear signatures of prey neuropeptides (Fig. 2A). Because several of these neuropeptides are restricted to one or a few prey lineages, for example, NTS in chordates and luqin in protostomes, our results support the hypothesis that cephalopods use their venom for all three prey phyla and have evolved specific toxins to target the particular prey they encounter.

The unexpected discovery of NTS-like sequences in octopus venom, prompted us to further investigate the biochemistry, pharmacology, and biological functions of octotensins.

### Octotensins are potent agonists of fish and human neurotensin receptor 1 (NTSR1)

Many bioactive peptides require post-translational modifications for activity (*30-32*). Because octotensins were not directly isolated from venom, we analyzed their sequences for putative modification sites and found conserved serine residues predicted to undergo *O*-glycosylation using neural network-based *O*-glycan prediction software (Fig. S10). *O*-glycosylations have also been described for several other cephalopod toxins (*12, 33*) and the enzymes catalyzing this modification are expressed in the posterior salivary glands (Table S3). Together, these observations suggest that native octotensins may be glycosylated. Thus, we synthesized both the glycosylated and non-glycosylated isoforms for further study.

In humans, NTS is the endogenous ligand for two class A G protein-coupled receptors (GPCRs): neurotensin receptor 1 (NTSR1) and neurotensin receptor 2 (NTSR2). Teleost fish, the proposed target of octotensins, have been suggested to encode only a single NTS receptor, orthologous to human NTSR1 (*34*). Phylogenetic and synteny analyses support that NTSR2 has been lost in this and several other lineages (Fig. S11-12). To test whether octotensins can activate the fish receptor, we performed the inositol monophosphate One (IP-One) Gq assay on zebrafish (*Danio rerio*) NTSR1 (Dre-ntsr1), a canonical Gα_q/11_-coupled receptor. Our results demonstrate that octotensins from three different species, OT-Eme, OT-Hma, and OT-Ovu1 are potent agonists of Dre-ntsr1 with potencies comparable to human NTS (EC_50_: OT-Eme 90 pM, OT-Hma: 150 pM, OT-Ovu1: 174 pM, human NTS: 325 pM). Furthermore, we observed no difference between an *O*-glycosylated OT-Ovu1 peptide, OT-Ovu1g (containing Ser7-Gal-GalNAc), when compared to the non-glycosylated or a partially glycosylated, OT-Ovu1pg (containing Ser7-GalNAc), isoform in these assays (Fig. 4A, Table S4). Similar results at fish NTSR1 were obtained using a modified β-arrestin recruitment assay (PRESTO-Tango (*35*), Fig. S13, Table S5). Furthermore, all peptides also robustly activated human NTSR1-mediated IP-One accumulation with comparable potencies to human NTS (Fig. 4B, Table S6). Human NTSR1 shares 60% sequence identity with the zebrafish receptor.

**Fig. 4.**
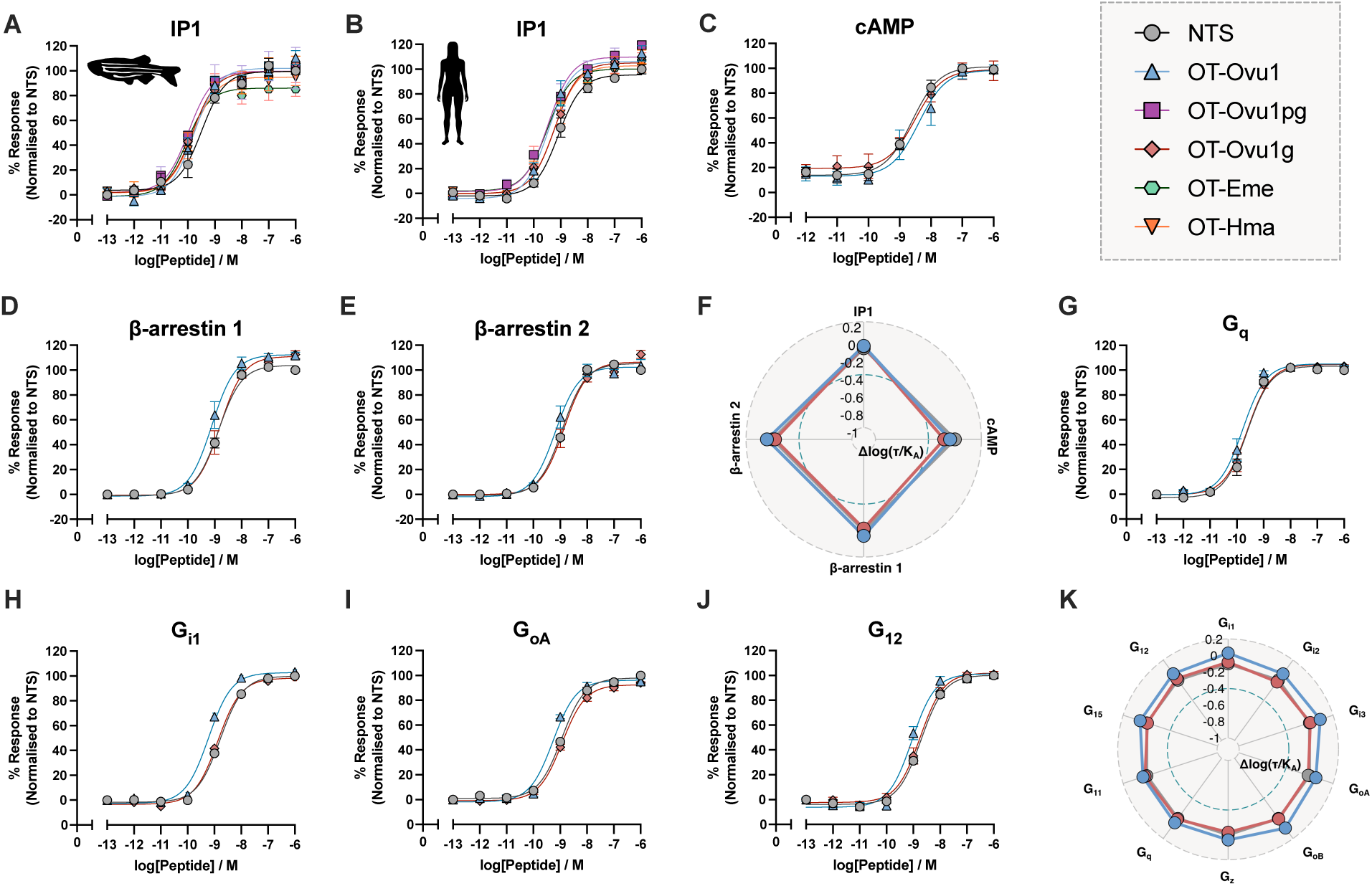
Octotensins mimic the pharmacology of chordate neurotensin at fish and human NTSR1. (A- B) Dose-response curves of NTS, three OT-Ovu1 isoforms from *O. vulgaris* (OT-Ovu1, OT-Ovu1g, and OT-Ovu1pg), and octotensins OT-Hma and OT-Eme from *O. maculosa* and *E. megalocyathus*, respectively, showing inositol monophosphate (IP) responses in cells transiently expressing zebrafish or human NTSR1. (C - E) cAMP signaling and β-arrestin 1/2 recruitment induced by NTS, OT-Ovu1, and OT-Ovu1g. (G - J) G-protein dissociation assays (G_q_, G_i1_, G_oA_ G_12_). All data represent means ± S.E.M. of 3 biological replicates with technical duplicates. (F, K) Downstream signaling and G protein bias plot (Δlog(τ/K_A_)) for OT-Ovu1 and OT-Ovu1g using NTS as physiological reference.

In addition to canonical Gα_q/11-_coupling, NTSR1 can engage other Gα proteins, such as Gα_i/o_ (*36*), and undergo receptor desensitization via β-arrestin 1/2 recruitment to prevent overstimulation. To comprehensively characterize signaling, we evaluated both non-glycosylated OT-Ovu1 and glycosylated OT-Ovu1g at human NTSR1 using a series of downstream signaling assays. Measurements of cAMP signaling and β-arrestin 1/2 recruitment revealed equipotent responses for both isoforms relative to human NTS (Fig. 4C-F, Table S7-8). Furthermore, NTS and both OT-Ovu1 and OT-Ovu1g potently induced dissociation of Gα_q/11_, Gα_i/o_, and G_12_ proteins (Fig. 4G-K, Fig. S14-15, Table S9-10), whereas none of the peptides triggered Gα_s_ dissociation (Fig. S14).

Together, these results demonstrate that octotensins replicate the pharmacological profile of chordate neurotensins, including their ability to signal through multiple pathways. Furthermore, glycosylation does not alter the potency or efficacy at either the fish or human NTSR1.

### Cryo-EM reveals that OT-Ovu1g structurally mimics chordate NTS at human NTSR1

To determine whether the pharmacological mimicry of octotensins is mirrored structurally, we obtained cryo-EM reconstructions of OT-Ovu1g bound to human NTSR1 in complex with the G_i3_ heterotrimer and scFv16. Consistent with previously reported NTSR1/Gi structures, we observed two receptor orientations relative to the G protein, related by a ∼45° rotation, a non-canonical conformation at 2.1 Å resolution and a canonical at 2.2 Å resolution (Fig. 5A-B, Fig. S16-18, (*37*). Both conformations exhibited highly similar orthosteric binding pockets (Fig. S16). Notably, despite differences in the N-termini of OT-Ovu1 and human NTS, the C-terminal RRPYIL motif of OT-Ovu1 which is conserved in all octotensins identified in this study adopts a pose largely superimposable with that of human NTS bound to NTSR1 (Fig. 5B-C) (*38, 39*). Mutations of receptor residues interacting with this motif (R323^6.55^, F326^6.58^, W334^ECL3^, and Y339^7.28^) either caused 10-100-fold reductions in OT-Ovu1 potency, or were deleterious for receptor signaling (R322^6.54^, Y342^7.31^ and Y346^7.35^) (Fig. 5D-E), demonstrating that the peptide mimicry relies on this motif for effective receptor engagement and signaling.

**Fig. 5.**
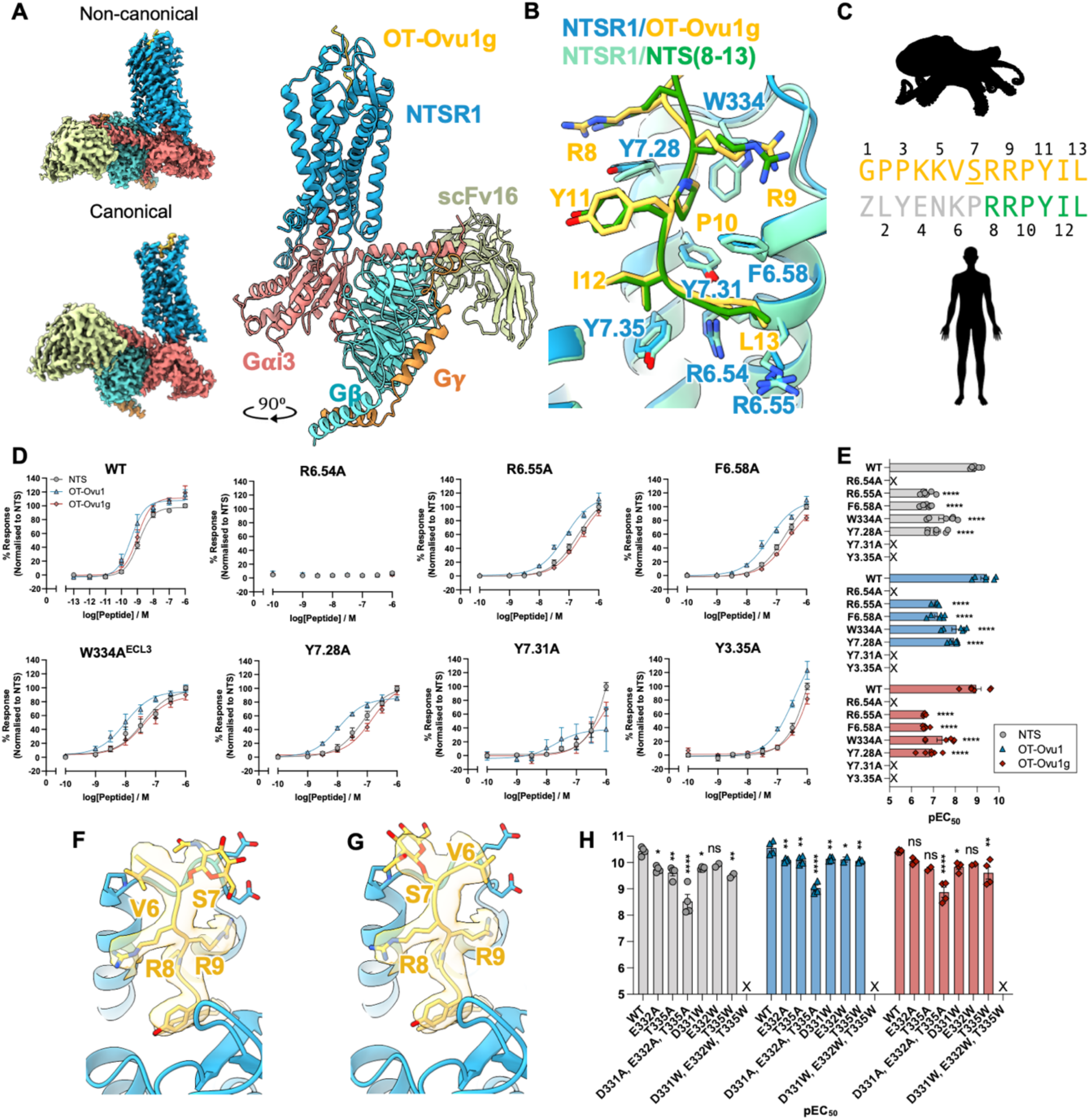
Octotensin Ovu1g engages human NTSR1 in a near-identical manner to human NTS. (A) Cryo-EM maps of the OT-Ovu1g NTSR1 Gi3 complex in non-canonical and canonical conformations (left) and cryo-EM structure of the non-canonical conformation. NTSR1 (blue), OT-Ovu1 (yellow), Gα_i3_ (rose), Gβ (cyan), Gγ (orange) and scFv16 (beige yellow). (B) Overlay of OT-Ovu1g–NTSR1 (yellow/blue) with human NTS(8-13)–NTSR1 (green/pale green; PDB: 4grv). Core OT-Ovu1g motifs (yellow) and receptor contact residues (blue, Ballesteros-Weinstein numbering) are highlighted. (C) Alignment of OT-Ovu1g with human NTS. Ser7 glycosylation underlined. (D-E) NTSR1 core mutations reduce IP accumulation mediated by NTS (gray), OT-Ovu1 (blue) and OT-Ovu1g (red). (D) Dose-response curves and (E) pEC_50_ values. X: non-determinable potency. Error bars show mean ± S.E.M. (n=3). **** p < 0.0001 vs. WT (one-way ANOVA, Dunnett’s test). (F-G) Unsharpened cryo-EM map around the flexible Ser7 glycosylation site and Val6 of OT-Ovu1g. (H) pEC_50_ values for NTSR1 WT and mutants potentially forming contacts with the glycosylation site. *, p > 0.05, **, p > 0.01, ***, p > 0.001, ****, p < 0.0001 and ns, non-statistically significant.

We also resolved densities around Ser7, the glycosylation site, and Val6, though resolution was insufficient to definitively model this portion of the peptide (Fig. 5F-G). Generating the two possible poses of the peptide and the Gal-GalNAc moiety placed the sugars near the ECL3, either adjacent to D331, E332, and T335 or near P336 depending on the model. However, mutations of these residues did not significantly alter receptor activation, further indicating that glycosylation has little effect on NTSR1 signaling (Fig. 5H, Fig. S19).

Collectively, these results show that octotensins mimic NTS not only pharmacologically but also structurally, suggesting that these peptides have evolved under selective pressure to reproduce the core receptor interactions of vertebrate neurotensin.

### Octotensins induce acute hypotension and increase heart rate in rodents

Having established that octotensins evolved convergently to mimic both the pharmacology and structure of vertebrate neurotensin, we next sought to define their biological activity. In humans, neurotensin regulates a wide range of peripheral and central processes, including cardiovascular function, gut motility, thermoregulation, and pain (*40-43*). Neurotensin was named for its neuronal origin and its ability to alter vascular tension, producing acute hypotension when administered peripherally (*43*). Based on these important cardiovascular effects of neurotensin, we tested whether octotensins disrupt cardiovascular function *in vivo*. Intravenous administration of 20 nmol/kg OT-Ovu1 and OT-Ovu1g induced a rapid and sustained decrease in diastolic, systolic, and mean arterial blood pressure compared to vehicle controls (Fig. 6A-C), accompanied by an increase in heart rate (Fig. 6D). Although maximal changes in blood pressure and heart rate did not differ significantly between the glycosylated and non-glycosylated isoforms, the glycosylated peptide appeared more potent, as observed by visible cyanosis in three of four treated animals and lethality in one case. Together, these findings suggest that octotensins facilitate prey capture by inducing acute hypotension, leading to systemic hypoperfusion and compromised organ function.

**Fig. 6:**
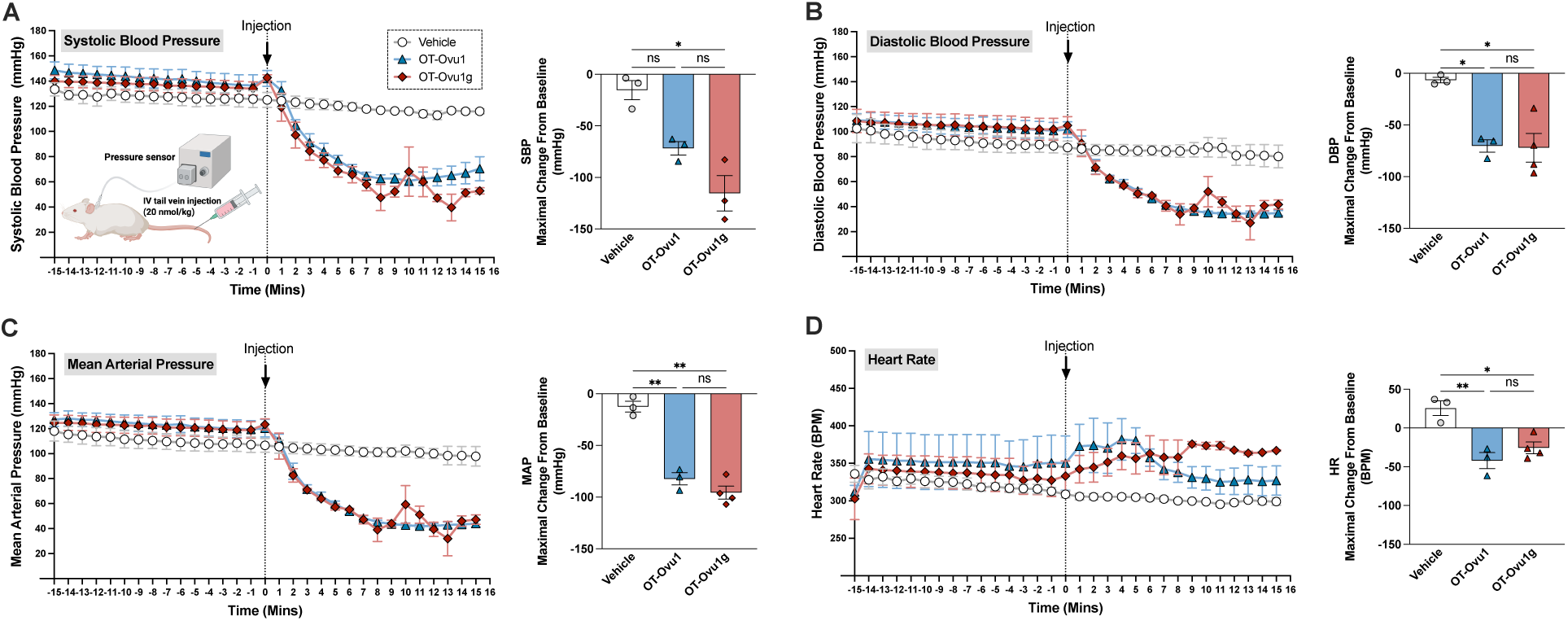
Octotensins significantly reduce blood pressure in anesthetized rats. <u>Blood pressure</u> and heart rate were measured in male, Sprague Dawley rats (8-10 weeks) via direct cannulation of carotid artery with fluid-filled catheter connected to pressure transducer under anesthesia. Continuous measurements of (A) systolic, (B) diastolic, (C) mean arterial blood pressure (in mmHg) and (D) heart rate (in BPM) before and after 20 nmol/kg of OT-Ovu1 (n=3), OT-Ovu1g (n=4), and vehicle (n=3) bolus IV injections via tail vein. Maximal change from baseline of blood pressure and heart rate are depicted in scattered plots. The data are represented as mean ± S.E.M. Two-way ANOVA with Tukey’s post hoc for multiple comparisons was performed using to denote significance (* p < 0.05, ** p < 0.01, ns, non-statistically significant). Illustration in (A) was prepared using Biorender.

Notably, the octopus tachykinin eledoisin, isolated from the musky octopus *Eledone muschata*, has also been shown to induce pronounced hypotension in rats (*44*), consistent with the well-established hypotensive activity of tachykinin family peptides, including substance P (*45, 46*). Because octotensins are encoded on the same protein precursors as octopus tachykinins (Fig. 2), and both peptide families potently activate vertebrate receptors (Fig. S9, Fig. 4) and induce hypotension (Fig. 6 (*44*)), we propose that these doppelgänger peptides act in concert to disrupt cardiovascular function in fish prey. To our knowledge, this is the first example in which a single venom precursor encodes two peptide toxins that mimic distinct, evolutionarily unrelated neuropeptides to act on the same physiological circuit.

## General discussion

Many venomous animals are dietary specialists and only prey on specific species. This allows for targeted molecular evolution of their toxins to obtain high potency and efficacy (*47-51*). In contrast, generalist predators face a more complex evolutionary challenge: generating venoms that maintain effectiveness across diverse prey lineages. Cephalopods exemplify this opportunistic generalism, and their remarkable behavioral flexibility, advanced nervous systems, and capacity for sophisticated learning make them especially intriguing models for studying venom evolution and function. By using doppelgänger toxins as a tool to probe biotic interactions, we reveal that, in addition to broad-spectrum enzymatic toxins, cephalopods have evolved lineage specific toxins that target particular prey phyla.

The evolutionary strategy of combining prey-specific toxins with broadly acting ones carries significance well beyond ecology and evolution. Venoms are treasure troves of bioactive compounds, many of which have already provided biomedical tools and drug leads (*52*). Venoms from animals that prey on vertebrates are of particular interest for biomedical research, as their molecular targets are often conserved across vertebrate species, including humans. Although cephalopods are known to capture fish, the contribution of venom to this process had not been investigated. Here, we provide evidence that octopus venom is used for fish prey capture. The potent activity of octotensins (and tachykinins) at vertebrate receptors suggests that cephalopod venoms contain additional vertebrate-targeting components that remain to be characterized. We note the expression of many peptides and proteins in cephalopod venom glands that have no known function. While we were unable to demonstrate fish-targeting toxins in non-octopodiform cephalopods, these species likely also express such molecules that have not yet been detected, in part because they lack recognizable sequence features. Taken together, our identification of doppelgänger toxins that mimic neurotensin – octotensins – underscores the potential of cephalopod venoms as sources of human-relevant bioactive compounds and provides a foundation for further investigation into their possible applications in drug discovery.

We demonstrate that octotensins evolved *de novo* as doppelgänger of a vertebrate hormone. This molecular mimicry underlies their potent activity at vertebrate receptors, with cryo-EM revealing that octotensins engage NTSR1 through a binding mode that is nearly indistinguishable from that of the endogenous chordate peptide. Functionally, octotensins induce acute hypotension accompanied by an increase in heart rate, strongly suggesting that these peptides facilitate prey capture by disrupting cardiovascular function. Targeting blood pressure is a common strategy among venomous animals and has evolved independently through distinct molecular mechanisms, including bradykinin-potentiating and natriuretic peptides from snakes (*53*), bradykinin-like peptides from wasps and frogs (*26, 54, 55*), and adrenomedullin-like peptides from ticks (*56*). The discovery of snake venom bradykinin-potentiating peptides directly led to the development of the angiotensin-converting enzyme inhibitor captopril (tradename Capoten) (*57*), now widely used to treat hypertension.

Although toxins that induce hypotension through NTSR1 are unlikely to serve as viable antihypertensive drug leads due to the broad peripheral effects associated with NTSR1 agonism, neurotensin analogs have attracted considerable interest as therapeutics for central pain. This includes contulakin-G, a neurotensin-like peptide from the venom of the fish-hunting cone snail *Conus geographus*, that produces potent analgesia in mammalian pain models and demonstrated efficacy in a phase 1 clinical trial (*58*). Interestingly, an *O*-glycosylation at Thr10 (Gal-GalNAc) is essential for the *in vivo* analgesic activity of contulakin-G (*59, 60*). Consistent with this, we find that, in a postoperative paw-incision model with intrathecal administration, OT-Ovu1g but not the non-glycosylated isoform, OT-Ovu1, significantly reduced post-incisional mechanical hypersensitivity (Fig. S20). However, this effect was observed only at very high doses (200 nmol) that also caused transient locomotor impairment, limiting the therapeutic potential of OT-Ovu1g as an analgesic. Because only the glycosylated isoform, OT-Ovu1g, provided analgesia and also showed a trend toward more pronounced hypotensive effects than the non-glycosylated isoform, despite indistinguishable NTSR1 engagement and activity *in vitro*, glycosylation likely enhances *in vivo* stability or enables interactions not recapitulated *in vitro*. Similar roles have been proposed for this modification in contulakin-G (*61*), although the underlying mechanisms remain unresolved.

Contulakin-G is the only other known neurotensin-like peptide identified from venom. Although both contulakin-G and octotensins originate from molluscan venoms, they do not share evolutionary ancestry with each other or with chordate neurotensin. Our data show that octotensins evolved *de novo* within the SF1 venom gene family (Fig. 3), whereas contulakin-G arose from the cone snail C-conotoxin superfamily in a single species, *Conus geographus.* The C-conotoxin gene family itself originated through duplication and diversification of an endogenous somatostatin-related neuropeptide gene (*23, 62*). Sequence alignments show no similarity among these three precursors beyond the neurotensin motif (Fig. S21).

The independent evolution of doppelgänger peptides of neurotensin in both octopuses and the fish hunter *Conus geographus* reveals an apparent vulnerability of this neuropeptide signaling system in fish prey. More broadly, these findings provide another fascinating example of *de novo* evolution of peptide toxins that bear no evolutionary relationship to the neuropeptides they mimic. In octopuses, this phenomenon is particularly pronounced: two distinct neuropeptide mimics are encoded on a single toxin precursor, and an additional toxin from the same superfamily (SF1) encodes a third doppelgänger peptide that mimics myosuppressin in *Octopus kaurna*. Together, these findings illustrate the extraordinary plasticity of venom genes. While such convergent molecular mimicry may seem improbable, we recently identified an analogous evolutionary pattern in bradykinin-like peptides from hymenoptera venoms and frog skin secretions (*26*), suggesting that the *de novo* evolution of neuropeptide mimics may be more widespread than previously recognized.

Our study has some limitations that warrant further investigation. First, our discoveries are based on transcriptomic and genomic data. We did not confirm the presence of octotensins at the protein level. Future studies are needed to validate the mature peptides and their glycosylation status. Second, given the wide range of physiological effects reported for neurotensin, octotensins may target other circuits beyond the cardiovascular system. Finally, given the functional activity of octotensins as hypotensive agents, we propose that these peptides are primarily used for capturing rather than deterring fish. Future field observation studies are needed to investigate this hypothesis.

Beyond these limitations, the discovery of octotensins demonstrates how doppelgänger peptides provide a window into predator-prey interactions. Determining prey specificity of toxins typically relies on behavioral or toxicity studies, but these are limited by ethical concerns and the need for field observations or specialized facilities, making them impractical for rare or difficult-to-keep species. As sequencing data increasingly outpace ecological and toxicological studies, we show that a sequence-based approach can provide valuable insights into toxin identity, pharmacology, and likely ecological function. The identification of the SF1 toxin superfamily, which encodes various peptides mimicking prey neuropeptides, illustrates this principle. Computational tools that detect receptor-targeting motifs and predict functionally relevant post-translational modifications further enhance this strategy, and ongoing advances in these methods will continue to expand its power. More broadly, the approach outlined here can be extended to other venomous animals, as well as to parasites and pathogenic microbes that use molecular mimicry to manipulate another organism’s physiology and behavior.

## Acknowledgments

We would like to thank Dr. Celeste M. Hackney and Dr. Aymeric Rogalski for their helpful feedback and discussion on this paper. We thank Dr. Gaya P. Yadav for cryoEM data collection at the Laboratory for Biomolecular Structure and Dynamics (LBSD) of Texas A&M University. The LBSD is supported, in part, by the Department of Biochemistry & Biophysics, AgriLife, and the Texas A&M University.

## Author contributions

Conceptualization: TLK, HYY, JPT, HSH

Methodology: TLK, HYY, JPT, ZB, SP, LHH, NS, KC, AAV, ZA, MJR, KTS, HSH

Investigation: TLK, HYY, JPT, IBLR, SP, LHH, EE, ZB

Visualization: TLK, HYY, SP, MJR, HSH

Funding acquisition: TLK, KJJ, KTS, MJR, AP, RTP, HSH

Project administration: HSH

Supervision: KJJ, RTP, MJR, AP, KTS, HSH

Writing – original draft: TLK, HYY, HSH

Writing – review & editing: TLK, HYY, JPT, IRLR, ZB, KC, EE, KJJ, NS, ZA, LHH, SP, RTP, MJR, AP, KTS, HSH

## Data and materials availability

Newly generated code is available at https://github.com/thomaslundkoch/fuzreg. All data are available in the main text, the supplementary materials, or references in those.

## Supplementary Materials

Materials and Methods

Supplementary Text

Figs. S1 to S21

Tables S1 to S11

References 63-104

Data S1 to S4

Movies S1-S2

## Supplementary Materials

### Materials and Methods

#### Genomic and transcriptomic data

RNAseq data from anterior and posterior salivary glands were downloaded from NCBI (Table S1). All genomic data and annotations were downloaded from NCBI, except for the genome of *A. fangsiao*, which is available for download from figshare (Table S8). We used fqtrim (*63*) to trim adapters of de-multiplexed raw reads, and prinseq-lite (*64*) for quality trimming and filtering. BBnorm ecctool was used for error correction. Filtered reads were assembled using Trinity v2.2.1 (*65*) using k-mer length of 31, and mimimum k-mer coverage of 10. Expression (transcripts per million/tpm) was calculated using Trinity RSEM plugin align_and_estimate_abundance (*66*).

#### Hierarchical clustering of toxin candidates

We extracted secreted proteins from the assembled transcriptomes by identifying sequences encoding a signal sequence using SignalP v6 (*67*). In this subset of secreted proteins, we extracted sequences with a tpm above 1000. Since toxins genes are not always expressed at such high levels, we used these highly expressed proteins as BLASTp queries (e-value 1e-3) to extract homologous proteins with lower expression. This set of likely toxin genes were clustered using CLANS (*68, 69*), which formed 44 superfamilies of proteins with high sequence similarity (Fig. S1). We extracted the combined superfamily transcript expression for each sequence read experiment (some containing multiple sequence read runs). This matrix was used as input for hierarchical clustering. The hieratical clustering was performed with the R package pheatmap with Ward’s D2 method of clustering using the Euclidean distance of the logarithm of gene superfamily expression.

#### Structural predictions

For each superfamily, we selected one representative sequence for tertiary structure prediction using Colabfold with standard settings (*70, 71*). The one structure of five predicted with highest pLDDT was used for structural similarity searches using foldseek (*72*).

#### Doppelgänger toxin identification

We used multiple approaches to identify doppelgänger toxins. First, we employed homology searches using BLASTp (*73*). We used the toxin candidates as queries against the NCBI non-redundant (nr) protein dataset with e-value of 0.1 as well as a custom database of known neuropeptides (*23, 74, 75*) as queries against the toxin candidate dataset again with e-value 0.1.

Second, we build evolutionarily informed models of selected neuropeptides to search the toxin candidate dataset with profile hidden Markov models and the custom program fuzreg. For each of the neuropeptides, we first performed homology searches using BLASTp with phylogenetically diverse representatives of the relevant phylum. For arthropod neuropeptides, we searched for orthologs in *Daphnia pulex, Ixodes scapularis, Homarus americanus, Drosophila melanogaster, Tribolium castaneum,* and *Apis mellifera*. For molluscan peptides, we searched for orthologs from *Aplysia californica, Mytilus galloprovincialis, Lottia gigantea, Pomacea canalicullata, Crassostra gigas,* and *Octopus vulgaris*. For chordate neuropeptides, we searched for orthologs in *Canis lupus familiaris, Branchiostoma floridae, Callorhinchus milii, Ciona intestinalis, Danio rerio, Gallus gallus, Lethenteron reissneri, Notamacropus eugenii, Ornithorhynchus anatinus, Petromyzon marinus, Protopterus annectens,* and *Xenopus tropicalis.* The precursor sequences were aligned using the L-INS-I method of mafft v7.520 with 1000 iterations. Based on known peptide annotations from Uniprot and previous papers (*76-78*), we extracted the peptide encoded region for model building. Profile hidden Markov models were build using hmmer v.3.4 and searched against the toxin candidate database.

As an alternative method for doppelgänger toxin identification, we build the software fuzreg (https://github.com/Thomaslundkoch/fuzreg). Fuzreg used the same alignment as the profile hidden Markov model, but builds a regular expression based on the alignment using a set of customizable heuristics. For each column in the alignment, the characters were combined to a set of allowed characters (e.g., [AGT]) and if any sequence has a gap, this position would be optional in the regular expression (?). Here, we collapsed positions where more than 3 different amino acids occurred to be any amino acid (‘.’). After the regular expression is build, we used a fuzzy search to identify sequence stretches with similarity to the query expression. The fuzzy search allows for matches with a given Levenshtein distance to the query, such that several insertions, deletions, and substitutions can be a match. The allowed Levenshtein distance used was set to 0.4 times the length of the regular expression. The matches were ranked according to an alignment score to either the human, *Lottia gigantea*, or *Homarus americanus* neuropeptide sequence. The alignment scores are show in Figure 2 as the similarity score. Following searches, sequences with similarity scores above 50 were visually inspected to determine whether they represented true or false positives. A false positive is defined as a hit that, upon manual inspection, does not appear to encode a doppelgänger toxin. Inspection focused on features such as the presence of cleavage sites that could give rise to the predicted peptide, the number of cysteines, and other relevant characteristics.

#### Clustering analyses

Using the identified superfamily 1 sequences, we used tblastn against additional assembled full body octopod and cephalopod transcriptomes with the online BLAST interface at e-value 0.1. The identified transcripts were downloaded and translated. A set of non-redundant sequences were used as input in CLANS for clustering, which performs all-against-all BLASTp with BLOSUM62 substitution matrix. The clustering creates a map of points representing individual sequences with connections, which represents sequence similarities. The minus log blast p-values were used as attractive forces in a uniformly repulsive force field to group similar sequences.

Mature octopus tachykinin-like toxins and bilaterian tachykinins were clustered as described above. The mature octopus tackykinin-like toxins were predicted as in Kanda (*11*), and mature neuropeptides were obtained from (*79*).

#### Residue-level sequence diversity

The octotensin encoding transcripts and neurotensin precursor genes were aligned using the L-INS-I method of mafft v7.520 with 1000 iterations. The conservation score for each residue was calculated with rate4site (*80*). Additionally, we calculated the moving average of the conservation score over the 5 preceding residues. The signal sequence cleavage sites were identified using SignalP v6.0.

#### Genomic and synteny analyses

Using the genomes of highest quality (*O. vulgaris, O. sinensis,* and *A. fangsiao*), we first identified the region encoding the Doppelgänger toxin genes and used available genomic annotation to identify genes upstream and downstream. This identified nuclear receptor GRF (NR-GRF) and fatty-acid binding protein/lipocalin (FABP) as the best markers of the region.

SPlign was used to identify the exon boundaries of the identified transcripts. Most of the Doppelgänger peptide ORF are split on two exons, where the second exon encode 10-20 amino acids. The SPlign analysis allowed us to split the transcripts into the precursor encoding exons, which were used for subsequent homology search. We used cd-hit (-c 0.9) to remove highly similar exons and used tblastx (e-value 0.01) with the reduced exon dataset as a query against the genomes.

#### Phylogenetic reconstruction

We extracted cytochrome oxidase subunit III nucleotide sequences from the mitochondrial genomes of the species with available genomes (Table S8) and *E. scolopes* as an outgroup. The sequences were aligned using mafft v7.520 (File S3), and a maximum likelihood tree was constructed using IQ-tree v2.2.2.3. The TIM2+F+G4 model of evolution was chosen according to the Bayesian information criterion with seed 294876.

#### Peptide prediction

The positions of the likely mature peptides were predicted using the deep learning tool DeepPeptide (*22*) using the esm2 model (*81*).

#### Glycosylation prediction

We predicted potential *O*-glycosylation sites of Ser/Thr residues in the octotensin-encoduing precursors using the online version of NetOGlyc v4.0 (https://services.healthtech.dtu.dk/services/NetOGlyc-4.0/) (*82*).

#### Neurotensin receptor evolution

In humans, neurotensin is the endogenous ligand of two G protein-coupled receptors of class A/ rhodopsin-type, NTSR1 and NTSR2, as well as a sortilin-1 receptor. Previous reports have found that fish seemingly only possess a single neurotensin receptors belonging to the NTSR1 family, whereas amphibians possess both (*34*). Since this initial report, the amount of chromosomal level genome assemblies has expanded greatly allowing a comprehensive comparative approach to investigate the evolution of NTSRs in chordates. As the likely intended target of octotensin are teleost fish NTSRs, it is crucial to know how many and what family these belong to.

To reconstruct the NTSR gene phylogeny, we searched for homologs of human NTSR1 and NTSR genes in a range of chordates, including urochordates and cephalochordates. While neither urochordates nor cephalochordates seemingly encode NTSR genes, we could identify homologs from 25 vertebrate species (species indicated in the Fig. S13 legend). Using neuromedin U receptors as an outgroup, we reconstructed the NTSR gene evolution using both Bayesian and maximum likelihood approaches (Fig. S13, File S4). Maxmimum likelihood reconstruction was performed using IQ-TREE v2.2.2.3, model of amino acid evolution was selected based on Bayesian Information Criterion from ModelFinder to be Q.mammal+G4. Branch labels represent IQ-TREE’s UFboot values. The Bayesian tree was constructed using MrBayes v3.2.7a using a mixed model of amino acid evolution. The reconstruction was run for 10,000,000 generations at which point the two chains had converged.

We could identify NTSR2 genes in placentals, amphibians, and in lungfish and coelacanths. The NTSR2 gene has seemingly been lost in many vertebrate linages, including cartilaginous and teleost fish, reptiles, birds, monotremes, and marsupials. Despite the monophyletic grouping of the two cyclostome NTSR genes with the NTSR1 family, it is possible that one of these cyclostome genes in fact belongs to the NTSR2 family, but groups with the NTSR1 gene. To investigate this possibility, we performed synteny analyses of the gene regions surrounding the NTSR genes using protein annotations from NCBI (Fig. S14). The genes LPIN1, GREB1, and ROCK1 have strong linkage to the NTSR2 gene. However, when we searched for those genes in the genomes apparently lacking the NTSR2 gene, we could not identify any potential NTSR2 gene that might have been overlooked. This strongly suggest that the NTSR2 genes were indeed lost in those lineages.

#### Peptide synthesis

OT-Ovu1, OT-Hma, OT-Eme and Octopus tachykinin I were synthesized by solid-phase peptide synthesis in 0.1 or 0.05 mmol scale using preloaded Fmoc-protected Tentagel R HMPA resin from Rapp Polymere (Tuebingen, Germany) for C-terminal carboxylic acid sequences. Fmoc-protected amino acids, coupling reagents, and solvents used for the synthesis were purchased from Iris Biotech (Marktredwitz, Germany). The synthesis was performed on a Syro I instrument from Biotage (Uppsala, Sweden). Coupling conditions were room temperature (RT) for 2 × 120 min using 5.2 equiv amino acids, 4.7 equiv N-[(1H-benzotriazol-1-yl)(dimethylamino)methylene]-N-methylmethanaminium hexafluorophosphate N-oxide, 5.2 equiv 1-hydroxy-7-azabenzotriazol, and 8 equiv N,N-diisopropylethylamine in dimethylformamide (DMF) relative to resin. Deprotection was performed at RT with 40% piperidine in DMF for 3 min followed by 20% piperidine in DMF for 15 min. Washing steps were with 2 × N-methyl-2-pyrrolidone, 1 × dichloromethane (DCM), and 1 × DMF. After completion of the peptide assembly, the peptidyl resin was washed with 3 × DCM and dried. The peptide was released from approximately 0.025 mmol resin using a mixture (2 ml) of 95% trifluoroacetic acid (TFA), 2.5% triethylsilyl, and 2.5% water over 2.5 h. For methionine-containing sequences, the 2 mL mixture was containing 87.5% trifluoroacetic acid (TFA), 2.5% triethylsilyl, 2.5% water, 2.5% thioanisole, and 5% ethanethiol, or alternatively 87.5% trifluoroacetic acid (TFA), 2.5% triethylsilyl, 2.5% water, 2.5% thioanisole, and 5% ethanethiol with approximately 0.4 mg Sodium iodide (NaI). Cold diethylether (13 ml at −20 °C) was added, and the mixture was further cooled to −85 °C for 30 min before the peptide was isolated by centrifugation. The peptide was redissolved in a mixture of TFA, water, and acetonitrile (ACN) and purified on a Dionex Ultimate 3000 HPLC system (Thermo Fisher, Waltham, USA) using a Luna C18(2) column from Phenomenex (Torrance, USA, 5 μm, 100 Å, 250 × 10 mm). A gradient of 5–100% ACN in water/0.1% formic acid (or 0.1% TFA for methionine containing sequences) was applied. The product was isolated and lyophilized by freeze-drying. The final product was analyzed by liquid chromatography (LC)–mass spectrometry (MS) on a Dionex Ultimate 3000 ultrahigh-performance LC system from Thermo Fisher connected to an Impact HD mass spectrometer (Bruker, Bremen, Germany). Glycosylated peptides were synthesized commercially by Polypeptide (USA) using glycsolyated Fmoc-L-Ser amino acids synthesized by CarboSyn (USA, Fmoc-L-Ser (Gal β(1-3)GalNAc)-OH, peracetate and Fmoc-L-Ser (α-D-GalNAc(Ac)3)-OH). All peptides were validated by HPLC and ESI-MS prior to use.

#### Chemical reagents

Unless otherwise specified, all chemical and cell culture reagents were sourced from Sigma and ThermoFisher respectively. Neurotensin and Substance P were obtained from GenScript. OT-Ovu1, OT-Hma and OT-Eme were first synthesized in house using solid phase peptide synthesis (SPPS) as detailed in “Peptide synthesis”, and larger amounts of the non-glycosylated and glycosylated peptides were further custom-synthesized by GenScript and Polypeptide (San Diego, USA), respectively. Fmoc-L-Ser(Gal(β(1-3)GalNAc) was purchased from CarboSyn (USA). The mass and purity (> 95 %) of the peptides were verified by RP-HPLC and MS prior to use.

#### Cell culture

HEK293T cells were given by Prof. Hans Braüner-Osborne (University of Copenhagen, Denmark) and were cultured in DMEM supplemented with 10% FBS (Gibco) and 100 U/mL penicillin and 100 µg/ml streptomycin at 37°C in a humidified incubator supplemented with 5% CO_2_. HTLA cells were given by Prof. Bryan Roth (University of North Carolina, Chapel Hill, US) and were maintained in Dulbecco’s modified Eagle’s medium (DMEM) supplemented with 10% Fetal Bovine Serum (FBS) (Biowest) and 100 U/mL penicillin and 100 µg/mL streptomycin at 37°C in a humidified incubator supplemented with 5% CO_2_. Constant selective pressure for tTA-dependent luciferase reporter and a β-arrestin 2 tobacco etch virus protease fusion gene were maintained with the addition of 100 µg/mL hygromycin B and 2 µg/ml puromycin (Tango growth medium). Mycoplasma testing was conducted regularly to ensure the lack of mycoplasma contamination.

#### DNA constructs

3xHA-tagged human NTSR1 construct was obtained from cDNA.org (cDNA resource center, Bloomsburg PA, US). Constructs for Dre-ntsr1 (sequence accession number: XP_009295023.1) inserted in pcDNA3.1(+) plasmid or designed as per Tango construct template (*35*) were custom-synthesized by Twist Biosciences. cAMP sensor using YFP-Epac-RLuc (CAMYEL) (*83*) was given kindly by Prof. Hans Braüner-Osborne (University of Copenhagen, Denmark). mVenus-β-arrestin 1/2 constructs were kind gifts from Dr. Tao Che (Washington University, St. Louis). C-terminally NanoLuciferase (NLuc)-tagged human NTSR1 inserted in pcDNA3.1(+) used in the β-arrestin recruitment studies were customed synthesized by GenScript. TRUPATH G protein constructs were gifts from Prof. Bryan Roth (University of North Carolina, Chapel Hill) (Addgene kit #1000000163). All NTSR1 mutants (i.e. E332A, T335A, D331A-E332A-T335A, D331W, E332W, T335W, D331W-E332W-T335W, R6.54A, R6.55A, F6.58A, W334A, Y7.28A, Y7.31A and Y7.335A) inserted in pcDNA3.1(+)-N-terminal-HA used in this study were custom synthesized by GenScript. All DNA constructs were sequenced and confirmed prior to use.

#### IPOne Gq assay

HEK293T cells were seeded onto 6-well plates at a density of 900,000 cells/well. Cells were transfected the next day with 3xHA-tagged human NTSR1, Dre-ntsr1 or NTSR1 mutants (all at 1 µg) using Polyfect (Qiagen) according to manufacturer’s protocol. 48 hours post-transfection, HEK293T cells were split using 0.05% Trypsin-EDTA solution and cell density were determined using an automated cell counter. Manufacturer’s protocol were then largely followed in next steps. Cells were resuspended in stimulation buffer that contains lithium chloride provided by the IPOne Gq activation kit (Revvity). Cells were seeded at 40,000 cells / well on white 384 Optiplate (PerkinElmer) and were stimulated with peptide ligands for 1 hour at room temperature. Equal volume of Eu- and Tb-tagged antibodies diluted in lysis buffer provided by assay kit were added on the cells and were incubated for an hour. HTRF signal were determined using PHERAstarFSX platereader (BMG Labtech). HTRF ratio were determined by dividing 665 nm signal over 620nm signal.

#### PRESTO-Tango β-arrestin recruitment assay

The assay was done as described in (*35*). Briefly, HTLA cells (1.3 million cells/well) were seeded in 6-well plates and incubated overnight. On day 2, cells were transfected with 1.5 µg receptor constructs using Polyfect (Qiagen) according to manufacturer’s protocol. On day 3, 25,000 cells in 40 µL DMEM supplemented with 1 % dialyzed FBS per well were seeded in poly-L-lysine (PLL)-coated white clear-bottom 384 well plates (Corning) and incubated overnight. On day 4, the media was changed and 10 µL of each of the peptide was added at 5x final concentration. The peptides were resuspended in HBSS with 20 mM HEPES, 1 mM of CaCl_2_, 1 mM of MgCl_2_, pH adjusted to 7.4 with 1 M NaOH and supplemented with 0.1 % BSA. After overnight incubation, on day 5, the medium and the compounds were removed from the cells and 20 µL of 1:20 dilution of BrightGlo (Promega) in HBSS buffer supplemented with 0.01 % pluronic F68 was added to the cells. Plates were incubated for 20 minutes in the dark at room temperature. Luminescence was measured on a Molecular Devices SpectraMax iD5 (Molecular Devices) with each well integrated for 1 second. Relative luminescence units (RLU) were taken as raw data for the plotting of dose-response curves.

#### CAMYEL cAMP assay

HEK293T cells were seeded onto 6-well plates at a density of 900,000 cells/well. Cells were transfected the next day with 3xHA-tagged human NTSR1 (1 µg) and CAMYEL Epac sensor (1 µg) using Polyfect (Qiagen) according to manufacturer’s protocol. 16 to 24 hours post-transfection, HEK293T cells were seeded onto white 96 well plate (PerkinElmer) coated with PLL at a density of 40,000 cells/well and were allowed to grow overnight. On the day of assay, cells were washed once with phosphate-based saline (PBS) and 80 µL of assay buffer which composed of PBS with 0.5 mM MgCl_2_, adjusted to pH 7.4, were added onto each well. 10 µL of *Renilla* luciferase substrate coelenterazine-h (Nanolight) were added onto each well and incubated for 5 minutes. 10 µL of 10 x ligands were then added manually onto the plate and BRET signal was measured instantaneously using PHERAstarFSX platereader (BMG Labtech) equipped with BRET1 475 (30 nm bandwidth) and 535 (30 nm bandwidth) filters continuously for 30 min.

#### β-arrestin 1/2 recruitment assay

HEK293T cells were seeded onto 6-well plates at a density of 900,000 cells/well. Cells were transfected the next day with C-terminally-NLuc-tagged human NTSR1 (0.3 µg) and mVenus-β-arrestin 1/2 constructs (0.3 µg) at 1:1 ratio using Polyfect (Qiagen) according to manufacturer’s protocol. 16 to 24 hours post-transfection, HEK293T cells were seeded onto white 96 well plate (PerkinElmer) coated with PLL at a density of 40,000 cells/well and were allowed to grow overnight. On the day of assay, cells were washed once with PBS and 80 µL of assay buffer which composed of PBS with 0.5 mM MgCl_2_, adjusted to pH 7.4, were added onto each well. 10 µL of NLuc substrate furimazine (Promega) were added onto each well and incubated for 3 minutes. 10 µL of 10x concentrated ligands were then added manually onto the plate and BRET signal was measured instantaneously using PHERAstarFSX platereader (BMG Labtech) equipped with BRET1 475 (30 nm bandwidth) and 535 (30 nm bandwidth) filters continuously for 30 min.

#### TRUPATH G protein dissociation assay

The following protocol was adopted from (*84*). In brief, HEK293T cells were seeded onto 6-well plates at a density of 900,000 cells/well. Cells were transfected the next day with 3xHA-tagged human NTSR1 or NTSR1 mutants (0.3 µg), RLuc8-tagged G protein constructs (0.2 µg), pcDNA3.1-Gβ_3_ (0.2 µg), pcDNA3.1-Gγ_2_-GFP2 (0.2 µg) (at a ratio of 1.5:1:1:1 and the total DNA amount per well was 0.9 µg) using Polyfect (Qiagen) according to manufacturer’s protocol. 16 to 24 hours post-transfection, HEK293T cells were seeded onto white 96 well plate (PerkinElmer) coated with poly-L-lysine (PLL) at a density of 40,000 cells/well and were allowed to grow overnight. On the day of assay, cells were washed once with PBS and 80 µL of assay buffer which composed of PBS with 0.5 mM MgCl_2_, adjusted to pH 7.4, were added onto each well. 10 µL of Rluc8 luciferase substrate Prolume Purple (Nanolight coorperate) were added onto each well and incubated for 5 minutes. 10 µL of 10x concentrated ligands were then added manually onto the plate and BRET signal was measured instantaneously using PHERAstarFSX platereader (BMG Labtech) equipped with BRET2 410 (80 nm bandwidth) and 515 (30 nm bandwidth) filters continuously for 30 min.

#### BRET ratio calculation

BRET1 ratio was determined by dividing the BRET signal at 535 nm over that at 475 nm. BRET2 ratio was determined by dividing the BRET signal at 515 nm over that at 410 nm. The net BRET ratio was determined by subtracting the BRET ratio with drugs with the BRET ratio of vehicle control. Net BRET ratio at 30-min time points for each ligand concentration tested were determined and plotted against log(concentration) of drugs tested. Results of test ligands were normalized as percentage of maximal neurotensin response. Concentration response curves were fitted with a 4-parameter logistic equation (equation 1) using GraphPad Prism (v.10.6.0):

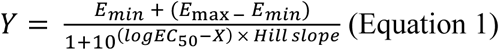

#### G protein bias quantification

The methods used in (*85*) to quantify G protein bias was largely followed. In brief, neurotensin was chosen to be the physiological reference ligand given its physiological significance as well as its full agonism at human NTSR1. The operational model (equation 2) was used to determine the transduction ratios (τ/K_A_).

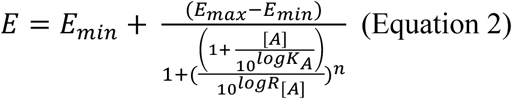

where E is the effect of the ligand, [A] is the concentration of test ligands, E_max_ is the maximal response of the system, E_min_ is the minimum response of the system, logK_A_ is the logarithm of the equilibrium dissociation constant, n is the slope of the transducer function, log R is the logarithm of the transduction ratio (τ/K_A_). The operational model was manually input in GraphPad Prism (v.10.6.0) according to (*85*). Each experiment was analyzed using the operational model which all individual experiments were fitted with a global shared slope. The LogR (equivalent to log[τ/K_A_]) values were estimated by GraphPad Prism (v 10.6.0) and were used to calculate the relative efficacy (RE) using equation 3 and 4.

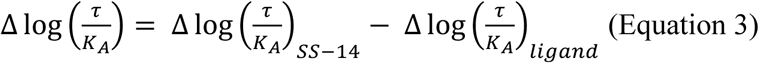

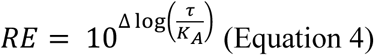

S.E.M. of 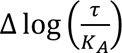 were calculated using the following equation 5:

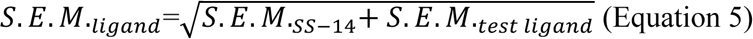

#### Expression and Purification of NTSR1

NTSR1-containing were expressed in Sf9 cells and pellets were snap-frozen in liquid nitrogen for storage. To start the purification of NTSR1, the cell pellets were thawed and subsequently lysed in a hypotonic lysis buffer containing 10 mM HEPES at pH 7.5, 1 mM benzamidine, 1 mM EDTA, 1 mM MgCl_2_, protease inhibitor cocktail, Pierce Universal Nuclease, and 100 μM TCEP, and stirred at 100 rpm for 1 hour at 4°C. Membranes were harvested via ultracentrifugation at 100,000xg for 35 minutes. Supernatant was discarded, and the pellets were resuspended in a solubilization buffer containing 20 mM HEPES pH 7.5, 500 mM NaCl, 1 mM MgCl_2_, protease inhibitor cocktail, 1 mM benzamidine, Pierce Universal Nuclease, and 100 μM TCEP. While gently stirring at 4°C, detergent was added dropwise to the resuspended pellets to a final concentration of 1% LMNG/0.1% CHS/0.1% Sodium Cholate. After stirring for 3 hours at 4°C, ultracentrifugation was done at 100,000xg for 35 minutes to remove any insoluble debris. The supernatant was supplemented with 20 mM imidazole and loaded over a TALON resin column that had been washed with 10 column volumes of a buffer containing 20 mM HEPES pH 7.5, 500 mM NaCl, 20 mM imidazole, and 0.1% LMNG/0.01% CHS. Protein was washed then and eluted by loading 2 column volumes of buffer containing 20 mM HEPES pH 7.5, 250 mM NaCl, 250 mM imidazole, 0.01% LMNG/0.001% CHS, and 10% glycerol over the column. Purified NTSR1 was concentrated down to 500 μL and subjected to SEC chromatography with a Superdex 200 column in buffer containing 20 mM HEPES pH 7.5, 100 mM NaCl, 5% glycerol, and 0.01% LMNG/0.001% CHS. Peak fractions were pooled together, concentrated, supplemented with 5% glycerol, and snap frozen with liquid nitrogen for use with complexation.

#### Expression and Purification of Dominant-Negative G_i3_

Purifying dominant negative (DN) G_i3_ began with thawing and resuspending cell pellets in lysis buffer containing 20 mM HEPES pH 7.5, 1 mM EDTA, 5% glycerol, 1 mM MgCl_2_, 5 mM β-mercaptoethanol, 100 µM GDP, protease inhibitor cocktail, and Pierce Universal Nuclease, and gently stirring for 30 minutes at 4°C. Resulting lysate was harvested via ultracentrifugation at 100,000xg for 35 minutes. Supernatant was discarded and pellets were resuspended in a solubilization buffer containing 20 mM HEPES pH 7.5, 100 mM NaCl, 1 mM MgCl_2_, 5% glycerol, 1% sodium chocolate, 100 µM GDP, protease inhibitor cocktail, Pierce Universal Nuclease, and 5 mM β-mercaptoethanol. Gentle stirring at 4°C for 1 hour was followed by ultracentrifugation for 35 minutes at 100,000xg, and the solubilized protein in the supernatant was supplemented with 30 mM imidazole and incubated with Ni-NTA beads for 1 hour on ice. Nickel beads were harvested via centrifugation at 300xg and subsequently packed into a chromatography column. The column was subjected to a series of washes of 10 column volumes with buffers containing 30 mM imidazole with 50% solubilization/50% E2 buffer, 25% solubilization/75% E2 buffer, 12.5% solubilization/87.5% E2 buffer, and 100% E2 buffer (E2 buffer consisted of 20 mM HEPES pH 7.5, 100 mM NaCl, 100 µM GDP, 1 mM MgCl_2_, 5% glycerol, 5 mM β-mercaptoethanol, and 0.05% LMNG/0.005% CHS). DN G_i3_ was eluted from the column with 3 column volumes of E2 buffer and 250 mM imidazole, followed by incubation in a 3,500 Da MWCO dialysis bag with 1 mg of 3C protease per 50 mg of G-protein for overnight dialysis at 4°C. The following day, the Ni-NTA column was washed with 10 column volumes of E2 buffer and 30 mM imidazole, and overnight cleavage was loaded over the column and flow through was collected. Two additional column volumes of E2 buffer and 30 mM imidazole were washed over the column, and flow through was collected. G-protein was concentrated down to 500 µL in a 30 kDa MWCO spin concentrator and subjected to SEC chromatography using a Superdex 200 column in a buffer with 20 mM HEPES pH 7.5, 100 mM NaCl, 20 µM GDP, 100 µM TCEP, 5% glycerol, 1 mM MgCl_2_, and 0.01% LMNG/0.001% CHS.

#### Expression and Purification of scFv16

scFv16 was purified as described previously (*86*) and purified scFv16 was concentrated to 13 mg/mL and snap frozen with liquid nitrogen in buffer containing 20 mM HEPES pH 7.5, 100 mM NaCl, and 15% glycerol for future use.

#### Formation and Purification of NTSR1-Octotensin-G-protein-scFv16 Complex

NTSR1 and octotensin were incubated together on ice for 1 hour. Complex formation began with mixing octotensin-bound receptor with a molar excess of G-protein and incubating for 1 hour on ice. scFv16, apyrase was added to remove GDP from the G-protein, and HRV 3C protease was added to cut the GFP tag off of the NTSR1, prior to overnight incubation on ice. The complex was diluted 5-fold with a buffer consisting of 20 mM HEPES pH 7.5, 100 mM NaCl, 100 μM octotensin, and 0.01% LMNG/0.001% CHS. Diluted complex was loaded over an M2 flag resin column and washed with buffer containing 20 mM HEPES pH 7.5, 100 mM NaCl, 10 μM octotensin, and 0.005% LMNG/0.0005% CHS. NTSR1-Octotensin-G-protein complex was eluted from the column with a buffer of 20 mM HEPES pH 7.5, 100 mM NaCl, 10 μM octotensin, and 0.005% LMNG/0.0005% CHS. Eluent is concentrated to a lower volume and loaded onto a size exclusion Superdex 200 column with a buffer of 20 mM HEPES pH 7.5, 100 mM NaCl, 10 μM octotensin, and 0.001% LMNG/0.0001% CHS/0.00033% GDN. Peak fractions of the complex were concentrated for cryo-EM analysis.

#### Cryo-EM Sample Prep and Imaging

Cryo-EM samples were prepared with a Vitrobot Mark IV held at 100% humidity and 4°C by application of 3 μL of sample to UltrAuFoil R1.2/1.3 300 mesh grids glow discharged in a Pelco unit for 45 seconds at 10 mA and plunge frozen in liquid ethane. Grids were imaged on a G4 Titan Krios at 300 kV with a K3 direct electron detector and a Bioquantum energy filter.

#### Data Processing and Model Building

All data was processed in cryoSPARC (*87*) beginning with patch motion correction and patch CTF estimation. Template-based picking with existing GPCR-G-protein complex templates was employed, particles extracted, and a single round of 2D classification was used to remove obvious junk particles. Further particle cleaning was performed with iterative rounds of *ab-initio* model generation with multiple classes and heterogenious refinement. Clean particles were subjected to initial non-uniform refinement, CTF refinement and reference-based motion correction before final non-uniform and receptor/Ras domain local refinement. Initial models came from PDB:6OSA and 6OS9 for most components (*37*) while dominant negative Gi3 came from PDB:7T10 (*88*). Manual model building was performed in Coot (*89*) with iterative rounds of real space refinement in Phenix (*90*).

#### Cardiovascular Assessment in Anesthetized Rats

*In vivo* experiments were conducted by Inotiv at an AAALAC International-accredited facility (AAALACi Fully Accredited), under OLAW Assurance #D16-00141 and USDA registration 43-R-0112. All procedures were performed in accordance with applicable institutional and federal guidelines for the care and use of laboratory animals. Male Sprague Dawley rats (8–10 weeks of age at study initiation) were obtained from Envigo. Animals were housed under controlled environmental conditions (temperature 72 ± 8 °F; relative humidity 30–70%) with a 12-hour light/dark cycle (lights on 6:00 AM–6:00 PM) and provided ad libitum access to chow (5L0B Rodent Diet 20) and water. Body weight was recorded prior to anesthesia. Animals were randomly assigned to one of three groups (n = 3 per group): vehicle control, test compound 1 (representing OT-Ovu1g), or Test Compound 5 (representing OT-Ovu1). Test compounds were administered as a single intravenous bolus at a dose of 20 nmol/kg in a dosing volume of 0.8 mL/kg via the tail vein. Test compounds were received as a lyophilized powder and stored at -20 °C until use. On the day of dosing, compounds were prepared fresh in sterile saline (0.9% NaCl; Braun). To prepare a 25 nmol/mL dosing solution, lyophilized peptide (30 nmol) was equilibrated to room temperature, briefly centrifuged, reconstituted in saline, and vortexed to ensure complete dissolution before final volume adjustment. Animals were anesthetized with Inactin (100 mg/kg, intraperitoneal). Body temperature was maintained using a heating pad throughout the procedure. Under sterile conditions, animals were placed in the supine position, and the carotid artery was surgically exposed, ligated distally, and cannulated with a fluid-filled catheter for direct arterial pressure measurement. Arterial blood pressure (systolic, diastolic, and mean arterial pressure) and heart rate were continuously recorded via the carotid catheter connected to a pressure transducer. Data were acquired using Ponemah™ software (DSI). Baseline cardiovascular parameters were recorded for 15 minutes prior to dosing, followed by continuous monitoring for 15 minutes after test article administration.

#### Mouse husbandry

Male and female mice in aged matched controlled cohorts from 5-10 weeks of age were used for all experiments and were obtained from Inotiv (Envigo). Male and female mice were used in the tail flick assay, as no sex differences were observed we moved forward with paw incision assay with male mice. Outbred CD-1 (also known as ICR) mice are commonly used in pharmacology and biomedical research. Mice were allowed to recover for at least 5 days after shipment before use in experiments. The mice were kept in an Association for Assessment and Accreditation of Laboratory Animal Care–accredited vivarium at the University of Texas Southwestern Medical Center under temperature control and 12-hour light/dark cycles with food (standard lab chow) and water available *ad libitum.* No more than five same sex mice were kept in a cage. The animals were monitored daily, including after surgical procedures, by trained veterinary staff. All experiments performed were in accordance with Institutional Animal Care and Use Committee–approved protocols at the University of Texas Southwestern Medical Center according to the guidelines of the National Institutes of Health (NIH) *Guide for the Care and Use of Laboratory Animals* handbook.

#### Mice behavioral experiments

Before any behavioral experiment or testing, the animals were brought to the testing room in their home cages for at least 1 hour for acclimation. Testing always occurred within the same approximate time of day between experiments, and environmental factors (noise, personnel, and scents) were minimized. All testing apparatus (cylinder, grid boxes, etc.) were cleaned between uses. The experimenter was blinded to treatment group by another laboratory member delivering coded drug vials, which were then decoded after collection of all data

#### Paw incision and mechanical allodynia assessment

Mechanical thresholds were determined before surgery using calibrated Von Frey filaments (Stoelting Co.) with the “up-down” method and four measurements after the first response per mouse. The size range of stimuli was between 2.44 (0.4 mN) and 4.56 (39.2 mN).

The starting filament was 3.61 (3.9 mN). The filament was placed perpendicular to the skin with a slowly increasing force until it bent; it remained bent for approximately 1 second and was then removed. Data were analyzed using the nonparametric method of Dixon, as described by Chaplan et al. (*91*). The mice were housed in a Von Frey apparatus (Bioseb invivo instruments) with Plexiglas walls and ceiling and a wire mesh floor. The surgery was performed by anesthesia with ∼2-5% isoflurane in 2% oxygen, preparation of the right plantar hind paw, sterilized with iodine and 70% ethanol 3 times, and a 5-mm incision made through the skin and fascia with a no. 10 scalpel. The muscle was elevated with curved forceps, leaving the origin and insertion intact, and the muscle was split lengthwise using the scalpel to not sever the tendon. 300 uL 1 mg/kg gentamycin was then administered. The wound was then closed with 5-0 nylon sutures. The next day, the mechanical threshold was again determined as described above, and OT-Ovu1g, OT-Ovu1and vehicle (sterile water) were given via intrathecal (i.t.) injection at 200 nmol dose (in a volume of 5 uL). No animals were excluded from these studies. Mechanical threshold compared to vehicle at both the 30-, 60-, 90-, and 120-minute time points were measured. A Two-way ANOVA with multiple comparisons was performed. Results correspond to means ± SEM.

#### Statistical analysis

Data analyses were performed in GraphPad Prism (v.10.2.3). One-way ANOVA with Tukey’s multiple comparison test was performed for the results of the pharmacological assay. Two-way ANOVA with Tukey’s multiple comparison test was performed for the results of paw incision model experiments. Statistical significance was taken as p < 0.05.

### Supplementary Text

#### Note 1. Cephalopod diet and predatory behavior

Understanding the dietary habits and predatory behaviors of cephalopods is crucial in unraveling the complexities of their adaptation to varied ecosystems. Employing diverse methods for prey detection, such as stomach content analysis, prey remains in middens, direct observations, isotopic assessment, trophic tracing, and molecular prey identifications, provides a comprehensive view of their feeding ecology.

##### Decapods

*Sepia officinalis*, an opportunistic feeder off the coast of Portugal, displays a diverse diet encompassing approximately 50 prey items from various taxa, including polychaetes, cephalopods, crustaceans, bivalves, gastropods, and teleosts (*92*). Noteworthy is the shift in diet based on size, with smaller individuals favoring crustaceans, while larger ones show a preference for teleost fish. Geographical variations in diet patterns have been observed, emphasizing the influence of local conditions (*93*).

The ommastrephid squid *Illex coindetii*, inhabiting the Mediterranean, mirrors the diet pattern observed in *S. officinalis*, featuring a combination of crustaceans, teleost fish, and fish. Larger individuals exhibit a preference for fish, while smaller ones focus on crustaceans. Seasonal variations add complexity, with a winter shift towards increased consumption of crustaceans based on stomach contents and isotopic analyses (*94*).

##### Octopods

In a comprehensive four-year observation, *Octopus bimaculoides* demonstrated a diverse diet, engaging with 59 different prey species, including crustaceans, mollusks, and teleost fish. Among its prey, crabs emerged as the preferred choice, with diet variations linked to habitat preferences, such as exclusive mussel consumption for those residing on mussel beds (*95*).

*Octopus vulgaris* off the coast of South Africa showcases a diet primarily composed of crustaceans, followed by mollusks, fish, and polychaetes. Larger individuals exhibit an increased proportion of teleost fish in their diet (*96*). Diet variations based on location, depth, and habitat underscore the adaptability of these cephalopods to their surroundings. Octopus caught at exposed reefs at low tide showed much higher proportions of mussel in their stomach contents (*97*). *O. vulgaris* caught at depth (>15m) tend to show a higher proportion of teleost fish in their diet in South Carolina, western Mediterranean, and at the coast of northwestern Africa (*96*)

*Octopus maorum*, residing off the coast of southeast Tasmania, demonstrates specialization, with certain individuals favoring either crustaceans or fish. Stomach content analysis further reveals individual variations in dietary preferences (*98*). In Northern China, *Octopus minor*’s dietary preferences were investigated using DNA barcoding of stomach contents, indicating a primary focus on fish consumption, with minor contributions from cephalopods and crustaceans (*99*). *Octopus insularis*, prevalent in the western Gulf of Mexico, exhibits a diet primarily centered around crustaceans, with infrequent occurrences of fish and mollusks based on stomach content analysis. Seasonal variations in the diet of *Octopus minor* in Chile further highlight the dynamic nature of cephalopod feeding behaviors (*100*).

##### Specialized Behaviors

Beyond their diverse diets, cephalopods exhibit specialized hunting behaviors. For instance, some species, like *Eledone cirrhosa* and *O. vulgaris*, employ drilling techniques to incapacitate prey (*101, 102*). The approach strategies of *O. bimaculoides* and larger Pacific striped octopus vary depending on the prey item, showcasing rapid approaches for crabs and slower tactics for shrimps (*4, 103*). Additionally, observations of *O. insularis* highlight its ability to attack visually encountered small fish using rapid arm projections and jetting maneuvers (*104*).

#### Note 2. Neuropeptide abbreviations

##### Chordate peptide abbreviations

AGRP: agouti-related peptide
Apelin: apelin
B-end: beta-endorphin
Calc: calcitonin
CART: cocaine- and amphetamine-regulated transcript protein
CCK: cholecystokinin
CGRP: calcitonin gene related peptide
EKA: endokinin A
EKC: endokinin C
GAL: galanin
GCG: glucagon
GHR: ghrelin
GIP: glucose-dependent insulinotropic polypeptide
GLIB: gonadoliberin
GLP-1: glucagon-like peptide 1
GLP-2: glucagon-like peptide 2
GRP: gastrin-releasing peptide
IAPP: islet amyloid polypeptide
Ins-A: insulin A-chain
Ins-B: insulin B-chain
Kiss: kisspeptin
L-enk: Leu-enkephalin
M-enk: Met-enkephalin
MCH: melanin concentrating hormone
MSH: melanocyte-stimulating hormone
NaPA: natriuretic peptide A
NaPB: natriuretic peptide B
NapC: natriuretic peptide C
NKA: neurokinin A
NKB: neurokinin B
NMB: neuromedin B
NMN: neuromedin N
NMU: neuromedin U
NP-AF: neuropeptide AF
NP-SF: neuropeptide SF
NPS: neuropeptide S
NPW: neuropeptide W
NPY: neuropeptide Y
NTS: neurotensin
ORX-a: orexin A
ORX-b: orexin B
OS: obestatin
OXT: oxytocin
PACAP: pituitary adenylate cyclase-activating polypeptide
PHM: peptide histidine methioninamide
PPP: pancreatic polypeptide
QRFP: orexigenic neuropeptide QRFP
SLIB: somatoliberin
SP: substance P
SST: somatostatin
UT2B: urotensin II b
VIP: vasoactive intestinal peptide

##### Molluscan peptide abbreviations

ACH: achatin
AKG: adipokinetic hormone
Ast-C: allatostatin C
AT: allatotropin
Buc: buccalin
CP: conopressin
FMRFa: FMRFamide
FRYa: FRYamide
Fuc: fucculin
GGNG: GGNG
HFAa: HFAamide
L11: elevenin
LRF: LRF
Luqin: luqin
Mm1: myomodullin 1
Mm2: myomodullin2
NdW: NdWFamide
NPK: Neuropeptide KY
NPY: neuropeptide Y
PH4: prohormone 4
Pleu: pleurin
QSG: QSG
RFa: RFamide
sCAP: small cardioactive peptide
Sensorin: sensorin
SIFa: SIFamide
SuK: sulfakinin
TK: tachykinin
Wh: whitnin
WWS: WWS

##### Crustacean peptide abbreviations

ACP: adipokinetic hormone
Ast-A: allatostatin-A
Ast-B: allatostatin-B
Ast-C: allatostatin C
Ast-CC: allatostatin-CC
CCAP: crustacean cardioactive peptide
CCHa 1: CCHamide 1
CCHa 2: CCHamide 2
CCRF: CCRFamide
CNMa: CNMamide
CRZ: corazonin
DH31: diuretic hormone 31
DH44: diuretic hormone 44
ELFa: ELFamide
FMRFa: FMRFamide
L11: elevenin
MyoS: myosupressin
NPF: neuropeptide F
PDH: pigment dispersing hormone
Proct: proctolin
PSCK: periviscerokinin
RYa: RYamide
SIFa: SIFamide
sNPF: short neuropeptide F
SuK: sulfakinin
TK: tachykinin
Tris: trissin
VT: vasotocin

---

**Fig. S1.**
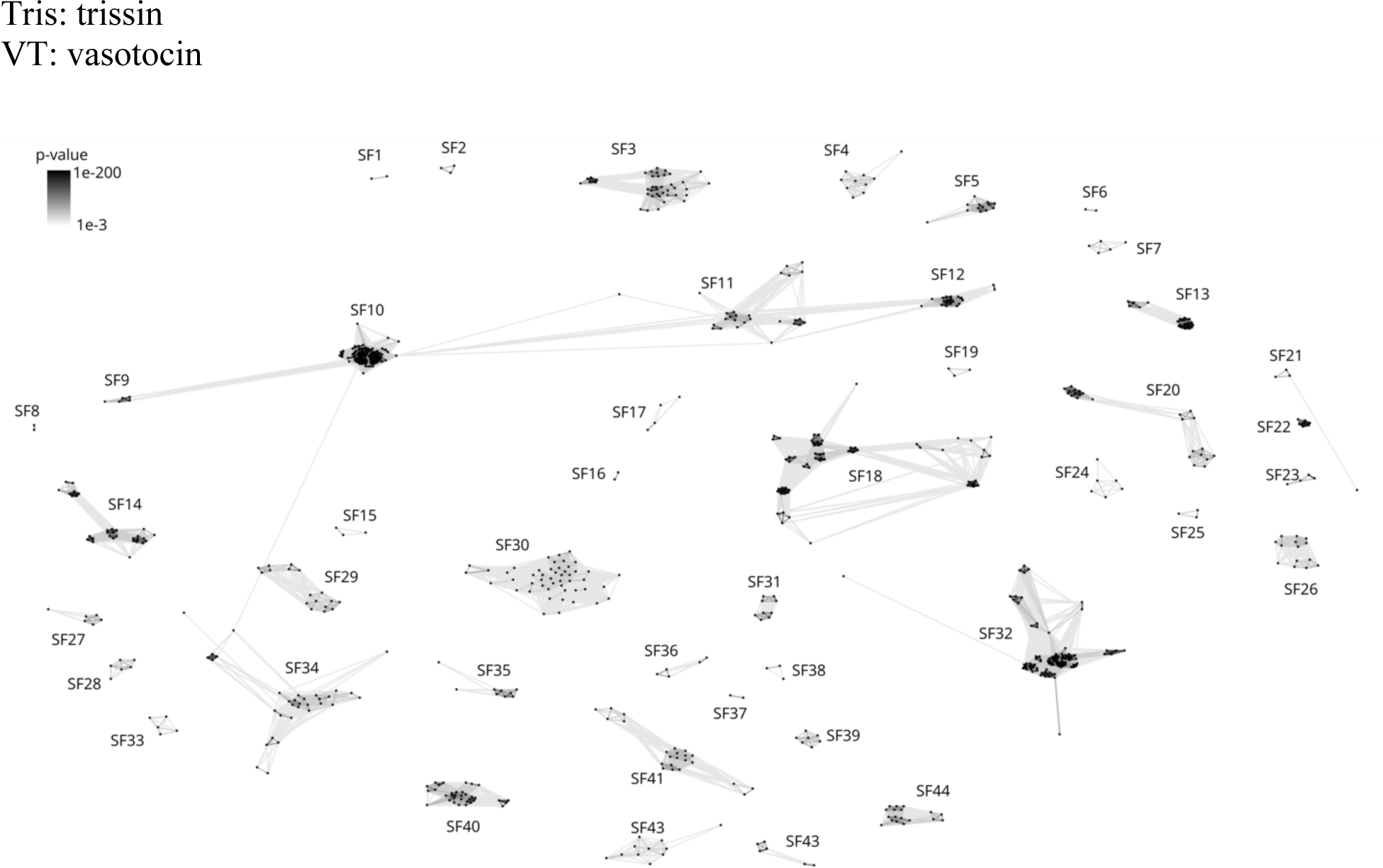
BLOSUM62 cluster map of highly expressed secreted proteins from cephalopod anterior and posterior salivary glands. Each node represents the full precursor of highly expressed salivary gland proteins and homologs. The connecting edges show the BLASTp p-value above 1e-3. The labels show the defined superfamilies.

**Fig. S2.**
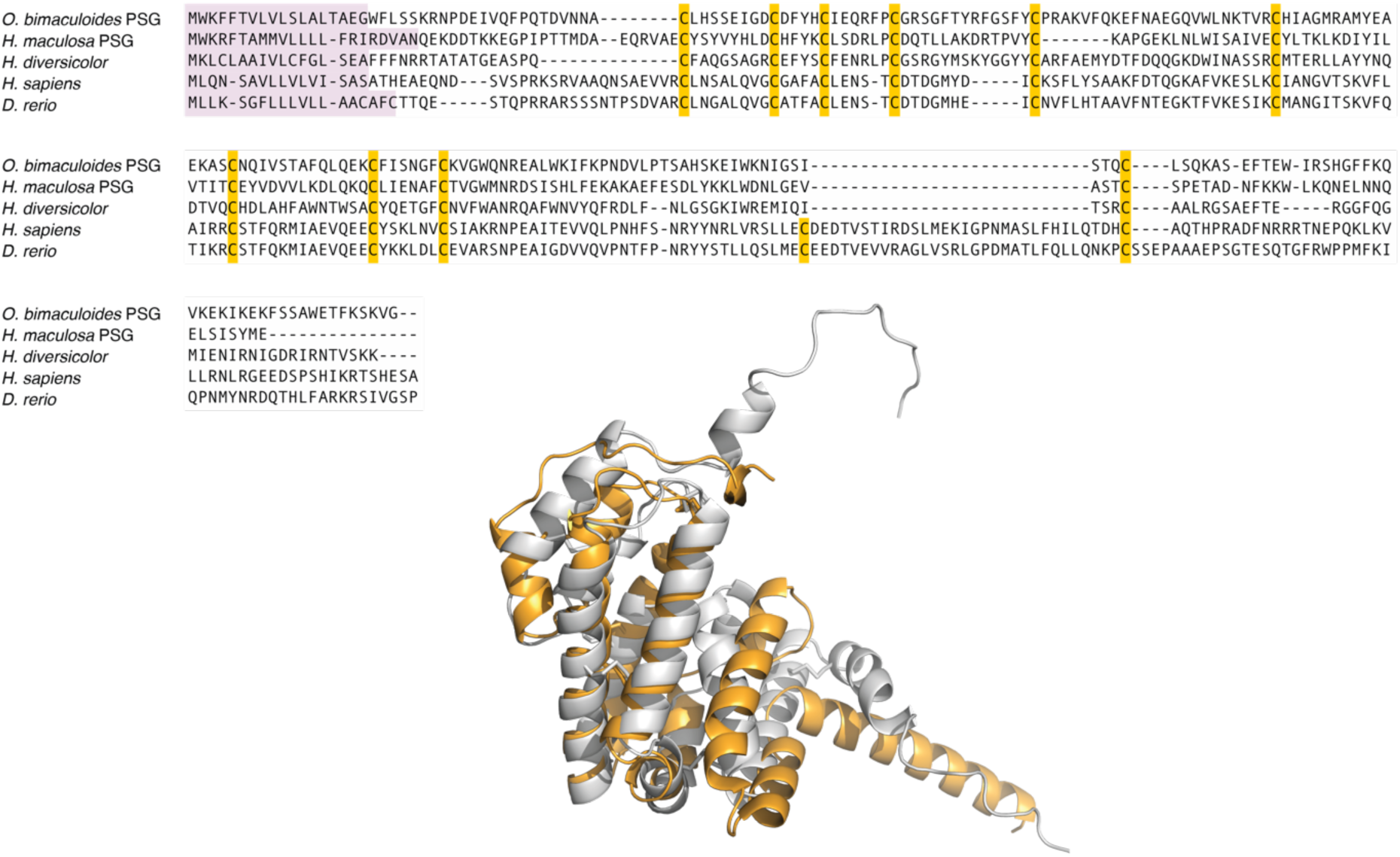
Stanniocalcins doppelgänger toxins from cephalopod PSG. Top: Sequence alignment of stanniocalcin doppelgänger toxins from *O. bimaculoides* and *H. maculosa* and human stanniocalcin 1 and 2. Stanniocalcin is a large signaling protein stretching from the end of the signal sequence to the C-terminus of the precursor. Cysteine residues are highlighted in yellow. Bottom: Overlay of their Alphafold predicted structures of human stanniocalcin-2 (Uniprot: O76061) with identified *O*. *bimaculoides* stanniocalcin shows their close similarities.

**Fig. S3.**
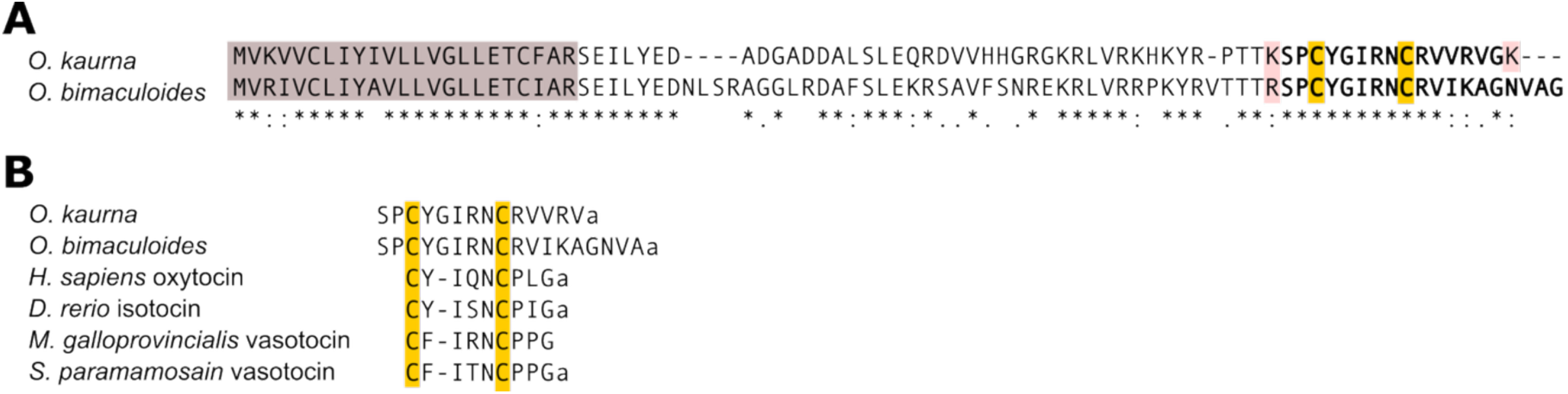
Potential oxytocin doppelgänger toxin from cephalopod superfamily 24 warranting further characterization. (A) Sequence alignment of the two precursors of SF24. The signal sequence is highlighted in brow, putative cleavage sites in red and cysteines in yellow. The likely doppelgänger toxins are bolded. The *O. kaurna* and *O. bimaculodies* transcripts are both highly expressed at 1016 and 2847 tpm, respectively. (B) Multiple sequence alignment of potential toxins with human mature oxytocin, zebrafish (*D.rerio*) isotocin, the mussel, *Mytilus gallopovincialis,* vasotocin, and the mud crab *Scylla paramamosain* vasotocin. ‘a’ denotes amidation.

**Fig. S4.**
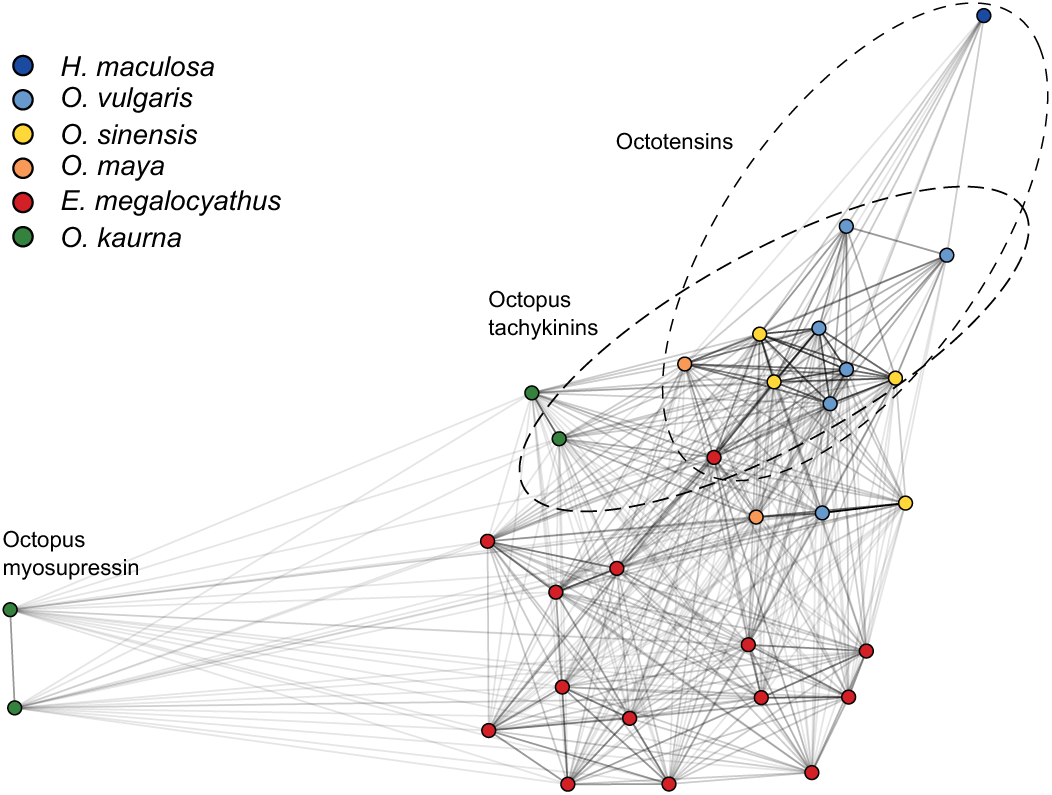
**Cluster** analysis of SF1 precursors. Graph-based clustering of the SF1 precursor sequence similarity calculated as the p-value from BLASTp using BLOSUM62 scoring matrix.

**Fig. S5.**
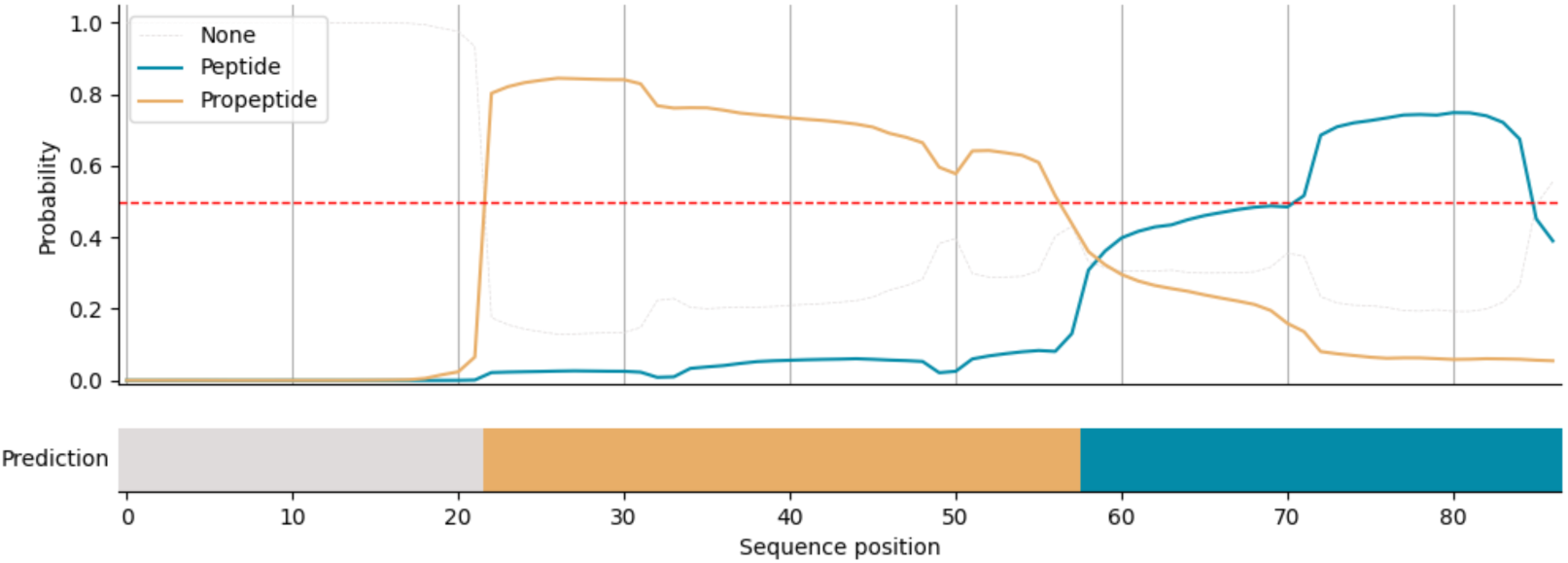
DeepPeptide prediction of peptide and propeptide regions of OT-Ovu1 precursor. The lines represent the predicted marginal probabilities, and the colored bars indicated the predicted peptide and propeptide regions along the top scoring path. DeepPeptide predicts OT-Ovu1 to be the mature peptide region (position 72-86).

**Fig. S6.**
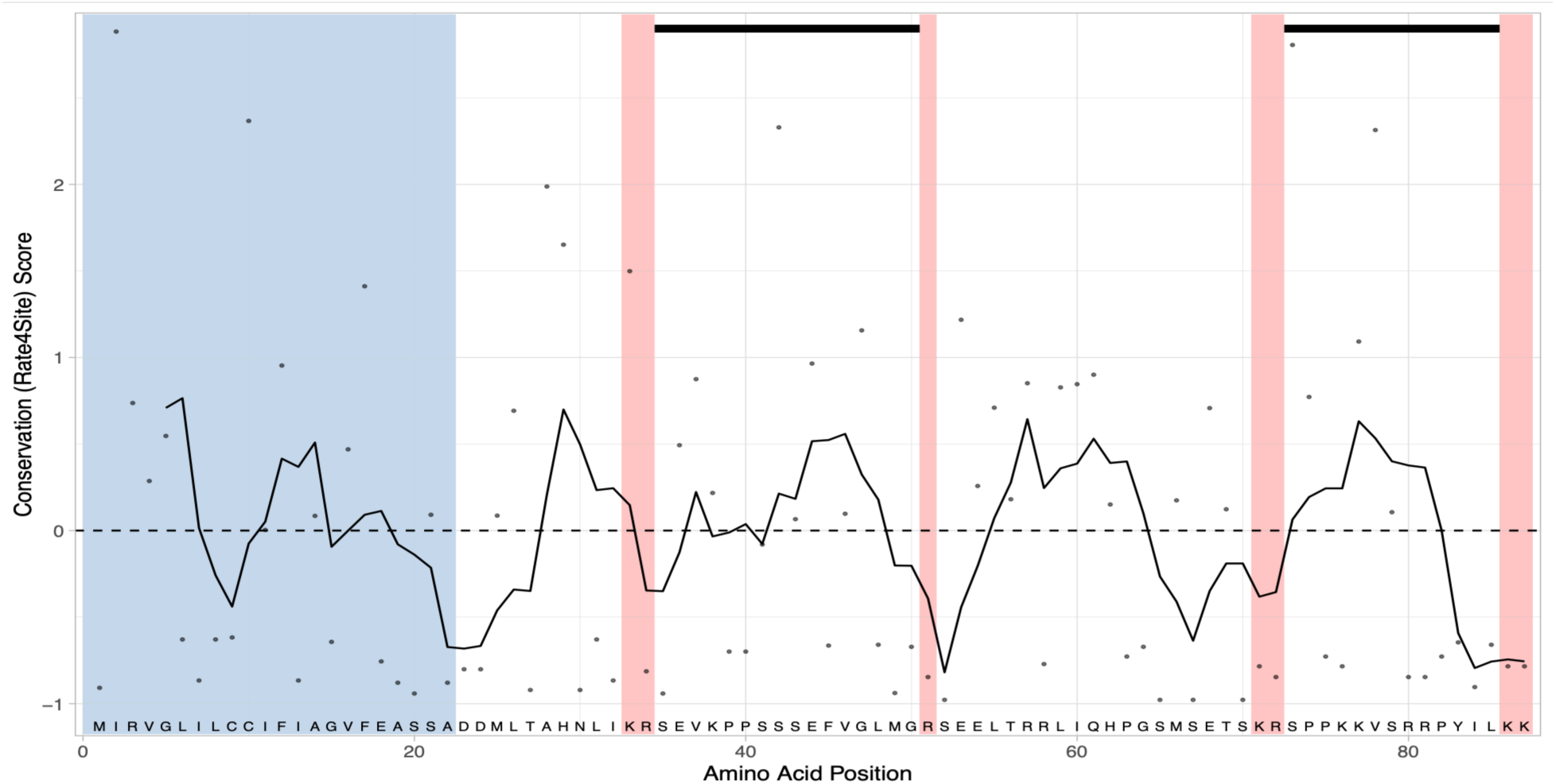
Evolutionary trace analysis of octotensin-encoding SF1 precursors. The conservation (rate4site) score was calculated for each position of an alignment of octotensin precursors. A high score shows elevated rate of evolution, whereas a low score indicates highly conserved residues. The solid line shows a moving average of the 5 preceding residues. The scores are mapped onto a single precursor sequence shown at the bottom. The signal sequence region is highlighted in blue and the predicted cleavage sites are show in red. The octotensin-encoding region have highly conserved residues in the C-terminus.

**Fig. S7.**
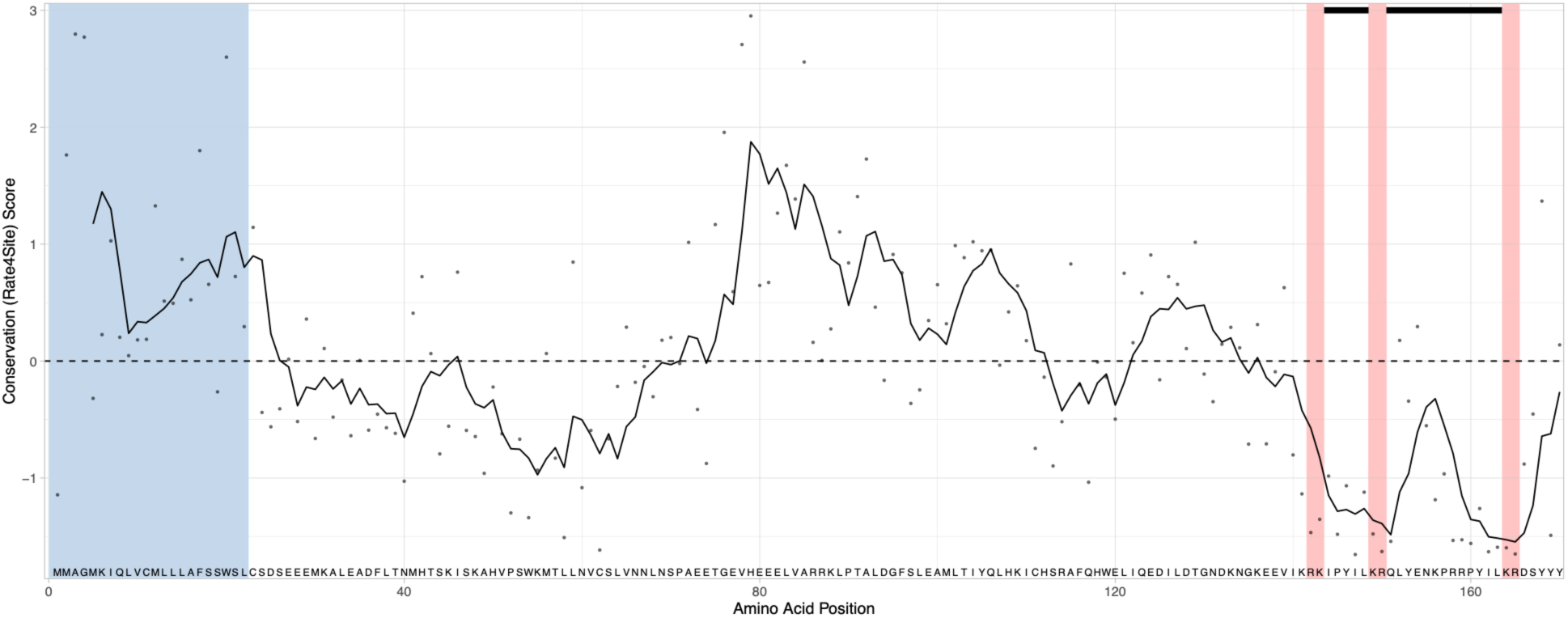
Evolutionary trace analysis of vertebrate neurotensin- and neuromedin N-encoding precursors. The conservation (rate4site) score was calculated for each position of an alignment of vertebrate neurotensin- and neuromedin N-encoding precursors. A high score shows elevated rate of evolution, whereas a low score indicates highly conserved residues. The solid line shows a moving average of the 5 preceding residues. The scores are mapped onto a single precursor sequence (human) shown at the bottom. The signal sequence region is highlighted in blue and the predicted cleavage sites are show in red. The neuromedin-N and neurotensin encoding regions are overlined in the C-terminal region of the precursor.

**Fig S8.**
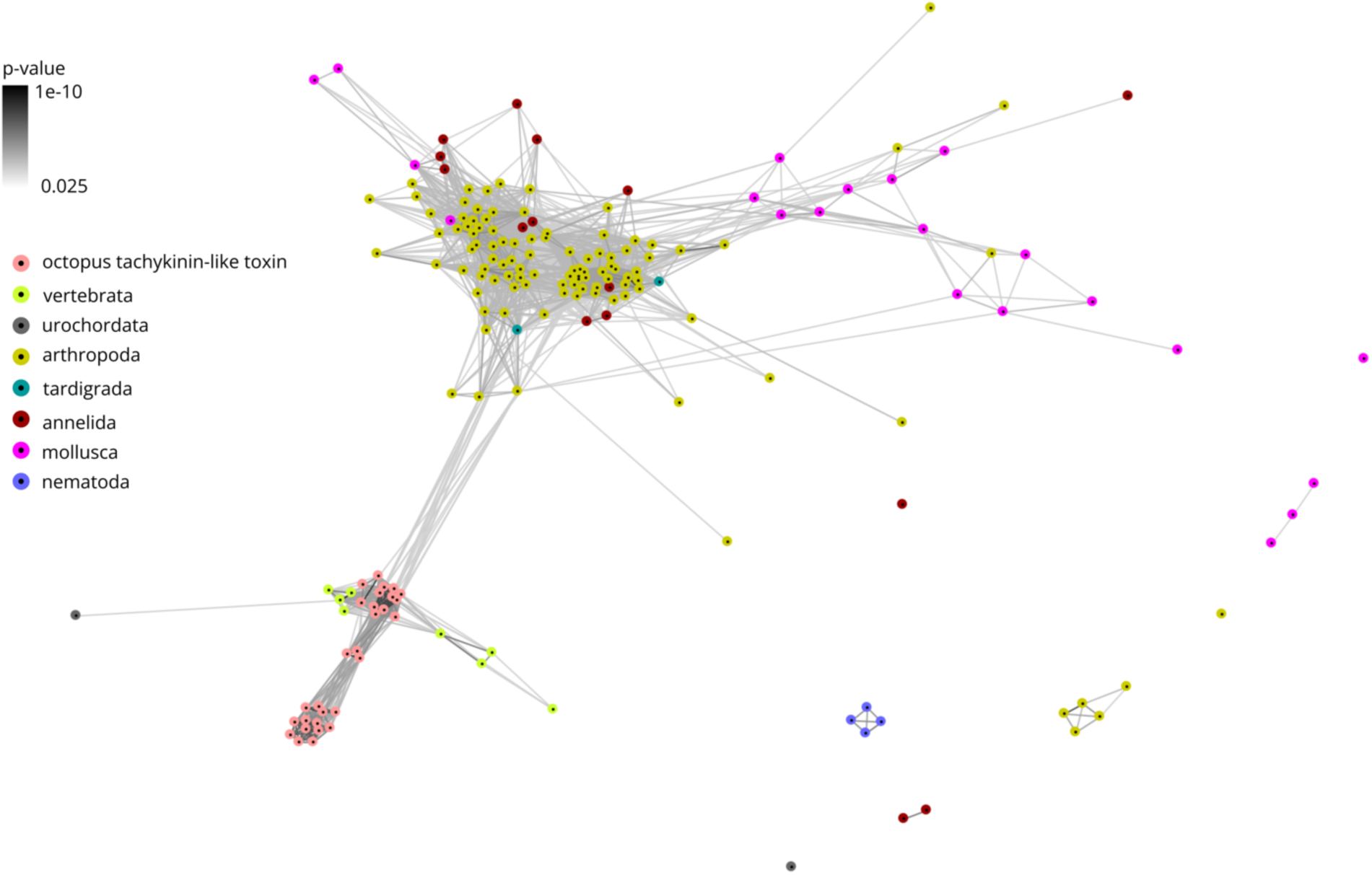
BLOSUM62 cluster map of bilaterian tachykinins and octopus tachykinin-like toxins. The depicted nodes represent individual mature neuropeptides and octopus tachykinin-like toxins and edges show BLASTp p-values > 0.025. Nodes are colored according to phylum. The octopus tachykinin-like toxins in the lower left corner cluster closely with vertebrate mature tachykinins.

**Fig. S9.**
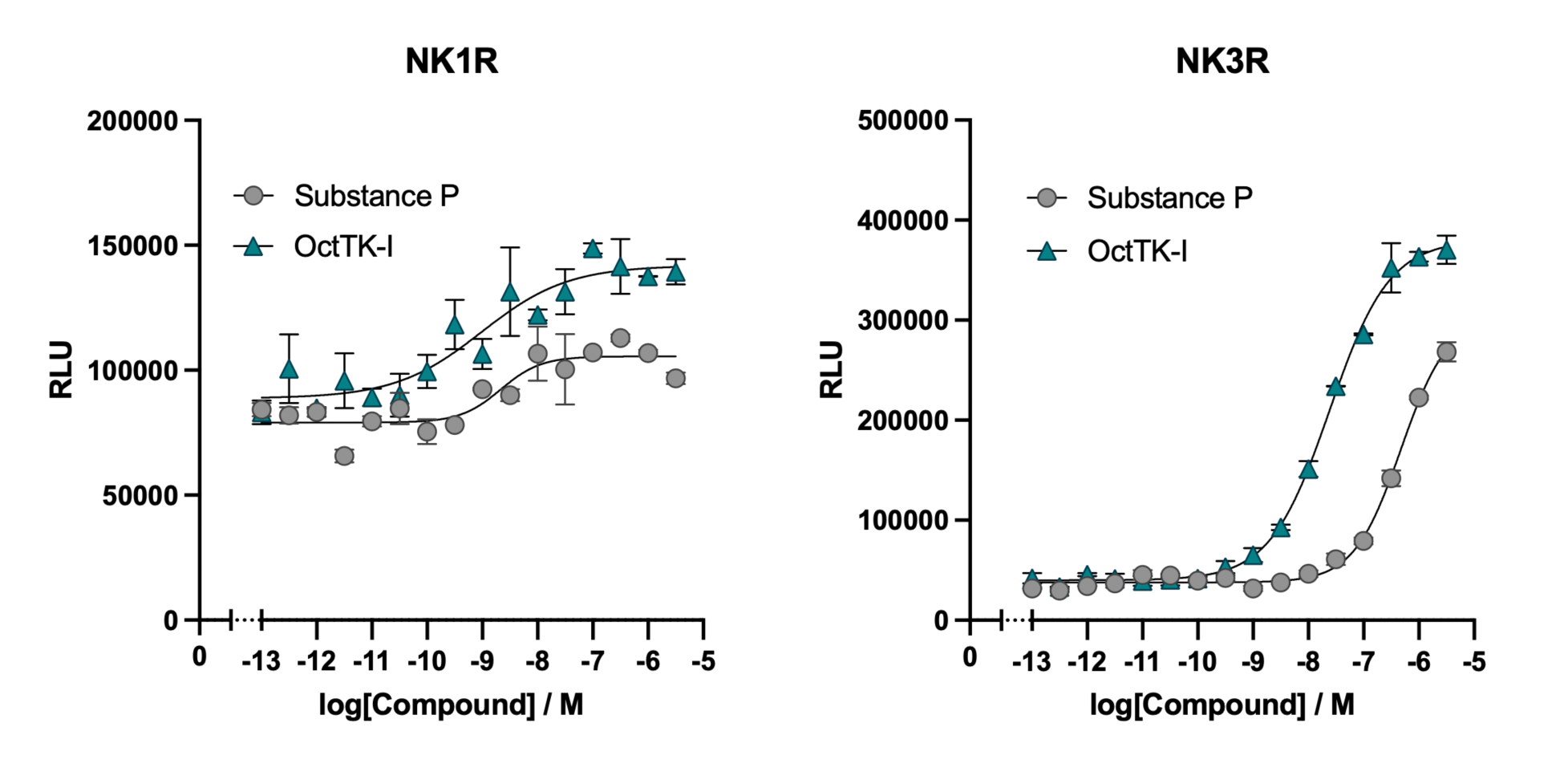
Octopus tachykinin I from *O. vulgaris* (OctTK-I) is a potent activator of the human neurokinin 1 (NK1R) and 3 (NK3R) receptors. Representative dose-response curves of the activation of NKR1 and NKR3 by the endogenous human ligand Substance P and OctTK-I measured using the PRESTO-Tango β-arrestin recruitment assay. EC_50_ of OctTK-I and Substance P at NK1R: 7.23 ± 5.52 nM and 11.64 ± 7.8 nM, respectively. EC_50_ of OctTK-I and Substance P at NK3R: 34.5 ± 8.3 nM and 312.2 ± 59.0 nM, respectively. Data represent means ± S.E.M. of 5 biological replicates with technical duplicates. The human neurokinin 2 (NK2R) receptor was not tested due to technical difficulties.

**Fig. S10.**
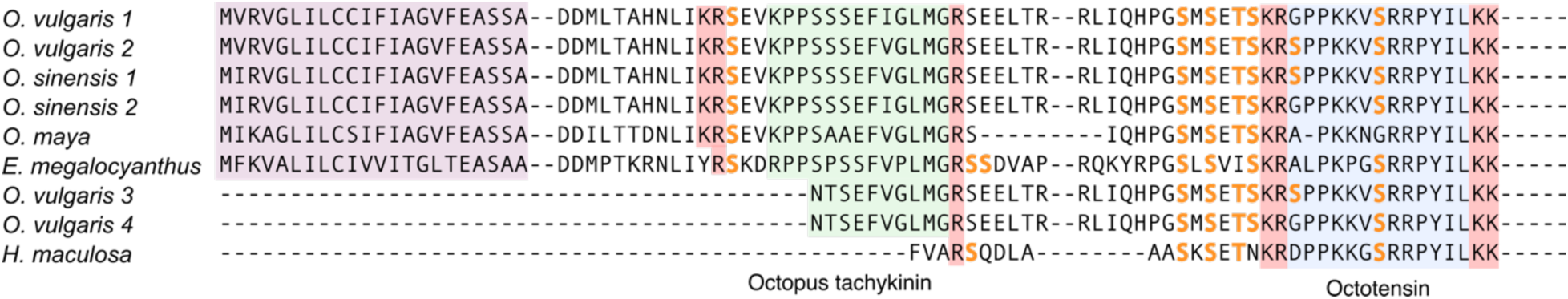
Predicted O-glycosylation of octotensin-precursors. NetOGlyc 4.0 analysis of potential O-glycosylated Ser/Thr residues in octotensin-encoding SF1 precursors. The bolded serine and threonine residues are predicted to be glycosylated (score above 0.5). In all cases, the serine residues of octotensin are predicted to be glycosylated with high confidence. Octotensins are highlighted in blue and octopus tachykinin-like toxins are highlighted in green. Red represents cleavage sites. Signal peptides are purple.

**Fig. S11.**
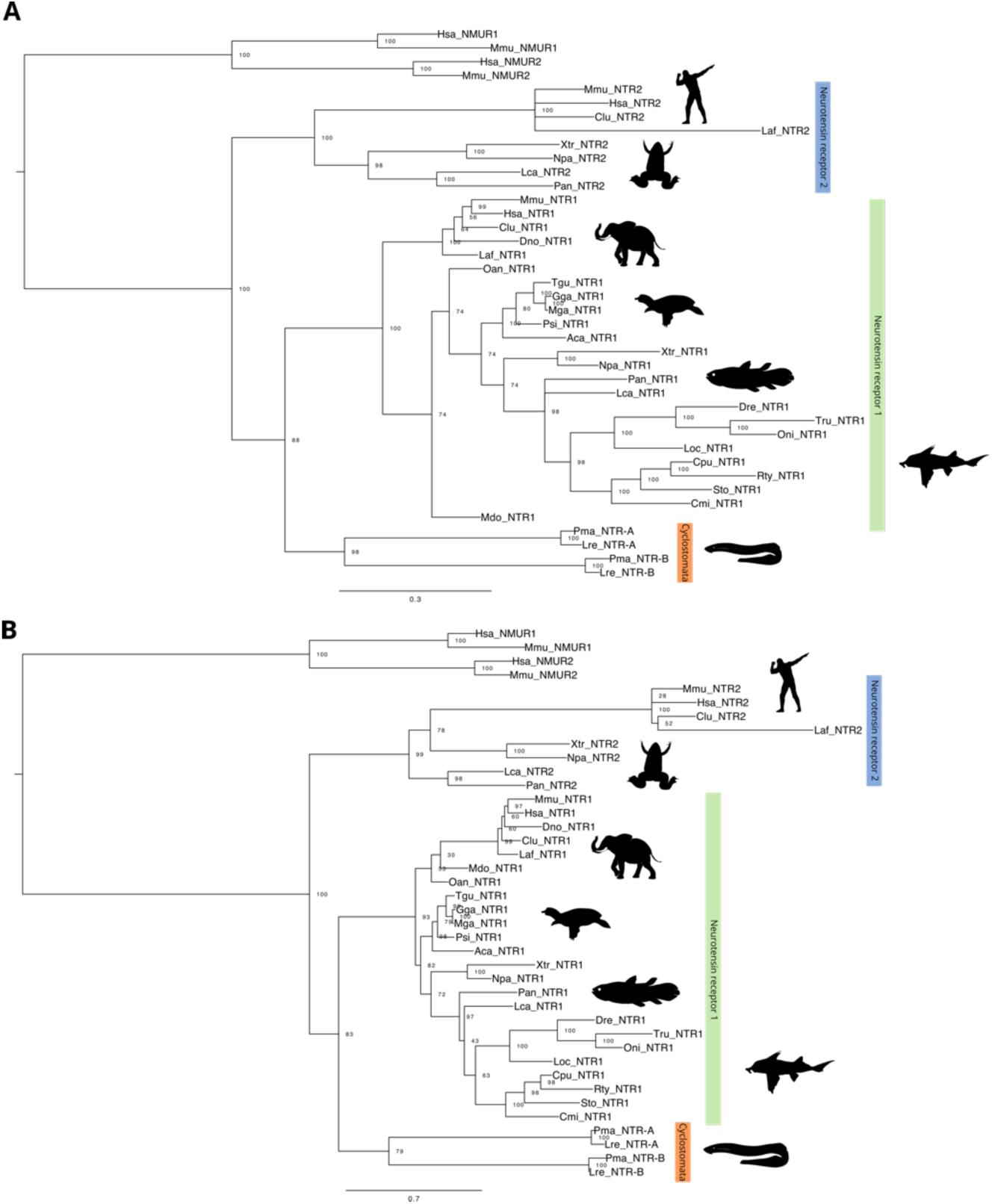
Neurotensin receptor gene trees. **(A)** Bayesian and reconstruction of vertebrate neurotensin receptors. Branch values represent the posterior probability. (B) Maximum likelihood phylogenetic reconstruction of vertebrate neurotensin receptors. Both trees were rooted with neuromedin N receptors. The overall topology of the two trees is highly similar, with the major differences being the position of the monotreme and marsupial NTSR1 genes, which groups with other mammals in the maximum likelihood tree, but not in the Bayesian tree. The trees show clear groupings of gnathostome NTSR1 and NTSR2 branches, as well as a separate cyclostome branch grouping most closely with NTSR1 genes. Hsa: *Homo sapiens,* Mmu: *Mus musculus*, Clu: *Canis lupus familiaris*, Laf: *Loxodonta africana*, Xtr: *Xenopus tropicalis*, Npa: *Nanorana parkeri*, Lca: *Latimeria chalumnae*, Pan: *Protopterus annectens*; Dno: *Dasypus novemcinctus*, Mdo: *Monodelphis domestica*, Oan: *Ornithorhynchus anatinus*, Tgu: *Taenopygia guttata*, Gga*: Gallus gallus*, Mga: *Meleagris gallopavo*, Psi: *Pelodiscus sinensis*, Dre: *Danio rerio*, Tru: *Takifugu rubripes*, Oni: *Oreochrmis niloticus*, Loc: *Lepisosteus oculantus*, Cpu: *Chiloscyllium punctatum*, Rty: *Rhincodon typus*, Sto: *Scyliorhinus torazame*, Cmi: *Callorhinchus milii*, Pma: *Petromyzon marinus*, Lre: *Lethenteron reissneri*. Animal silhouettes were downloaded from https://phylopic.org/, with credits to Sarah Werning for xenopus (https://creativecommons.org/licenses/by/3.0/), Agnello Picorelli for elephant (https://creativecommons.org/licenses/by-nc-sa/3.0/), and Soledad Miranda-Rottmann for the pelodiscus (https://creativecommons.org/licenses/by/3.0/).

**Fig. S12.**
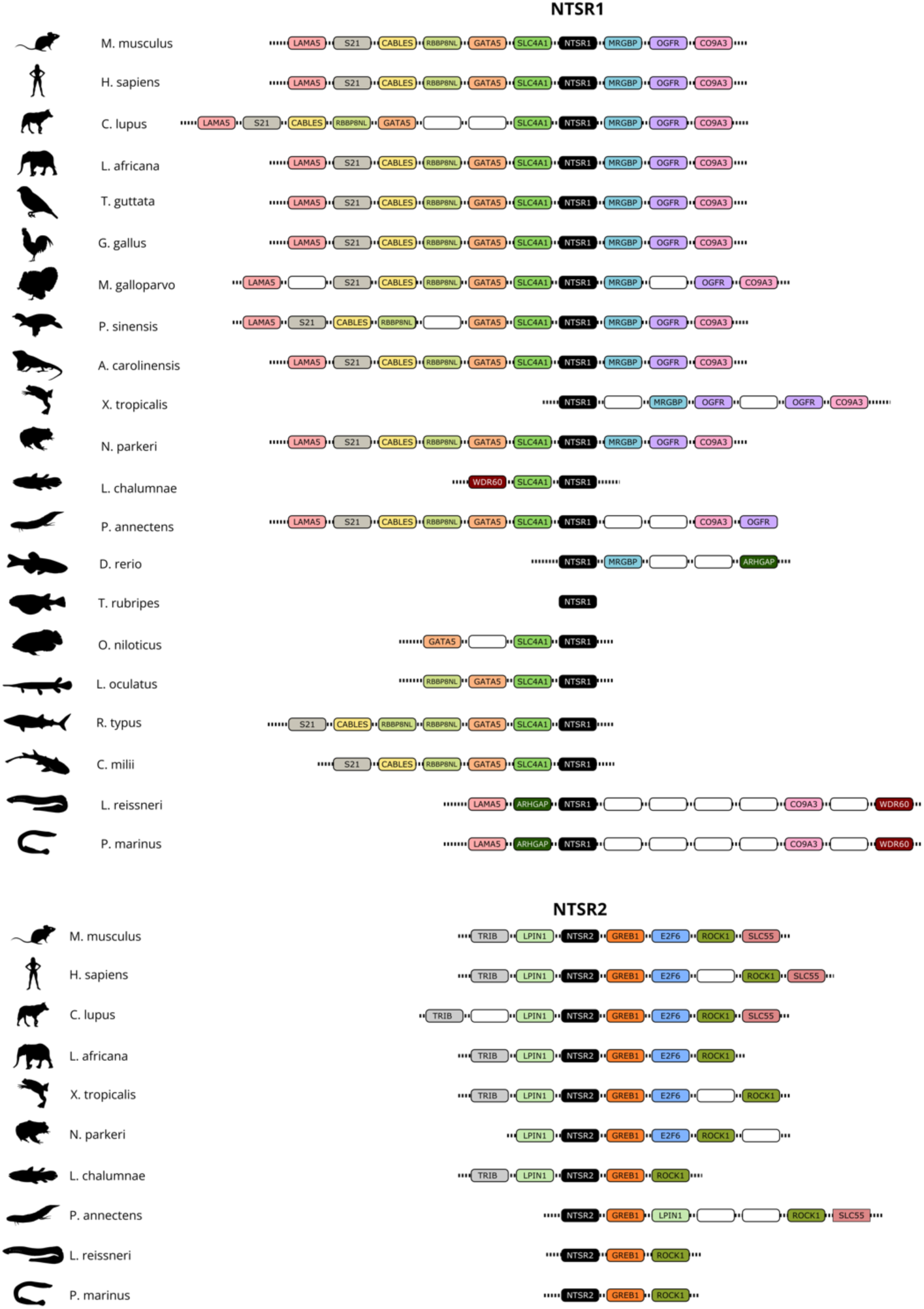
Syntenic schema of regions surrounding NTSR genes derived from genomic analysis. Both NTSR1 and NTSR2 genes are encoded in genomic regions with significant collinearity and strong local gene synteny with few exceptions. The gene synteny analysis reveals that the two cyclostome NTSR genes can be classified into NTSR1 and NTSR2 genes. Gene names are from annotations. Blank boxes represent genes with other annotations. Animal silhouettes were downloaded from https://phylopic.org/, with credits to Sarah Werning for the human (https://creativecommons.org/licenses/by/3.0/), and Soledad Miranda-Rottmann for pelodiscus (https://creativecommons.org/licenses/by/3.0/).

**Fig. S13.**
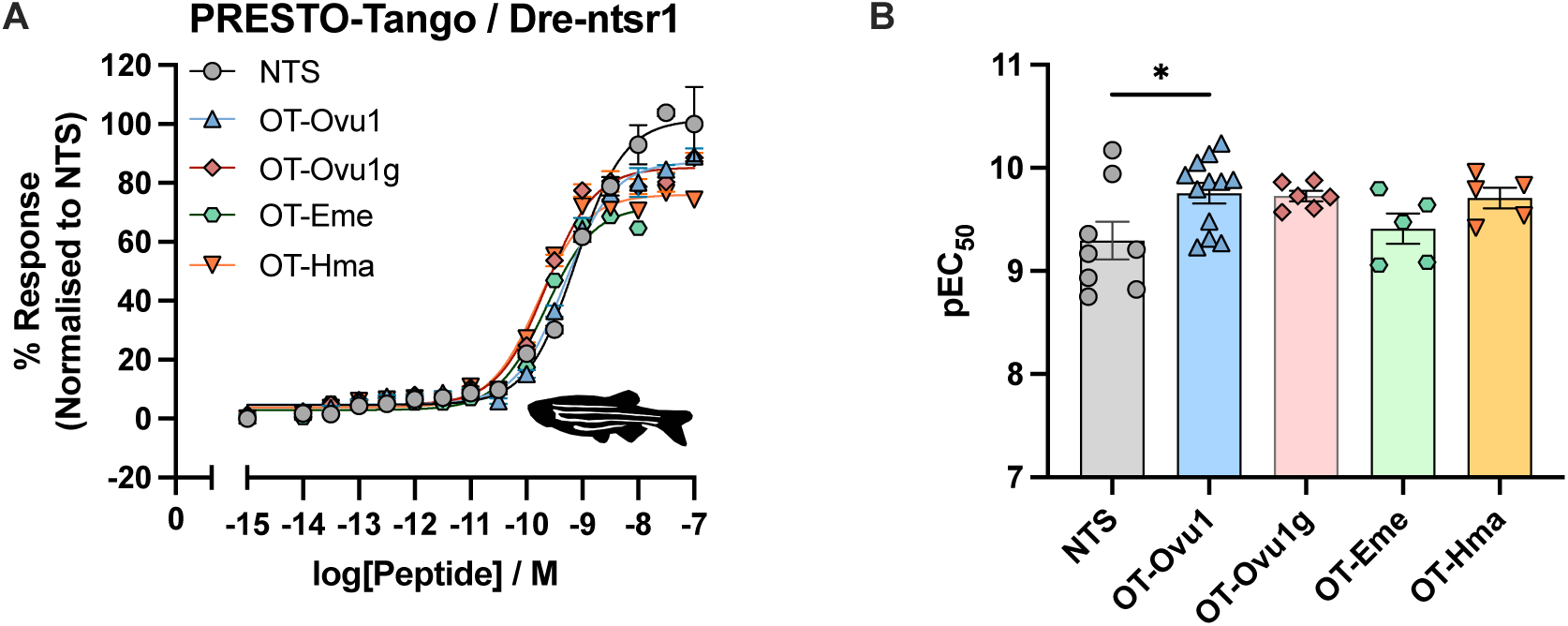
Octotensins identified from *O. vulgaris, H. maculosa and E. megalocyathus* activate the zebrafish NTSR1 (Dre-ntsr1). (A) Dose-response curves of the activation of Dre-ntsr1 by neurotensin, OT-Ovu1, OT-Ovu1g, OT-Hma and OT-Eme measured by PRESTO-Tango β-arrestin recruitment assay. Error bars represent means ± S.E.M. of 3-4 biological replicates in technical duplicates. (B) show the scattered plots of the potencies (pEC_50_) of NTS, OT-Ovu1, OT-Ovu1g, OT-Eme and OT-Hma at the Dre-ntsr1. Statistical significance of octotensins in comparison to human NTS were determined by one-way ANOVA with Dunnett’s multiple comparison test (* = p < 0.05). Pharmacological parameters of Dre-ntsr1 activation by different peptide ligands are detailed in Table S4.

**Fig. S14.**
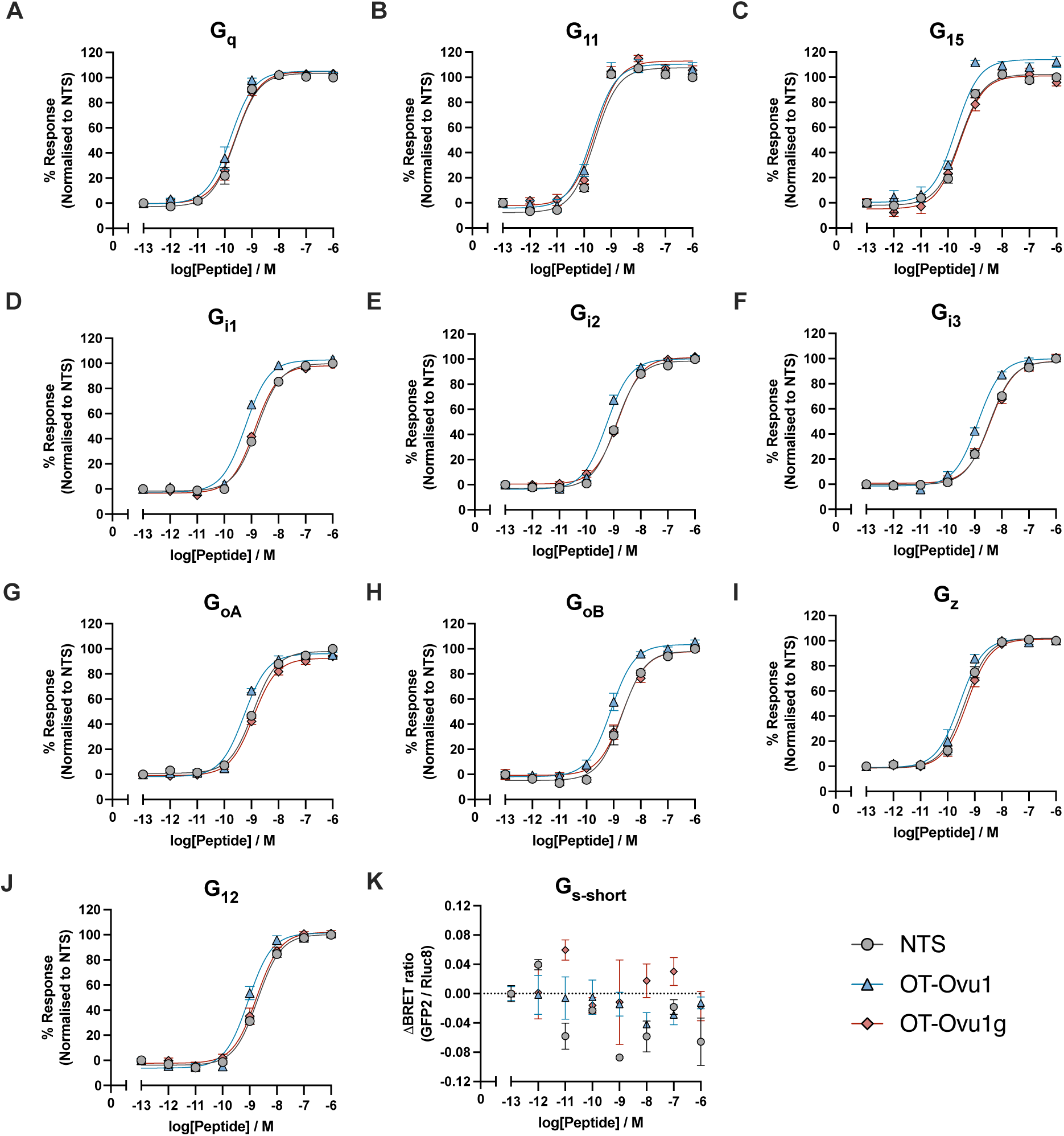
G protein dissociation mediated by NTS, non-glycosylated and glycosylated OT-Ovu1. (A - J) show the representative dose-response curves of NTS, OT-Ovu1 and OT-Ovu1g in mediating G proteins from G_q/11_, G_i/o_ and G_12_ families respectively. (K) show the changes in BRET ratio when NTS, OT-Ovu1 and OT-Ovu1g were tested for Gs-short dissociation. Dose response curves were not determined due to a lack of activation. All error bars show the represent means ± S.E.M. of 3 biological replicates with technical duplicates. Pharmacological parameters are detailed in Table S8.

**Fig. S15.**
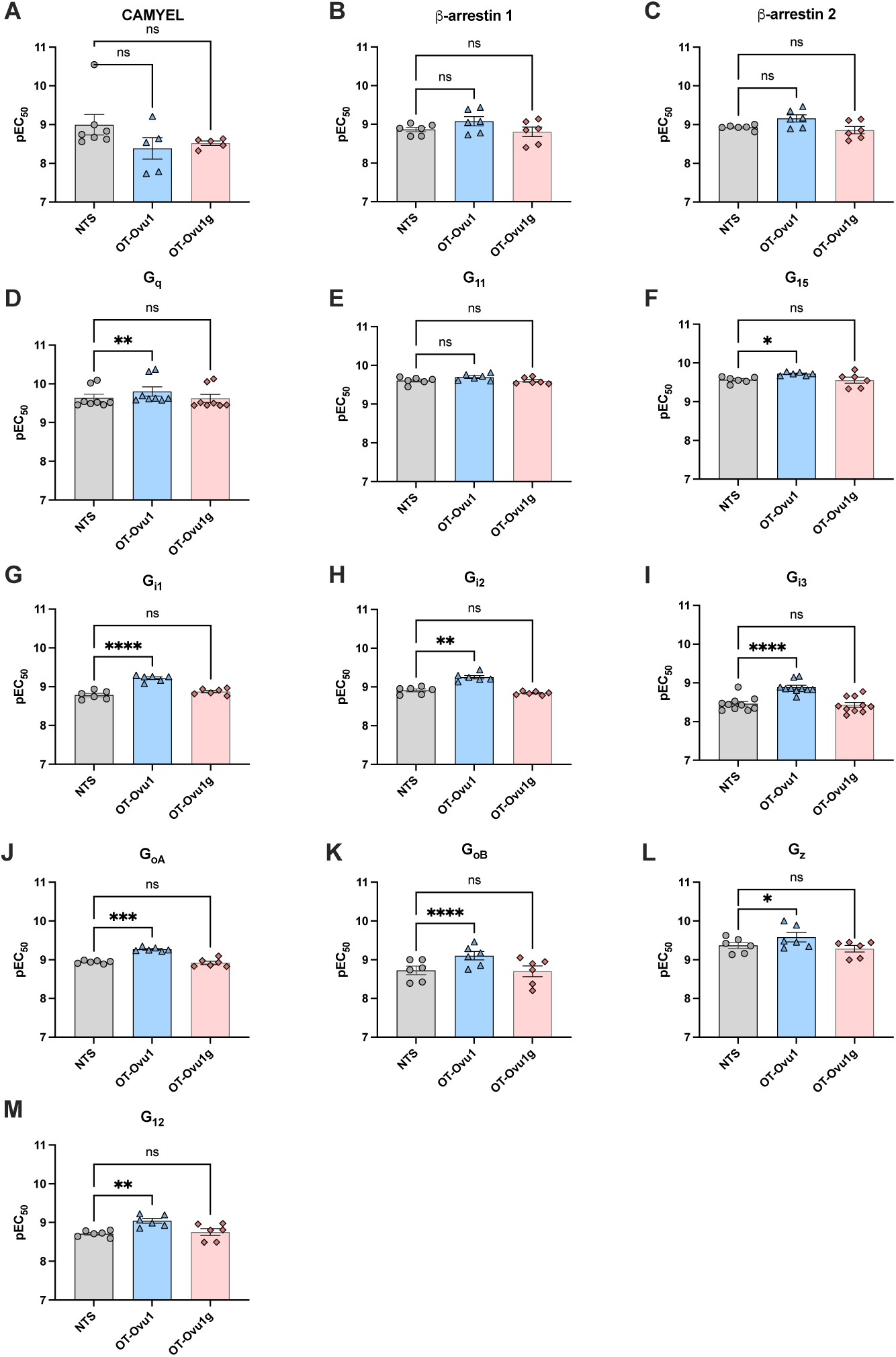
Scattered plots showing the potencies of neurotensin and non-glycosylated and glycosylated OT-Ovu1 at cAMP, β-arrestin recruitment, G protein dissociation assays. Scattered plots of the potencies (pEC_50_) of NTS, OT-Ovu1 and OT-Ovu1g at mediating (A) cAMP, and (B, C) β-arrestin 1/2 recruitment the human NTSR1. (D-M) Scattered plots of pEC_50_ values of NTS, OT-Ovu1 and OT-Ovu1g at dissociating different G proteins at the human NTSR1. Statistical significance of octotensins in comparison to human NTS were determined by one-way ANOVA with Dunnett’s multiple comparison test (* = P < 0.05; ** = P < 0.01; *** = P < 0.001; **** = P < 0.0001). Pharmacological parameters of G protein dissociation mediated by different peptide ligands are detailed in Tables S6 and 8.

**Fig. S16.**
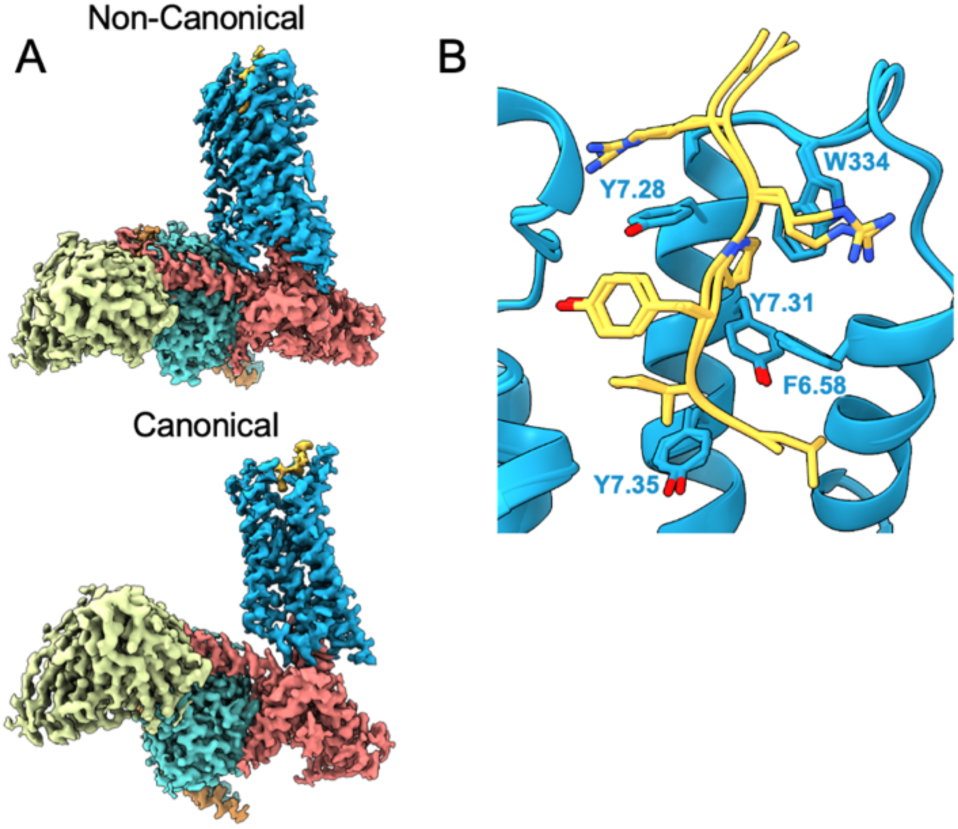
A) Cryo-EM maps for the non-canonical and canonical conformations of the OT-Ovu1g NTSR1 Gi3 complex. B) Overlay of the structures of the core octotensin OT-Ovu1g residues for the non-canonical and canonical conformation structures.

**Fig. S17.**
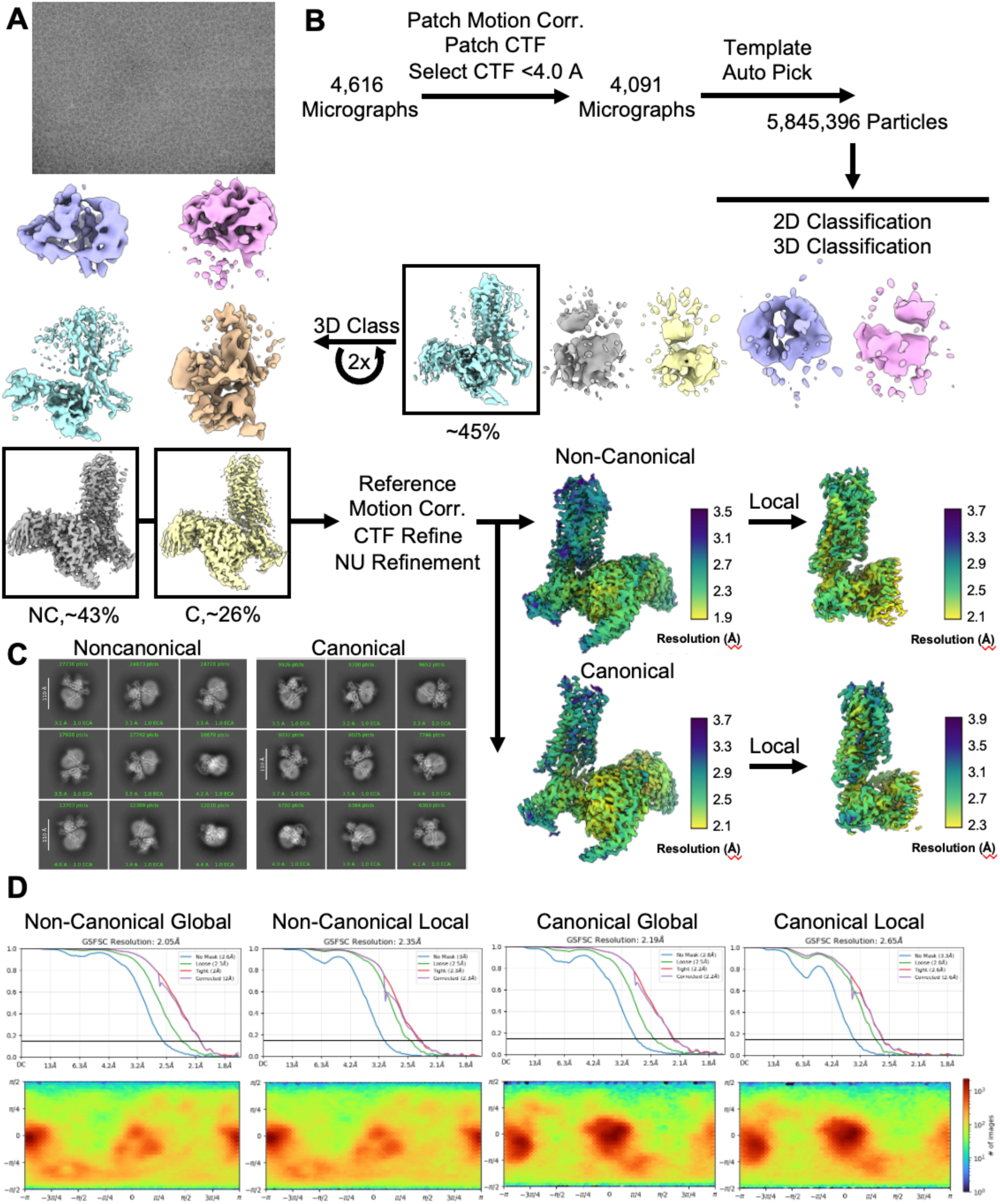
A) Representative cryo-EM micrograph of NTSR1 complex B) Cryo-EM processing workflow employed in this study including final maps with local resolution. C) 2D Class averages of the particles contributing to the two final structures in this work. D) FSC plots and Euler angle distributions for the maps derived in this work.

**Fig. S18.**
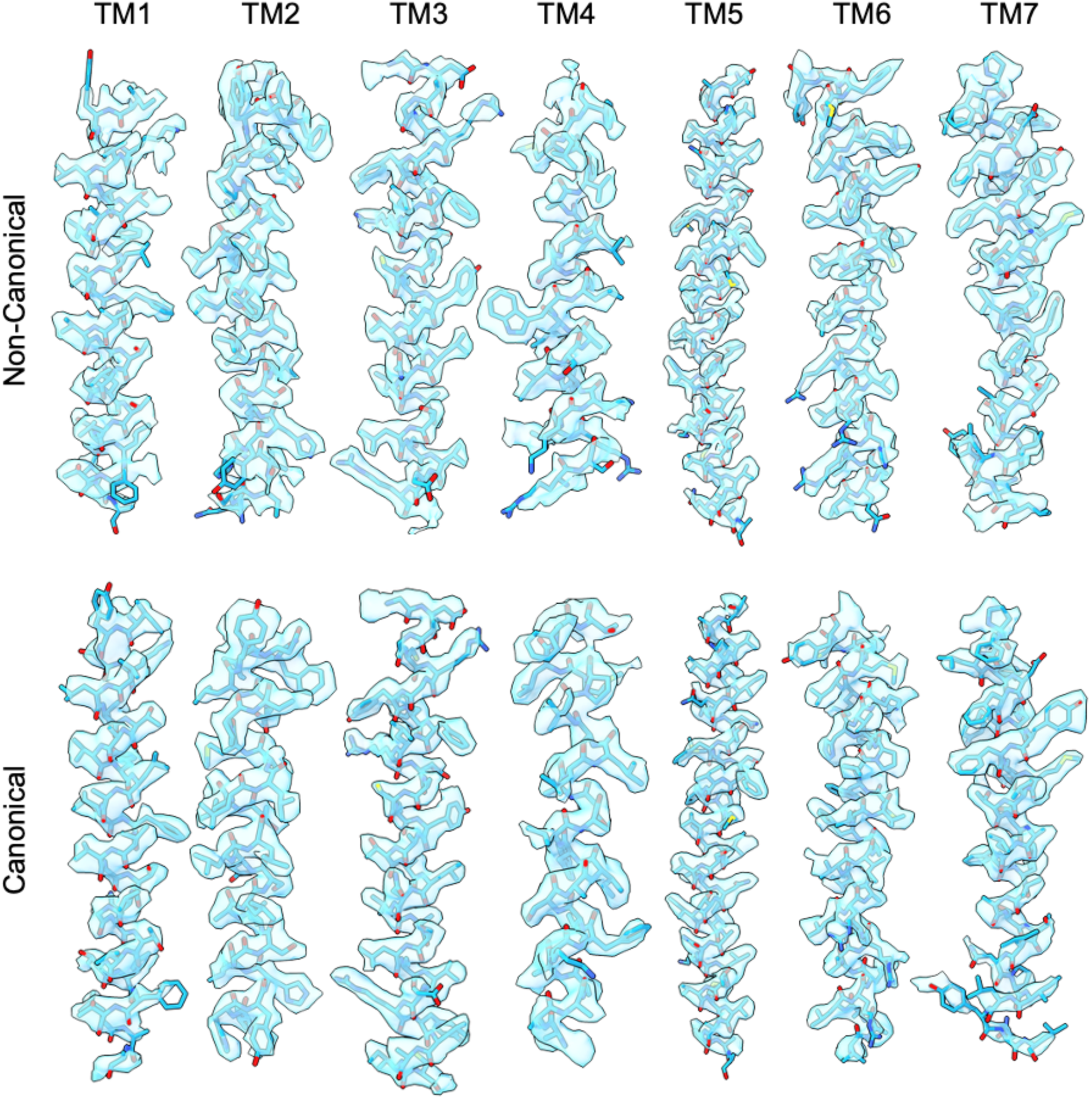
A) Map-Model agreement for the NTSR1 transmembrane helices and the local refined receptor maps for the canonical and non-canonical states resolved in this work.

**Fig. S19.**
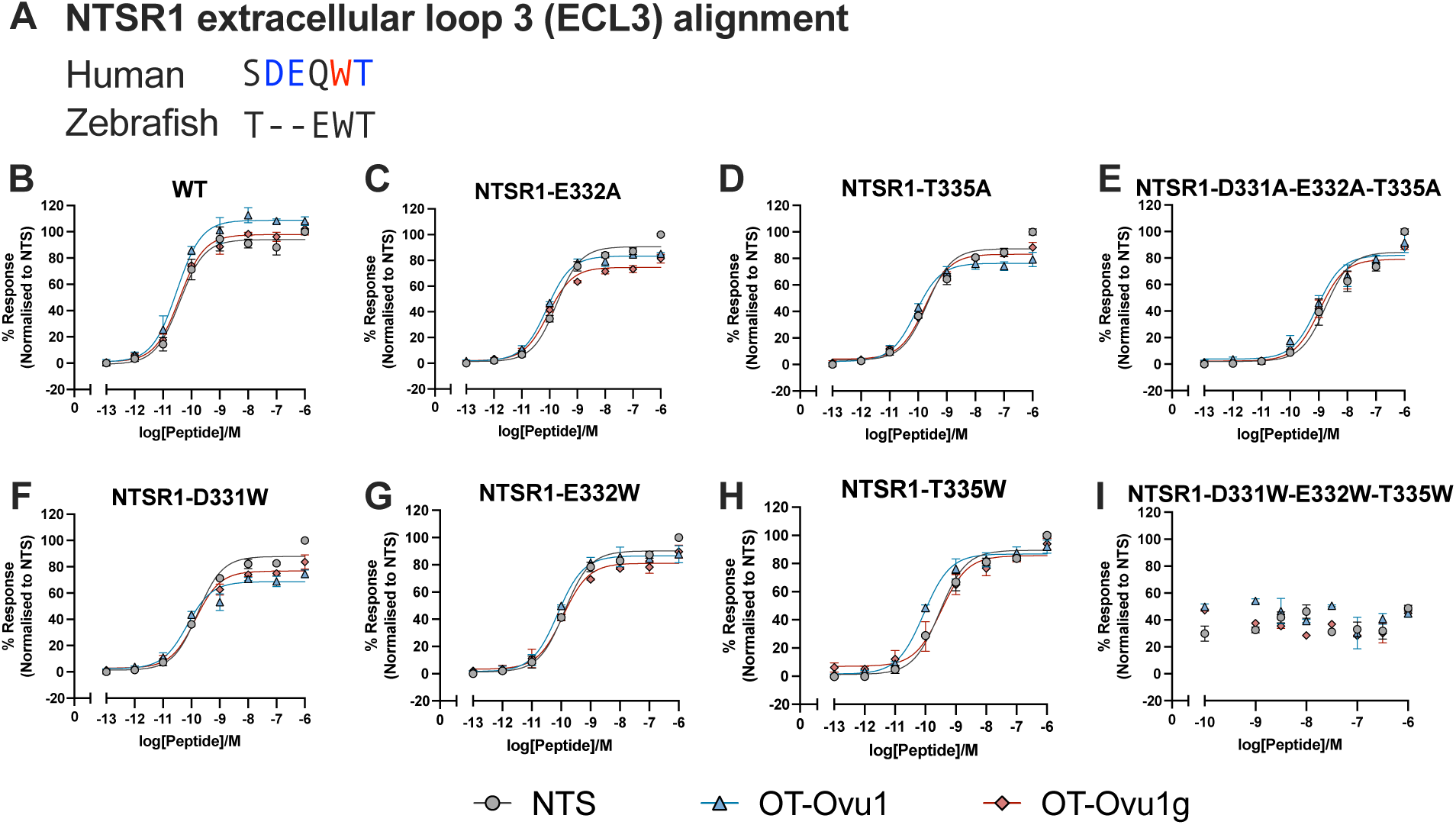
Single mutations at hNTSR1 extracellular loop 3 (ECL3) did not lead to significant loss in IP accumulation mediated by NTS, non-glycosylated and glycosylated OT-Ovu1. (A) shows the sequence alignment of hNTSR1 and Dre-ntsr1 ECL3. Single mutants (highlighted in blue) did not lead to significant reduction in NTSR1 signaling, while W334 mutation (highlighted in red) led to significant potency reduction (See Fig. 3C). (B - I) show the representative dose-response curves of NTS, OT-Ovu1 and OT-Ovu1g in IP accumulation tested in different NTSR1 mutations transiently transfected in HEK239T cells. All error bars show the represent means ± S.E.M. of at least 2 biological replicates with technical duplicates.

**Fig. S20.**
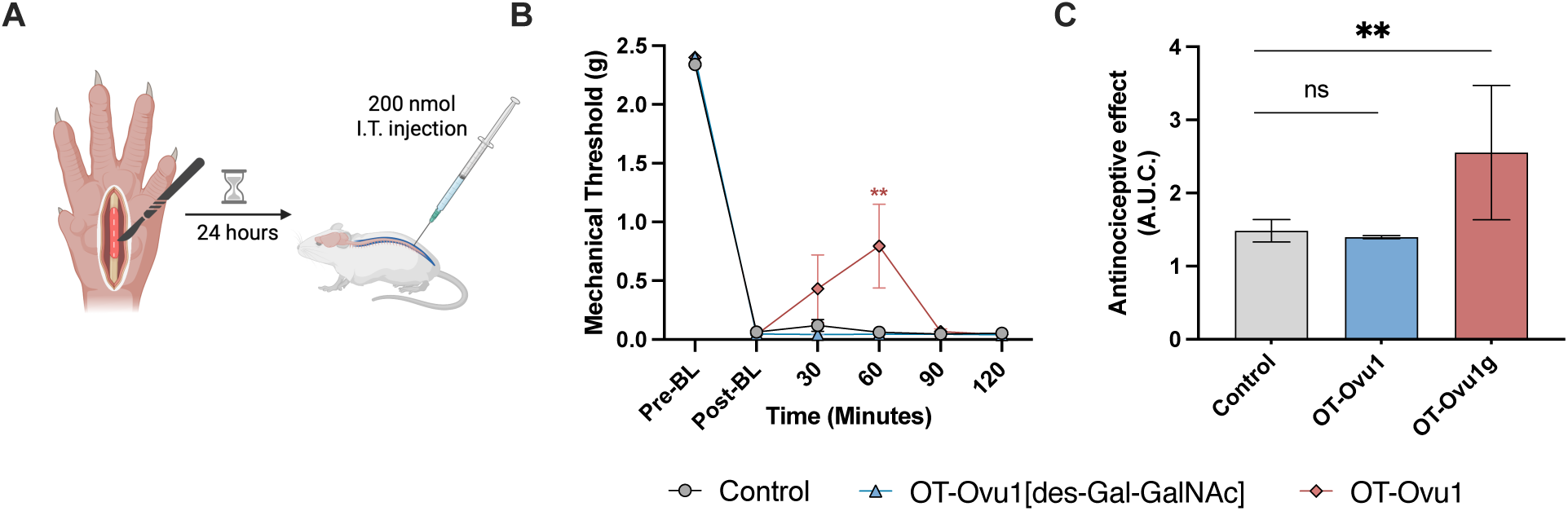
OT-Ovu1g but not OT-Ovu1 produces central analgesia albeit only at a high dose. (A) Schematic of post-operative paw incision pain model. OT-Ovu1g, OT-Ovu1 or vehicle were administered intrathecally (i.t.) (200 nmol/5μL). (B) Mechanical threshold was measured over time. Two-way ANOVA with Tukey’s test; means ± SEM; ** = P < 0.01, n = 8. (C) Area under the curve (A.U.C., baseline- 2 h) showing antinociceptive effects; mean ± SEM; Ordinary one-way ANOVA, multiple comparisons test, n = 8. ** = P < 0.01, ns = not significant. Mice treated with OT-Ovu1 and OT-Ovu1g showed symptoms of transient locomotor impairment.

**Fig. S21.**
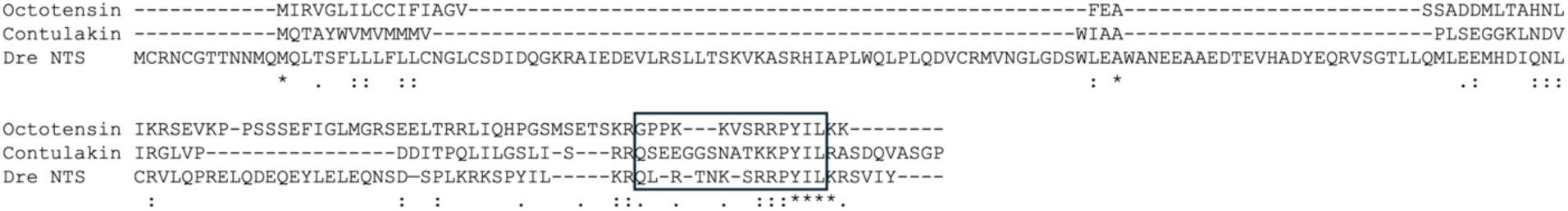
Alignment of the precursor sequences of octotensin Ovu-1, contulakin-G, and zebrafish neurotensin. Protein sequence alignment shows no similarity across the three precursors besides the peptide-encoding region of the C-terminus (boxed). Contulakin-G evolved from the C-superfamily of conotoxins that originally evolved through the recruitment of a somatostatin-like peptide-encoding gene (*19*) while OT-Ovu1evolved *de novo* from a gene region within the SF1 superfamily, demonstrating their independent convergent evolution.

**Table S1.** Transcriptome datasets of the anterior salivary gland (ASG) and posterior salivary gland (PSG) used in this paper.

| Species | Tissue | SRR | URL |
| --- | --- | --- | --- |
| *Haplochlana maculosa* | PSG | SRR11782480 | https://www.ncbi.nlm.nih.gov/sra/SRR11782480 |
| *Haplochlana maculosa* | PSG | SRR11782486 | https://www.ncbi.nlm.nih.gov/sra/SRR11782486 |
| *Haplochlana maculosa* | ASG | SRR11782488 | https://www.ncbi.nlm.nih.gov/sra/SRR11782488 |
| *Doryteuthis pealeii* | PSG | SRR18071813 | https://www.ncbi.nlm.nih.gov/sra/SRR18071813 |
| *Octopus bimaculoides* | PSG | SRR2047107 | https://www.ncbi.nlm.nih.gov/sra/SRR2047107 |
| *Octopus kaurana* | PSG | SRR3105321 | https://www.ncbi.nlm.nih.gov/sra/SRR3105321 |
| *Haplochlana maculosa* | PSG | SRR3105558 | https://www.ncbi.nlm.nih.gov/sra/SRR3105558 |
| *Haplochlana maculosa* | ASG | SRR3105561 | https://www.ncbi.nlm.nih.gov/sra/SRR3105561 |
| *Sepia officinalis* | PSG | SRR5204441 | https://www.ncbi.nlm.nih.gov/sra/SRR5204441 |
| *Sepia officinalis* | PSG | SRR5204442 | https://www.ncbi.nlm.nih.gov/sra/SRR5204442 |
| *Octopus minor* | PSG | SRR6349992 | https://www.ncbi.nlm.nih.gov/sra/SRR6349992 |
| *Octopus vulgaris* | PSG | SRR7130741 | https://www.ncbi.nlm.nih.gov/sra/SRR7130741 |
| *Octopus bimaculoides* | ASG | SRR7645290 | https://www.ncbi.nlm.nih.gov/sra/SRR7645290 |
| *Octopus bimaculoides* | ASG | SRR7645291 | https://www.ncbi.nlm.nih.gov/sra/SRR7645291 |
| *Octopus bimaculoides* | ASG | SRR7645293 | https://www.ncbi.nlm.nih.gov/sra/SRR7645293 |
| *Octopus bimaculoides* | ASG | SRR7645334 | https://www.ncbi.nlm.nih.gov/sra/SRR7645334 |
| *Octopus bimaculoides* | ASG | SRR7645337 | https://www.ncbi.nlm.nih.gov/sra/SRR7645337 |
| *Octopus bimaculoides* | PSG | SRR7645354 | https://www.ncbi.nlm.nih.gov/sra/SRR7645354 |
| *Octopus bimaculoides* | PSG | SRR7645355 | https://www.ncbi.nlm.nih.gov/sra/SRR7645355 |
| *Octopus bimaculoides* | PSG | SRR7645356 | https://www.ncbi.nlm.nih.gov/sra/SRR7645356 |
| *Octopus bimaculoides* | PSG | SRR7645357 | https://www.ncbi.nlm.nih.gov/sra/SRR7645357 |
| *Octopus bimaculoides* | PSG | SRR7645510 | https://www.ncbi.nlm.nih.gov/sra/SRR7645510 |
| *Octopus bimaculoides* | PSG | SRR7645511 | https://www.ncbi.nlm.nih.gov/sra/SRR7645511 |
| *Octopus bimaculoides* | PSG | SRR7645512 | https://www.ncbi.nlm.nih.gov/sra/SRR7645512 |
| *Octopus bimaculoides* | PSG | SRR7645513 | https://www.ncbi.nlm.nih.gov/sra/SRR7645513 |

**Table S2.**
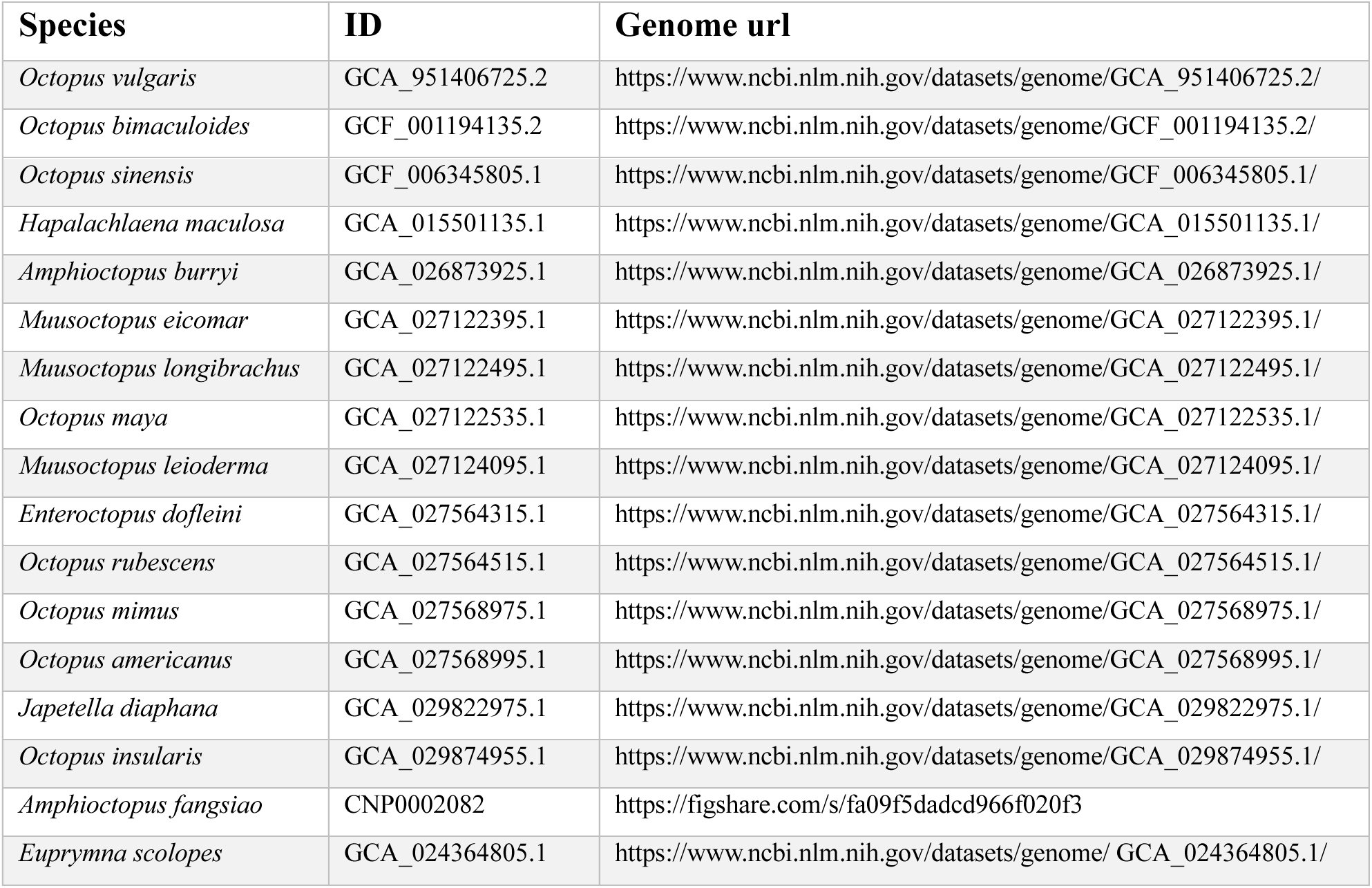
Genomes used in this paper.

**Table S3.** Annotated enzymes from posterior salivary gland transcriptomes of octopuses required for Gal-GalNAc modification. For each species the values show the highest e-value of identified enzymes belonging to the associated PANTHER and FunFam family with GALT (PTHR11675, FunFam G3DSA:3.90.550.10:FF:000005) and C1GLT (PTHR23033, FunFam G3DSA:3.90.550.50:FF:000017).

|  | <u>*H. maculosa*</u> |  | <u>*O. vulgaris*</u> |  | <u>*O. bimaculoides*</u> |  | <u>*O. kaurana*</u> |  | <u>*O. minor*</u> |  |
| --- | --- | --- | --- | --- | --- | --- | --- | --- | --- | --- |
|  | PTHR | FunFam | PTHR | FunFam | PTHR | FunFam | PTHR | FunFam | PTHR | FunFam |
| GALT<sup>a</sup> | 2.3E-245 | 1.2E-208 | 1.2E-51 | - | 9.5E-203 | 4.6E-209 | 3.8E-203 | 5.1E-207 | 8.5E-206 | 5.1E-207 |
| C1GLT<sup>b</sup> | 9.4E-214 | 1.7E-172 | 2.6E-83 | 3.2E-87 | 4.4E-91 | 5.0E-95 | 6.4E-49 | - | 1.9E-81 | 1.8E-87 |

<sup>a</sup>Polypeptide N-acetylgalactosaminyltransferase
<sup>b</sup>Glycoprotein-N-acetylgalactosamine 3-beta-galactosyltransferase

**Table S4.** Pharmacological parameters of NTS, OT-Ovu1, OT-Ovu1pg, OT-Ovu1g, OT-Eme and OT-Hma at the Dre-NTSR1 measured in IPOne Gq activation assay.

| Test ligands | Parameters | Values |
| --- | --- | --- |
| **NTS** | pEC<sub>50</sub><sup>a</sup> ± S.E.M. | 9.488 ± 0.119 |
|  | EC<sub>50</sub> (nM) | 0.325 |
|  | E<sub>max</sub><sup>b</sup> ± S.E.M. | 99.55 ± 3.26 |
|  | Span ± S.E.M. | 95.85 ± 4.52 |
|  | n | 6 |
| **OT-Ovu1** | pEC<sub>50</sub><sup>a</sup> ± S.E.M. | 9.760 ± 0.108 |
|  | EC<sub>50</sub> (nM) | 0.174 |
|  | E<sub>max</sub><sup>b</sup> ± S.E.M. | 102.03 ± 3.20 |
|  | Span ± S.E.M. | 103.38 ± 4.58 |
|  | n | 6 |
| **OT-Ovu1pg** | pEC<sub>50</sub><sup>a</sup> ± S.E.M. | 9.977 ± 0.193 |
|  | EC<sub>50</sub> (nM) | 0.106 |
|  | E<sub>max</sub><sup>b</sup> ± S.E.M. | 99.67 ± 5.35 |
|  | Span ± S.E.M. | 97.61 ± 8.00 |
|  | n | 6 |
| **OT-Ovu1g** | pEC<sub>50</sub><sup>a</sup> ± S.E.M. | 9.824 ± 0.118 |
|  | EC<sub>50</sub> (nM) | 0.150 |
|  | E<sub>max</sub><sup>b</sup> ± S.E.M. | 76.88 ± 4.33 |
|  | Span ± S.E.M. | 97.31 ± 4.78 |
|  | n | 6 |
| **OT-Eme** | pEC<sub>50</sub><sup>a</sup> ± S.E.M. | 10.046 ± 0.138 |
|  | EC<sub>50</sub> (nM) | 0.0899 |
|  | E<sub>max</sub><sup>b</sup> ± S.E.M. | 86.07 ± 3.37 |
|  | Span ± S.E.M. | 87.14 ± 5.21 |
|  | n | 5 |
| **OT-Hma** | pEC<sub>50</sub><sup>a</sup> ± S.E.M. | 9.825 ± 0.117 |
|  | EC<sub>50</sub> (nM) | 0.150 |
|  | E<sub>max</sub><sup>b</sup> ± S.E.M. | 94.83 ± 3.25 |
|  | Span ± S.E.M. | 93.22 ± 4.66 |
|  | n | 9.825 ± 0.117 |

Values were generated when the data were fitted to the four-parameter logistic equation. Means ± S.E.M of n individual result sets were shown.
<sup>a</sup> Negative logarithm of agonist concentration when reaching half maximal response.
<sup>b</sup> % of maximal response observed when stimulated with ligands relative to human neurotensin.

**Table S5.** Pharmacological parameters of NTS, OT-Ovu1, OT-Ovu1g, OT-Hma and OT-Eme at the Dre-ntsr1 measured by the PRESTO-Tango β-arrestin recruitment assay.

| Test ligands | Parameters | Values |
| --- | --- | --- |
| **NTS** | pEC<sub>50</sub><sup>a</sup> ± S.E.M. | 9.118 ± 0.055 |
|  | EC<sub>50</sub> (nM) | 0.762 |
| | E<sub>max</sub><sup>b</sup> ± S.E.M. | 101.57 ± 2.24 |
| | Span ± S.E.M. | 96.87 ± 2.45 |
|  | n | 8 |
| **OT-Ovu1** | pEC<sub>50</sub><sup>a</sup> ± S.E.M. | 9.313 ± 0.040 |
|  | EC<sub>50</sub> (nM) | 0.487 |
| | E<sub>max</sub><sup>b</sup> ± S.E.M. | 87.14 ± 1.32 |
| | Span ± S.E.M. | 82.32 ± 1.47 |
|  | n | 8 |
| **OT-Ovu1g** | pEC<sub>50</sub><sup>a</sup> ± S.E.M. | 9.648 ± 0.042 |
|  | EC<sub>50</sub> (nM) | 0.225 |
| | E<sub>max</sub><sup>b</sup> ± S.E.M. | 85.24 ± 1.24 |
| | Span ± S.E.M. | 80.53 ± 1.43 |
|  | n | 8 |
| **OT-Eme** | pEC<sub>50</sub><sup>a</sup> ± S.E.M. | 9.686 ± 0.041 |
|  | EC<sub>50</sub> (nM) | 0.206 |
| | E<sub>max</sub><sup>b</sup> ± S.E.M. | 67.60 ± 1.75 |
| | Span ± S.E.M. | 63.34 ± 2.02 |
| | Hill Slope | 1.776 ± 0.247 |
|  | n | 5 |
| **OT-Hma** | pEC<sub>50</sub><sup>a</sup> ± S.E.M. | 9.791 ± 0.044 |
|  | EC<sub>50</sub> (nM) | 0.162 |
| | E<sub>max</sub><sup>b</sup> ± S.E.M. | 73.99 ± 1.43 |
| | Span ± S.E.M. | 69.01 ± 1.804 |
| | Hill Slope | 1.516 ± 0.20 |
|  | n | 5 |

Values were generated when the data were fitted to the four-parameter logistic equation. Means ± S.E.M of n individual result sets were shown.
<sup>a</sup> Negative logarithm of agonist concentration when reaching half maximal response.
<sup>b</sup> % of maximal response observed when stimulated with ligands relative to human neurotensin.

**Table S6.** Pharmacological parameters of NTS, OT-Ovu1, OT-Ovu1g, OT-Eme and OT-Hma at the human NTSR1 measured in IPOne Gq activation assay.

| Test ligands | Parameters | Values |
| --- | --- | --- |
| **NTS** | pEC<sub>50</sub><sup>a</sup> ± S.E.M. | 9.089 ± 0.071 |
|  | EC<sub>50</sub> (nM) | 0.815 |
|  | E<sub>max</sub><sup>b</sup> ± S.E.M. | 95.51 ± 2.27 |
|  | Span ± S.E.M. | 97.52 ± 2.91 |
|  | n | 6 |
| **OT-Ovu1** | pEC<sub>50</sub><sup>a</sup> ± S.E.M. | 9.453 ± 0.095 |
|  | EC<sub>50</sub> (nM) | 0.353 |
|  | E<sub>max</sub><sup>b</sup> ± S.E.M. | 106.21 ± 3.11 |
|  | Span ± S.E.M. | 110.23 ± 4.11 |
|  | n | 6 |
| **OT-Ovu1pg** | pEC<sub>50</sub><sup>a</sup> ± S.E.M. | 9.464 ± 0.108 |
|  | EC<sub>50</sub> (nM) | 0.343 |
|  | E<sub>max</sub><sup>b</sup> ± S.E.M. | 109.85 ± 3.60 |
|  | Span ± S.E.M. | 108.26 ± 4.68 |
|  | n | 6 |
| **OT-Ovu1g** | pEC<sub>50</sub><sup>a</sup> ± S.E.M. | 9.222 ± 0.091 |
|  | EC<sub>50</sub> (nM) | 0.600 |
|  | E<sub>max</sub><sup>b</sup> ± S.E.M. | 105.12 ± 3.09 |
|  | Span ± S.E.M. | 105.16 ± 3.96 |
|  | n | 6 |
| **OT-Eme** | pEC<sub>50</sub><sup>a</sup> ± S.E.M. | 9.509 ± 0.098 |
|  | EC<sub>50</sub> (nM) | 0.309 |
|  | E<sub>max</sub><sup>b</sup> ± S.E.M. | 100.19 ± 2.76 |
|  | Span ± S.E.M. | 98.19 ± 3.74 |
|  | n | 6 |
| **OT-Hma** | pEC<sub>50</sub><sup>a</sup> ± S.E.M. | 9.246 ± 0.092 |
|  | EC<sub>50</sub> (nM) | 0.567 |
|  | E<sub>max</sub><sup>b</sup> ± S.E.M. | 102.78 ± 2.98 |
|  | Span ± S.E.M. | 103.06 ± 3.87 |
|  | n | 6 |

Values were generated when the data were fitted to the four-parameter logistic equation. Means ± S.E.M of n individual result sets were shown.
<sup>a</sup> Negative logarithm of agonist concentration when reaching half maximal response.
<sup>b</sup> % of maximal response observed when stimulated with ligands relative to human neurotensin.

**Table S7.** Pharmacological parameters of NTS, OT-Ovu1, OT-Ovu1g, OT-Eme and OT-Hma at the human NTSR1 measured in CAMYEL and β-arrestin recruitment assays.

| Pathway | Parameters | NTS | OT-Ovu1 | OT-Ovu1g |
| --- | --- | --- | --- | --- |
| **cAMP** | pEC<sub>50</sub><sup>a</sup> ± S.E.M. | 8.603 ± 0.102 | 8.396 ± 0.196 | 8.500 ± 0.179 |
|  | EC<sub>50</sub> (nM) | 2.492 | 4.018 | 3.163 |
| | E<sub>max</sub><sup>b</sup> ± S.E.M. | 101.43 ± 3.325 | 98.51 ± 5.897 | 98.91 ± 4.869 |
| | Span ± S.E.M. | 87.51 ± 3.90 | 85.40 ± 7.05 | 79.44 ± 5.92 |
|  | n | 7 | 6 | 5 |
| **β-arrestin 1** | pEC<sub>50</sub><sup>a</sup> ± S.E.M. | 8.859 ± 0.036 | 9.106 ± 0.072 | 8.799 ± 0.069 |
|  | EC<sub>50</sub> (nM) | 1.382 | 0.783 | 1.590 |
| | E<sub>max</sub><sup>b</sup> ± S.E.M. | 103.79 ± 1.30 | 112.39 ± 2.67 | 111.02 ± 2.64 |
| | Span ± S.E.M. | 104.74 ± 1.61 | 113.77 ± 3.44 | 111.78 ± 3.24 |
|  | n | 6 | 6 | 6 |
| **β-arrestin 2** | pEC<sub>50</sub><sup>a</sup> ± S.E.M. | 8.927 ± 0.031 | 9.184 ± 0.077 | 8.858 ± 0.061 |
|  | EC<sub>50</sub> (nM) | 1.183 | 0.657 | 1.388 |
| | E<sub>max</sub><sup>b</sup> ± S.E.M. | 105.30 ± 1.13 | 102.47 ± 2.58 | 106.51 ± 2.20 |
| | Span ± S.E.M. | 106.10 ± 1.41 | 104.33 ± 3.33 | 106.52 ± 2.73 |
|  | n | 6 | 6 | 6 |

Values were generated when the data were fitted to the four-parameter logistic equation. Means ± S.E.M of n individual result sets were shown.
<sup>a</sup> Negative logarithm of agonist concentration when reaching half maximal response.
<sup>b</sup> % of maximal response observed when stimulated with ligands relative to human neurotensin.

**Table S8.**
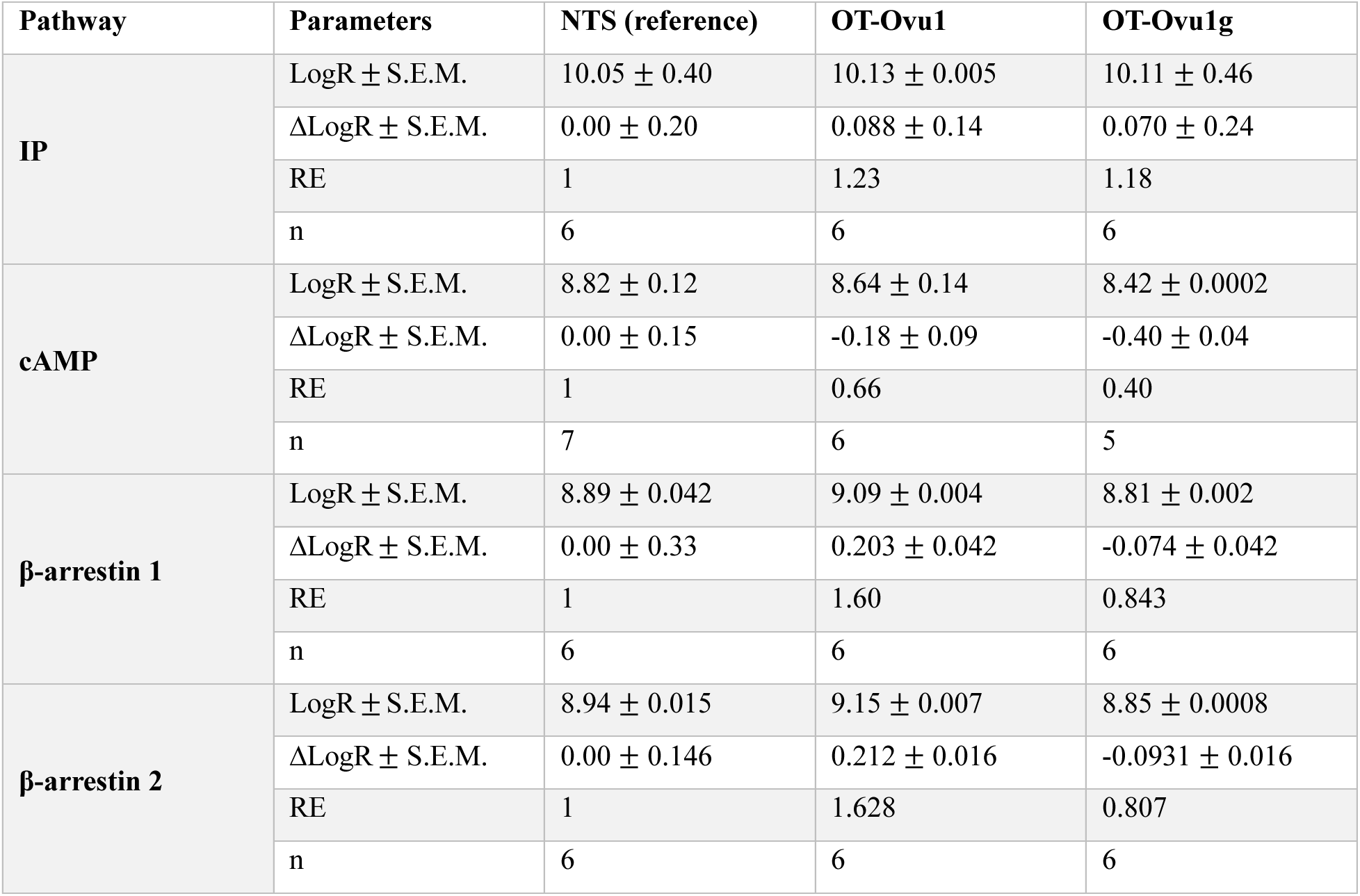
Bias factors of NTS, OT-Ovu1 and OT-Ovu1g mediated IP, cAMP and β-arrestin recruitment at the human NTSR1.

| Pathway | Parameters | NTS (reference) | OT-Ovu1 | OT-Ovu1g |
| --- | --- | --- | --- | --- |
| **IP** | LogR ± S.E.M. | 10.05 ± 0.40 | 10.13 ± 0.005 | 10.11 ± 0.46 |
| | ΔLogR ± S.E.M. | 0.00 ± 0.20 | 0.088 ± 0.14 | 0.070 ± 0.24 |
|  | RE | 1 | 1.23 | 1.18 |
|  | n | 6 | 6 | 6 |
| **cAMP** | LogR ± S.E.M. | 8.82 ± 0.12 | 8.64 ± 0.14 | 8.42 ± 0.0002 |
| | ΔLogR ± S.E.M. | 0.00 ± 0.15 | -0.18 ± 0.09 | -0.40 ± 0.04 |
|  | RE | 1 | 0.66 | 0.40 |
|  | n | 7 | 6 | 5 |
| **β-arrestin 1** | LogR ± S.E.M. | 8.89 ± 0.042 | 9.09 ± 0.004 | 8.81 ± 0.002 |
| | ΔLogR ± S.E.M. | 0.00 ± 0.33 | 0.203 ± 0.042 | -0.074 ± 0.042 |
|  | RE | 1 | 1.60 | 0.843 |
|  | n | 6 | 6 | 6 |
| **β-arrestin 2** | LogR ± S.E.M. | 8.94 ± 0.015 | 9.15 ± 0.007 | 8.85 ± 0.0008 |
| | ΔLogR ± S.E.M. | 0.00 ± 0.146 | 0.212 ± 0.016 | -0.0931 ± 0.016 |
|  | RE | 1 | 1.628 | 0.807 |
|  | n | 6 | 6 | 6 |

**Table S9.**
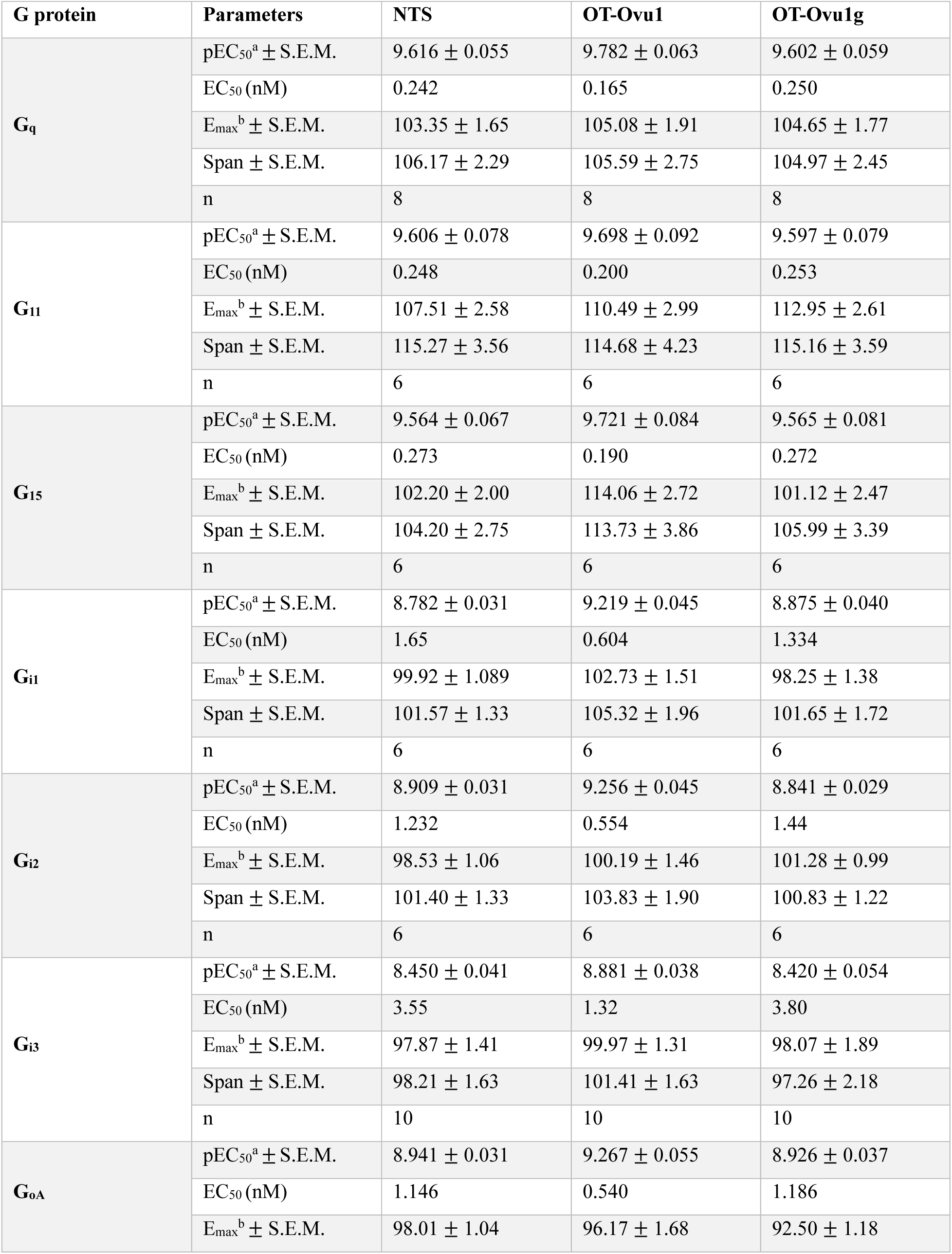
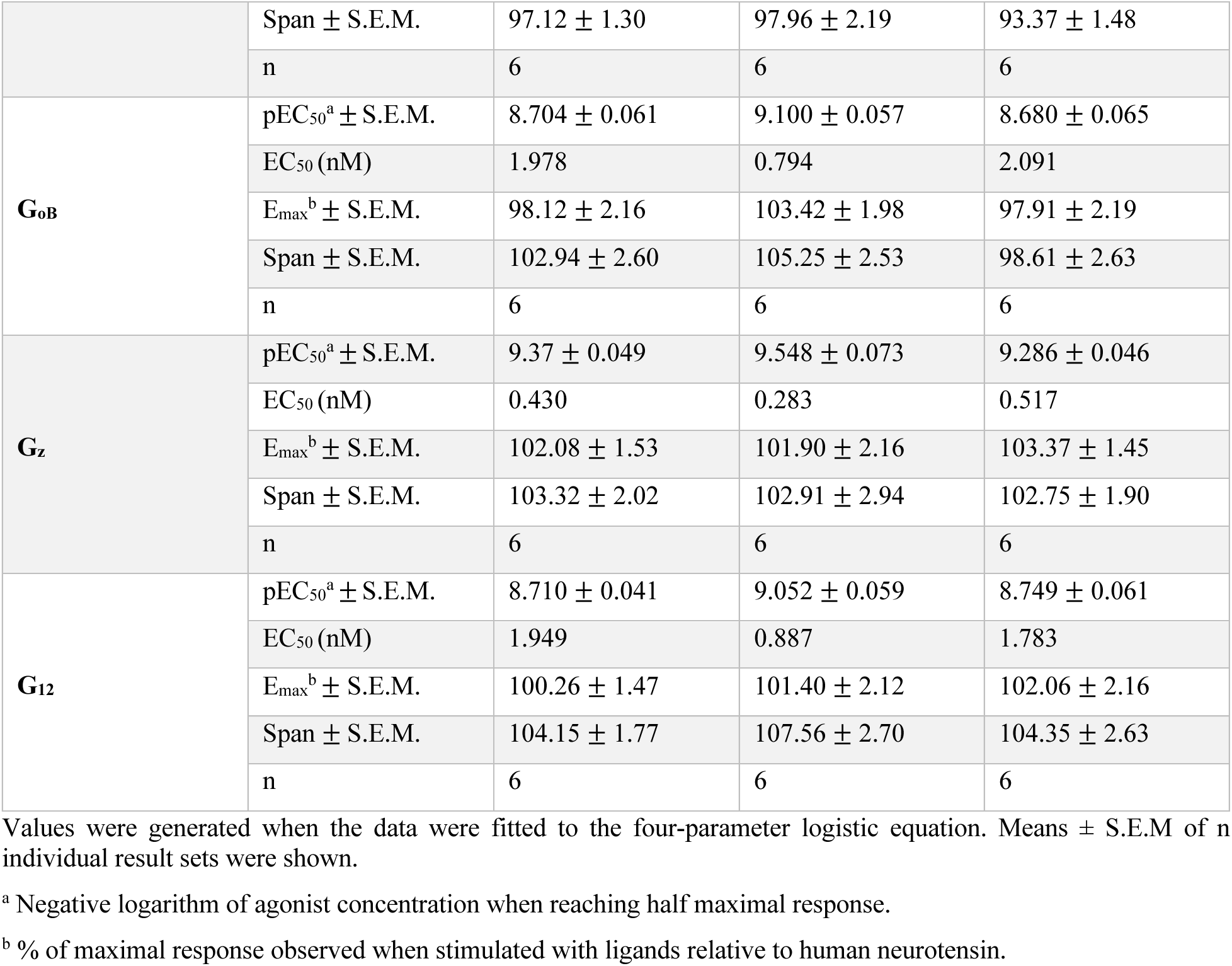
Pharmacological parameters of the G protein dissociation mediated by NTS, OT-Ovu1, OT-Ovu1g at the human NTSR1.

**Table S10.**
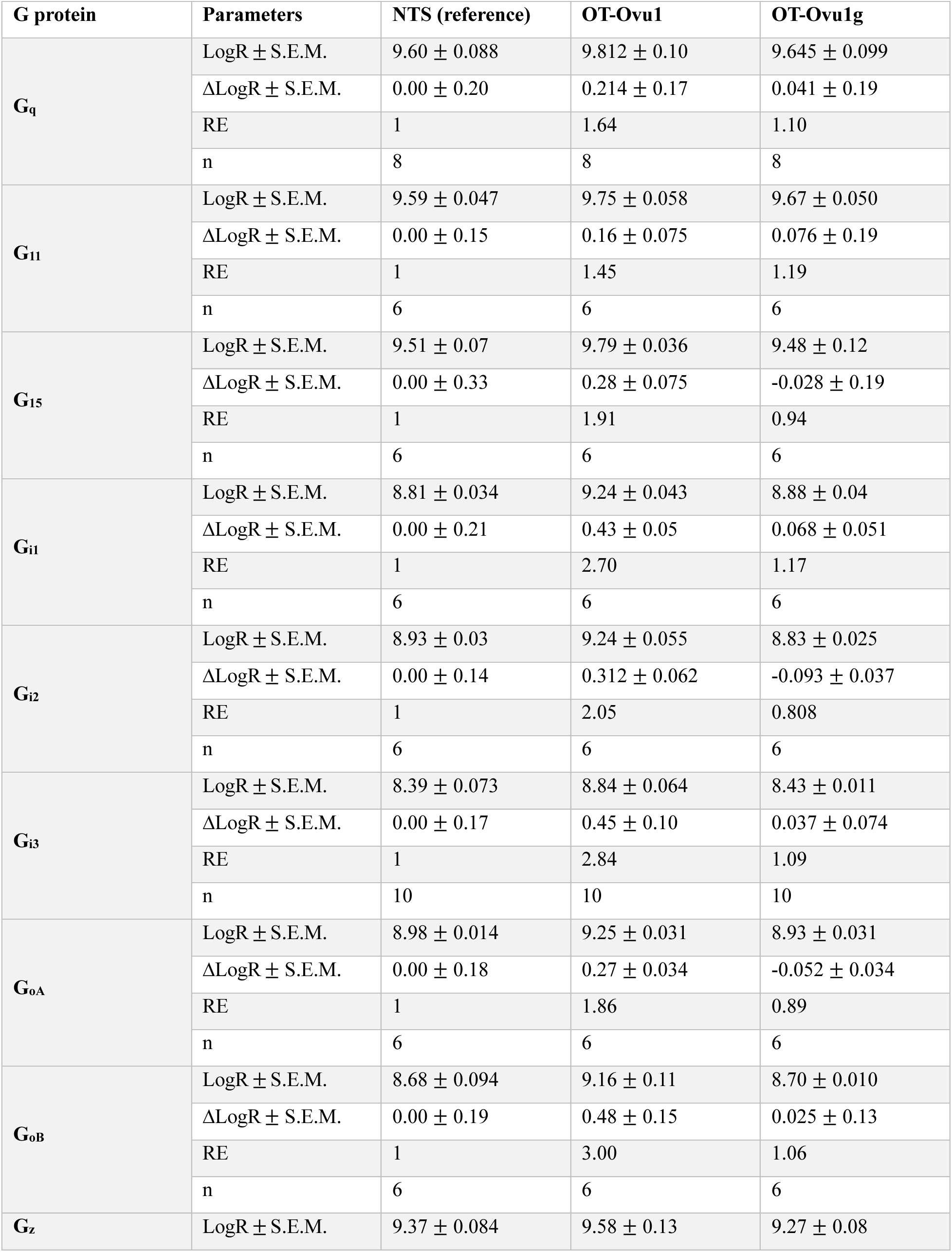
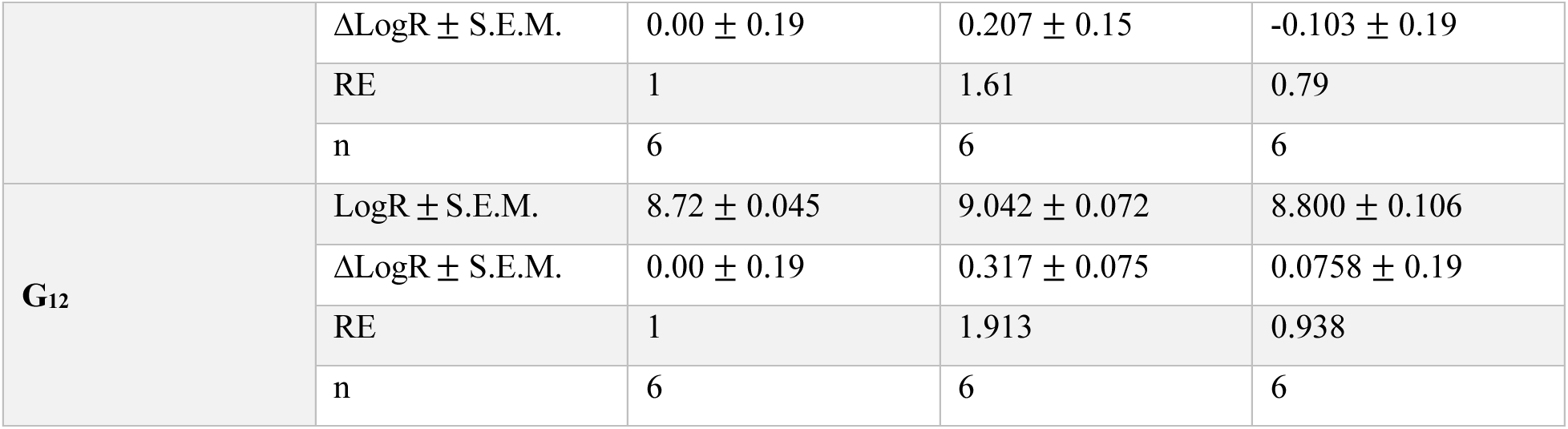
Bias factors of NTS, OT-Ovu1 and OT-Ovu1g mediated G protein dissociation at the human NTSR1.

